# Mitochondrial architecture rearrangements produce asymmetrical nonadaptive mutational pressures that subvert the phylogenetic reconstruction in Isopoda

**DOI:** 10.1101/607960

**Authors:** Dong Zhang, Hong Zou, Cong-Jie Hua, Wen-Xiang Li, Shahid Mahboob, Khalid Abdullah Al-Ghanim, Fahad Al-Misned, Ivan Jakovlić, Gui-Tang Wang

## Abstract

The phylogeny of Isopoda, a speciose order of crustaceans, remains unresolved, with different datasets often producing starkly incongruent phylogenetic hypotheses. We hypothesised that extreme diversity in their life histories might be causing compositional heterogeneity/heterotachy in their mitochondrial genomes, and compromising the phylogenetic reconstruction. We tested the effects of different datasets (mitochondrial, nuclear, nucleotides, amino acids, concatenated genes, individual genes, gene orders), phylogenetic algorithms (assuming data homogeneity, heterogeneity, and heterotachy), and partitioning; and found that almost all of them produced unique topologies. As we also found that mitogenomes of Asellota and two Cymothoida families (Cymothoidae and Corallanidae) possess inversed base (GC) skew patterns in comparison to other isopods, we concluded that inverted skews cause long-branch attraction phylogenetic artefacts between these taxa. These asymmetrical skews are most likely driven by multiple independent inversions of origin of replication (i.e., nonadaptive mutational pressures). Although the PhyloBayes CAT-GTR algorithm managed to attenuate some of these artefacts (and outperform partitioning), mitochondrial data have limited applicability for reconstructing the phylogeny of Isopoda. Regardless of this, our analyses allowed us to propose solutions to some unresolved phylogenetic debates, and support Asellota are the most likely candidate for the basal isopod branch. As our findings show that architectural rearrangements can produce major compositional biases even on short evolutionary timescales, the implications are that proving the suitability of data via composition skew analyses should be a prerequisite for every study that aims to use mitochondrial data for phylogenetic reconstruction, even among closely related taxa.

## Introduction

Significant taxonomic and phylogenetic uncertainty permeates the entire order of Isopoda. The members of this highly speciose (>10,000) order of crustaceans (class Malacostraca) exhibit a remarkable diversity in their life histories (comprising both free-living and parasitic species) and occupy almost all habitats on the planet Earth (marine, freshwater and terrestrial), from deep sea vents to the Antarctica. The traditional morphology-based taxonomic classification and identification of isopods is further (aside from the speciosity) hampered by great intraspecific morphological variation, sexual dimorphism, sequential hermaphroditism, relatively flexible host preference, and global distribution of many species (Joca, Leray, Zigler, & Brusca, 2015; L. S. F. Lins, Ho, Wilson, & Lo, 2012; Luana S.F. Lins, Ho, & Lo, 2017; Rudy, Rendoš, Luptáčik, & Mock, 2018; Shen et al., 2017; Wetzer, 2002; George D.F. Wilson, 2008). However, molecular data also appear to be an unreliable tool for the task, as different datasets (mitochondrial genes, mitochondrial genomes, nuclear genes, combined mitonuclear data) often produce very different topologies (Richard C. Brusca, 1981; Hata et al., 2017; Kilpert, Held, & Podsiadlowski, 2012; Luana S.F. Lins et al., 2017; Martin, Bruce, & Nowak, 2016; Poore & Bruce, 2012; Wetzer, 2002; G D F Wilson, 2009; Yu, An, Li, & Boyko, 2018; Zou et al., 2018). As a result, even the identity of the basal isopod clade (defined as the sister-clade to all other isopod lineages (Krell & Cranston, 2004)) remains debated. Traditionally (morphology and single gene-based studies), Phreatoicidea was regarded as the basal clade (Kilpert et al., 2012; Wetzer, 2002; G D F Wilson, 2009), but some studies resolved Phreatoicidea+Aselotta at the base (Shen et al., 2017; G D F Wilson, 2009; George D.F. Wilson, 1999; Yu et al., 2018), one study found Limnoriidea (Luana S.F. Lins et al., 2017) at the base, whereas a few studies even resolved parasitic Cymothoidae and Corallanidae (suborder Cymothoida) at the base (Hua et al., 2018; L. S. F. Lins et al., 2012; Luana S.F. Lins et al., 2017; G D F Wilson, 2009; Zou et al., 2018). As the Cymothoida was traditionally regarded as the most derived isopod clade (Richard C. Brusca, 1981; Kilpert et al., 2012; Wetzer, 2002; G D F Wilson, 2009), these alternative hypotheses cannot be described as minor topological instability. The monophyly of suborder Cymothoida is generally supported by the morphological data, but rejected by the molecular data (Brandt & Poore, 2003; Hua et al., 2018; Kilpert et al., 2012; Luana S.F. Lins et al., 2017; Shen et al., 2017; G D F Wilson, 2009; Yu et al., 2018; Zou et al., 2018). Among a number of other unresolved phylogenetic issues permeating this order are the monophyly of the suborder Oniscidea (supported by morphology, sometimes rejected by molecular data) and the existence of several ‘rogue’ species/taxa, such as *Ligia oceanica* (Ligiidae), *Eurydice pulchra*, and *Limnoria quadripunctata* (Limnoriidae), whose positions in the isopod clade often vary among studies (Kilpert et al., 2012; Luana S.F. Lins et al., 2017; Schmidt, 2008; Shen et al., 2017; Wetzer, Pérez-Losada, & Bruce, 2013; G D F Wilson, 2009; Yu et al., 2018).

Historically, variation found within gene sequences was typically considered to accumulate under a neutral equilibrium model, so commonly used phylogenetic reconstruction algorithms assume homogeneity in mutational rates. As this paradigm began to change during the last few decades (Wolff, Ladoukakis, Enríquez, & Dowling, 2014), this was accompanied by a growing amount of evidence that compositional heterogeneity can compromise phylogenetic reconstruction in some taxa and that evolutionary models operating under that prerequisite may not be suitable for all phylogenetic studies (Cameron, 2014; Hassanin, 2006; Kolaczkowski & Thornton, 2004; Lartillot, Brinkmann, & Philippe, 2007; Morgan et al., 2013; Phillips, McLenachan, Down, Gibb, & Penny, 2006; Sheffield, Song, Cameron, & Whiting, 2009; Zhong et al., 2011). However, the feud about the most suitable methodological approach to account for this heterogeneity remains unresolved, with most prominent contenders currently being the CAT models (Feuda et al., 2017), and partitioning schemes, i.e. different evolutionary models assigned to different character blocks, assuming homogeneity within each block (Whelan & Halanych, 2017). Although these two approaches account for rate heterogeneity across sites, they still assume that substitution rates for sites are constant across all included lineages. From the evolutionary perspective this is not a likely scenario, as substitution rates are likely to be both site- and lineage-specific (Crotty et al., 2017). Indeed, heterotachy, variations in lineage-specific evolutionary rates over time (Lopez, Casane, & Philippe, 2002), is widespread in eukaryotes (Baele, Raes, Van De Peer, & Vansteelandt, 2006).

Mitochondrial genomes (mitogenomes) generally provide much higher phylogenetic resolution than traditionally used morphological and single gene-based molecular markers (Nie et al., 2018), so mitochondrial phylogenomics is increasingly used to tackle phylogenetic controversies (Bourguignon et al., 2018; Cameron, 2014; Der Sarkissian et al., 2015; Lan et al., 2017; Li et al., 2017). Although the resolution of this approach is still limited by a very small number of available mitogenomes in isopods, overview of published studies shows that they also failed to produce results congruent with other approaches, failed to resolve the rogue taxa issues, and taken as a whole generated more questions than answers (Hua et al., 2018; Kilpert et al., 2012; Luana S.F. Lins et al., 2017; Shen et al., 2017; Yu et al., 2018; Zou et al., 2018).

The evolutionary history of Isopoda abounds in independent (major) life history innovations, such as free-living to parasitic lifestyle (Hata et al., 2017; Jones, Miller, Grutter, & Cribb, 2008; Ketmaier, Joyce, Horton, & Mariani, 2008; Poore & Bruce, 2012), and radical habitat expansions (L. S. F. Lins et al., 2012), such as sea to freshwater, or even water to land (Broly, Deville, & Maillet, 2013; Hata et al., 2017; George D.F. Wilson, 2008). It has been proposed that the Cymothoida may have originated in deep seas, subsequently expanded to shallow seas, and then to brackish and freshwater (likely on several independent occasions) (Hata et al., 2017). Signals of adaptation to high altitude (Hassanin, Ropiquet, Couloux, & Cruaud, 2009; Mishmar et al., 2003; Scott et al., 2011), deep-sea environment (Almeida, Maldonado, Vasconcelos, & Antunes, 2015), and shifts in physiological demands (Botero-Castro et al., 2018; Hassanin, 2006) have been identified in mitogenomes of a range of animals. It is therefore highly likely that radical adaptations to life in different environments, from the anoxic environment of deep sea-inhabiting isopod species (L. S. F. Lins et al., 2012) to terrestrial species, would produce strikingly different evolutionary pressures on genomes of species, and result in disparate evolutionary rates of mitochondrial genes, which are central to energy production via the oxidative phosphorylation (Gawryluk et al., 2016). In agreement with this hypothesis are uneven evolutionary rates (dN/dS) observed among isopod mitogenomes (Shen et al., 2017) and different mutational rates of protein-coding genes (PCGs) encoded on the majority (or plus) strand among different lineages of isopods (Lloyd et al., 2015). Conflicting phylogenetic signals among different mitochondrial regions have been reported in a number of metazoan groups, which indicates that different mitochondrial regions can accumulate substitutions in ways that are difficult to model, which can result in biased estimates of phylogeny (Meiklejohn et al., 2014). There is evidence that this compositional heterogeneity may be comparatively highly pronounced in mitogenomes of some arthropod taxa (Cameron, 2014; Hassanin, 2006; Liu, Li, Jakovlić, & Yuan, 2017). We therefore hypothesised that the aforementioned extreme life history diversity of isopods might cause pronounced compositional heterogeneity/heterotachy in their mitogenomes, and interfere with the reconstruction of the Isopoda phylogeny. Although limitations of mitochondrial (and molecular in general) data for inferring phylogenies have been widely discussed (Ballard & Whitlock, 2004; Edwards, Potter, Schmitt, Bragg, & Moritz, 2016; Grechko, 2013; Hassanin, Léger, & Deutsch, 2005; Rubinoff, Holland, & Savolainen, 2005; Talavera et al., 2011; Willis, 2017), a review of the existing literature reveals that most previous studies of evolutionary history of Isopoda ignored those limitations, or attempted to ameliorate them by using such strategies combined datasets (mtDNA, nuclear DNA, morphology) (Luana S.F. Lins et al., 2017; G D F Wilson, 2009), amino acid sequences (Kilpert et al., 2012; Luana S.F. Lins et al., 2017), or applying different models to each codon position (Hata et al., 2017). However, none of those studies attempted to use algorithms designed specifically to account for compositional heterogeneity/tachy, nor studied this problem directly. To test the hypothesis that compositional heterogeneity interferes with phylogenetic reconstruction in Isopoda, we used a number of different datasets: mitochondrial DNA (single genes, genomes, nucleotides, amino acids, gene orders) and nuclear DNA (*18S*); and methodological approaches: dataset partitioning, maximum likelihood (ML), Bayesian inference (BI), parsimony, PhyloBayes (PB) CAT-GTR model (heterogeneous), and GHOST (heterotachous).

## Materials and Methods

PhyloSuite (Zhang et al., 2018) was used to batch-download all selected mitogenomes from the GenBank, extract genomic features, translate genes into amino acid sequences, semi-automatically re-annotate ambiguously annotated tRNA genes with the help of the ARWEN (Laslett & Canbäck, 2008) output, automatically replace the GenBank taxonomy with the WoRMS database taxonomy, as the latter tends to be more up to date (Costello et al., 2013), generate comparative genome statistics tables, and conduct phylogenetic analyses (Flowchart mode) using a number of incorporated plug-in programs. Nucleotide and amino acid sequences of protein-coding genes (PCGs) were aligned in batches (using codon and normal-alignment modes respectively) with ‘--auto’ strategy, whereas rRNA genes (including the nuclear *18S*) were aligned using Q-INS-i algorithm, which takes secondary structure information into account, all implemented in MAFFT (Katoh & Standley, 2013; Katoh & Toh, 2008). Gblocks (Castresana, 2000; Talavera & Castresana, 2007) was used to remove ambiguously aligned regions from the concatenated alignments with default parameter settings. PhyloSuite was used to concatenate the alignments (all provided in the File S4). Data partitioning schemes were inferred using PartitionFinder2 (Lanfear, Calcott, Ho, & Guindon, 2012), and selection of the most appropriate evolutionary models for each partition was computed according to the Bayesian information criterion scores and weights using ModelFinder (Kalyaanamoorthy, Minh, Wong, Von Haeseler, & Jermiin, 2017). Chi-square (χ^2^) test for the homogeneity of character composition of aligned sequences was performed using IQ-TREE 1.6.8 (Trifinopoulos, Nguyen, von Haeseler, & Minh, 2016). Standard (homogeneous models) phylogenetic analyses were conducted using two programs integrated into PhyloSuite: MrBayes 3.2.6 (Bayesian inference, BI) (Ronquist et al., 2012) and IQ-TREE (Maximum Likelihood, ML). PAUP* 4.0 was used to conduct Parsimony analyses via heuristic searching (TBR branch swapping) and 500 random addition sequence replicates, and bootstrap branch support was calculated via heuristic searches on 1000 pseudo-replicate datasets (Swofford, 2002). The heterogeneous CAT-GTR model is implemented in PhyloBayes-MPI 1.7a (PB) (Lartillot et al., 2007), and the heterotachous GHOST model is implemented in IQ-TREE (Crotty et al., 2017). PhyloBayes was run on the beta version of the Cipres server (https://cushion3.sdsc.edu/portal2/tools.action) (Miller, Pfeiffer, & Schwartz, 2010), with default parameters (burnin = 500, invariable sites automatically removed from the alignment, two MCMC chains), and the analysis was stopped when the conditions considered to indicate a good run (PhyloBayes manual) were reached (maxdiff < 0.1 and minimum effective size > 300). The phylogenetic tree inferred from the gene-order (GO) dataset was reconstructed using MLGO (Hu, Lin, & Tang, 2014), with 1000 bootstrap replicates, and an input file generated by PhyloSuite. Phylograms and gene orders were visualized in iTOL (Letunic & Bork, 2007), and annotated using files generated by PhyloSuite. Skews were calculated and plotted using PhyloSuite and GraphDNA (Thomas, Horspool, Brown, Tcherepanov, & Upton, 2007).

## Results

### Mitochondrial datasets

As a majority of available isopod mitogenomes are incomplete, we were faced with the trade-off between the amount of data used in the analysis and the number of species used: after removing six (some incomplete and duplicates) of the 27 available isopod mitogenomes (Oct. 2018), the dataset comprised 8 complete and 13 partial sequences (Table 1). To attempt to resolve the debated issue of the basal Isopod clade with maximum resolution, we used a relatively large number of outgroups for phylogenetic analyses: a basal arthropod, Limulus polyphemus (Lavrov, Boore, & Brown, 2000), and a number of closely related non-isopod malacostracan taxa: three Decapoda, two Stomatopoda, two Amphipoda, one Mysida and one Euphausiacea species (Kilpert & Podsiadlowski, 2006; G D F Wilson, 2009). We conducted phylogenetic analyses using a number of different datasets: NUC - nucleotides of concatenated 13 PCGs and two rRNA genes (*rrnL* and *rrnS*); AAs – concatenated amino acid sequences of 13 PCGs; 15 single-gene datasets (13 PCGs + 2 rRNAs); gene families (nad+atp and cox+cytb); and gene orders. We also tested the performance of data partitioning, by conducting the same analyses on both non-partitioned and partitioned datasets. The best partitioning scheme divided the NUC dataset by individual genes, with only *cox2*/*cox3*, *atp6*/*nad3*, and *rrnS*/*L* placed together in a same partition. Further details available in Supporting Information (File S1).

**Table 1.**
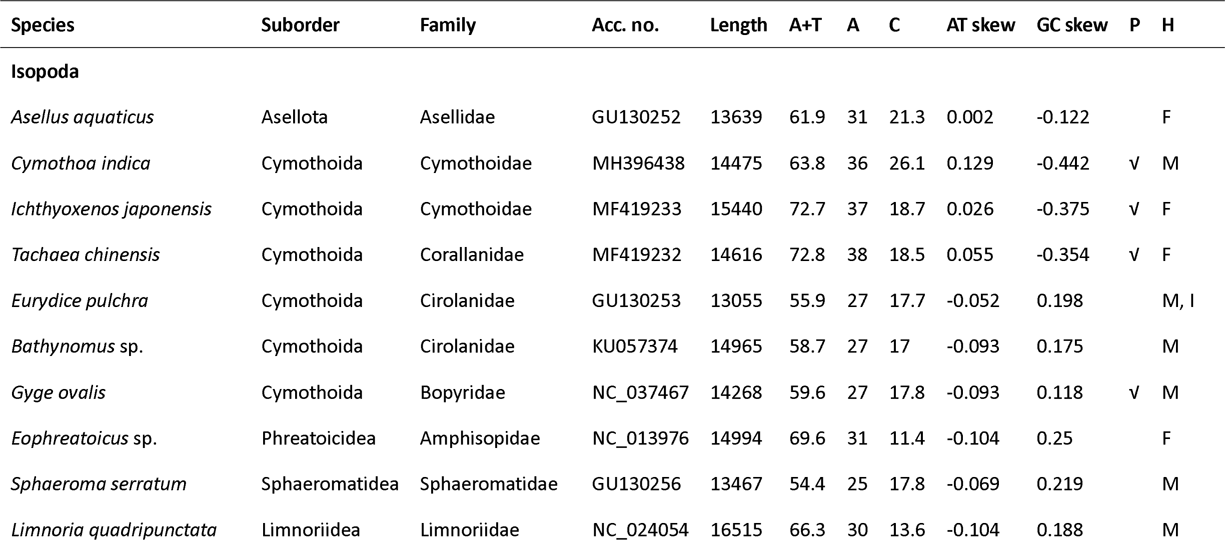
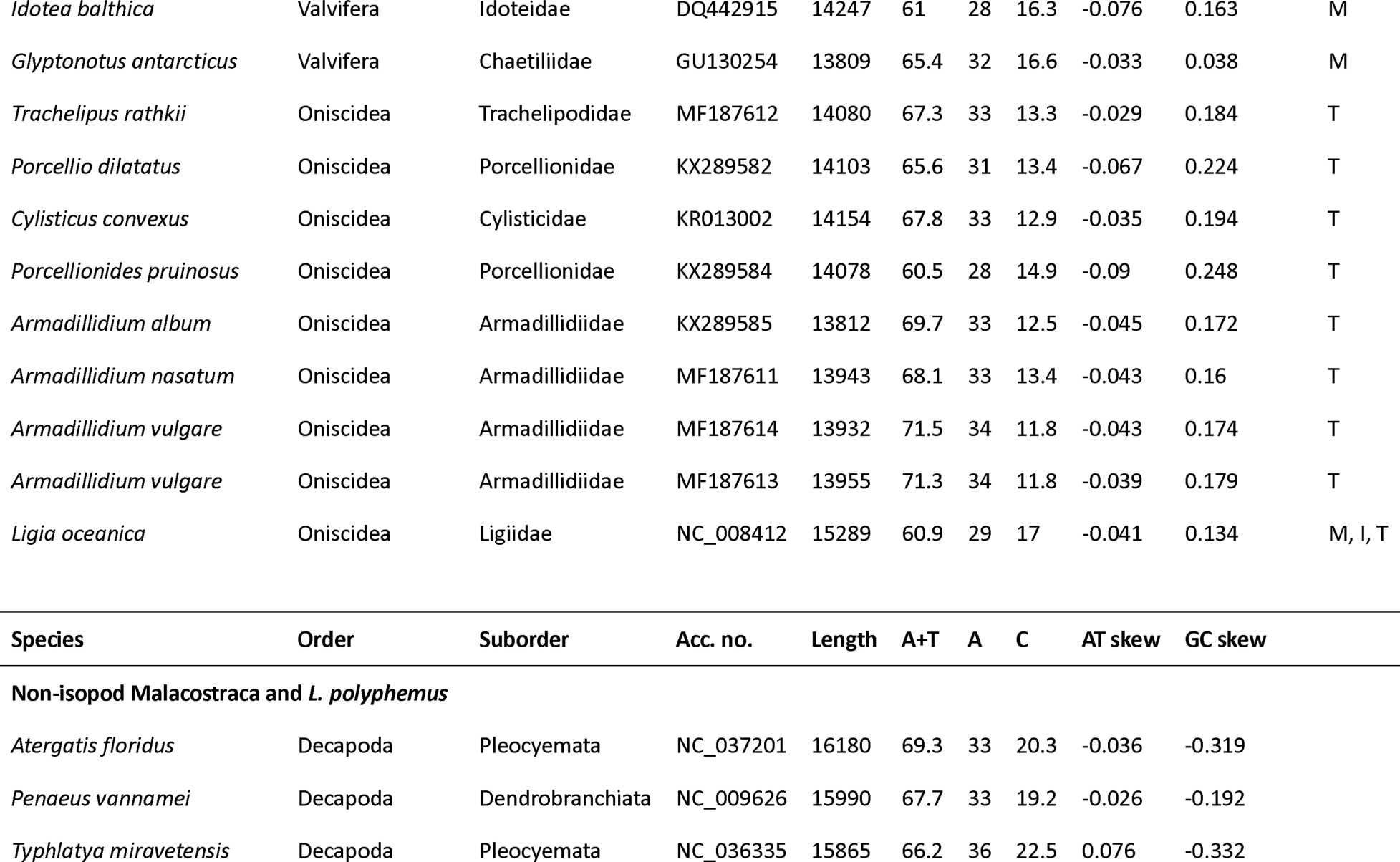
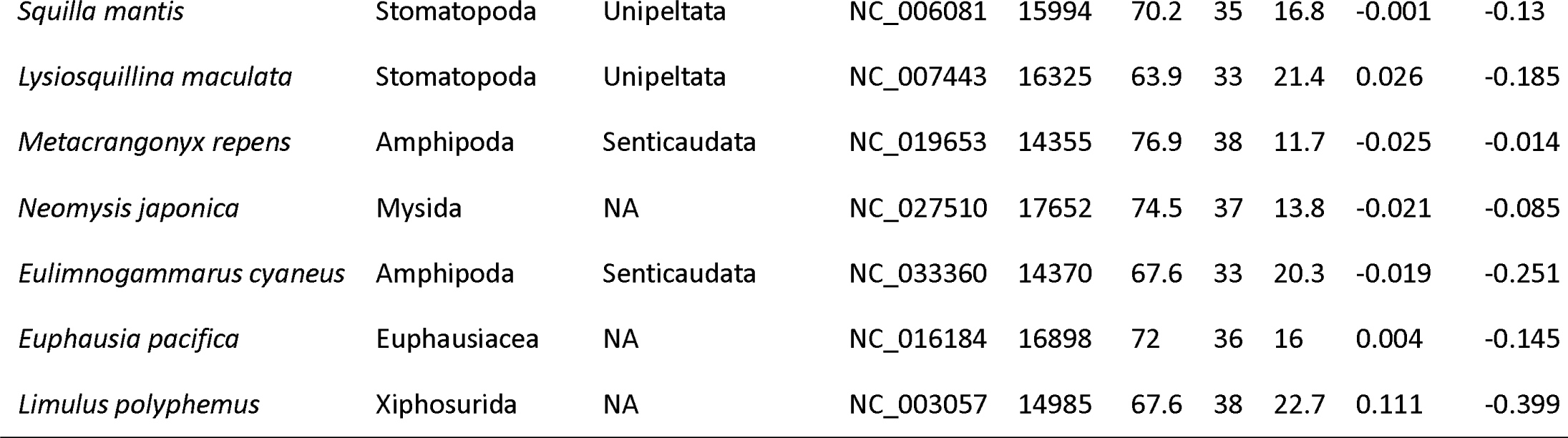
Taxonomy, length (bp), base composition (%) and skews of mitogenomes used in the analysis. P column indicates whether a species is parasitic. H indicates the habitat: F - freshwater, M - marine, T - terrestrial, and I - intertidal.

| Species | Suborder | Family | Acc. no. | Length | A+T | A | C | AT skew | GC skew | P | H |
| --- | --- | --- | --- | --- | --- | --- | --- | --- | --- | --- | --- |
| **Isopoda** |  |  |  |  |  |  |  |  |  |  |  |
| *Asellus aquaticus* | Asellota | Asellidae | GU130252 | 13639 | 61.9 | 31 | 21.3 | 0.002 | -0.122 |  | F |
| *Cymothoa indica* | Cymothoida | Cymothoidae | MH396438 | 14475 | 63.8 | 36 | 26.1 | 0.129 | -0.442 | √ | M |
| *Ichthyoxenos japonensis* | Cymothoida | Cymothoidae | MF419233 | 15440 | 72.7 | 37 | 18.7 | 0.026 | -0.375 | √ | F |
| *Tachaea chinensis* | Cymothoida | Corallanidae | MF419232 | 14616 | 72.8 | 38 | 18.5 | 0.055 | -0.354 | √ | F |
| *Eurydice pulchra* | Cymothoida | Cirolanidae | GU130253 | 13055 | 55.9 | 27 | 17.7 | -0.052 | 0.198 |  | M, I |
| *Bathynomus* sp. | Cymothoida | Cirolanidae | KU057374 | 14965 | 58.7 | 27 | 17 | -0.093 | 0.175 |  | M |
| *Gyge ovalis* | Cymothoida | Bopyridae | NC_037467 | 14268 | 59.6 | 27 | 17.8 | -0.093 | 0.118 | √ | M |
| *Eophreaticoicus* sp. | Phreatoicoidea | Amphisopidae | NC_013976 | 14994 | 69.6 | 31 | 11.4 | -0.104 | 0.25 |  | F |
| *Sphaeroma serratum* | Sphaeromatidea | Sphaeromatidae | GU130256 | 13467 | 54.4 | 25 | 17.8 | -0.069 | 0.219 |  | M |
| *Limnoria quadripunctata* | Limnoriidea | Limnoriidae | NC_024054 | 16515 | 66.3 | 30 | 13.6 | -0.104 | 0.188 |  | M |

|  |  |  |  |  |  |  |  |  |  |  |
| --- | --- | --- | --- | --- | --- | --- | --- | --- | --- | --- |
| *Idotea balthica* | Valvifera | Idoteidae | DQ442915 | 14247 | 61 | 28 | 16.3 | -0.076 | 0.163 | M |
| *Glyptonotus antarcticus* | Valvifera | Chaetiliidae | GU130254 | 13809 | 65.4 | 32 | 16.6 | -0.033 | 0.038 | M |
| *Trachelipus rathkii* | Oniscidea | Trachelipodidae | MF187612 | 14080 | 67.3 | 33 | 13.3 | -0.029 | 0.184 | T |
| *Porcellio dilatatus* | Oniscidea | Porcellionidae | KX289582 | 14103 | 65.6 | 31 | 13.4 | -0.067 | 0.224 | T |
| *Cylisticus convexus* | Oniscidea | Cylisticidae | KR013002 | 14154 | 67.8 | 33 | 12.9 | -0.035 | 0.194 | T |
| *Porcellionides pruinosus* | Oniscidea | Porcellionidae | KX289584 | 14078 | 60.5 | 28 | 14.9 | -0.09 | 0.248 | T |
| *Armadillidium album* | Oniscidea | Armadillidiidae | KX289585 | 13812 | 69.7 | 33 | 12.5 | -0.045 | 0.172 | T |
| *Armadillidium nasatum* | Oniscidea | Armadillidiidae | MF187611 | 13943 | 68.1 | 33 | 13.4 | -0.043 | 0.16 | T |
| *Armadillidium vulgare* | Oniscidea | Armadillidiidae | MF187614 | 13932 | 71.5 | 34 | 11.8 | -0.043 | 0.174 | T |
| *Armadillidium vulgare* | Oniscidea | Armadillidiidae | MF187613 | 13955 | 71.3 | 34 | 11.8 | -0.039 | 0.179 | T |
| *Ligia oceanica* | Oniscidea | Ligiidae | NC_008412 | 15289 | 60.9 | 29 | 17 | -0.041 | 0.134 | M, I, T |

| Species | Order | Suborder | Acc. no. | Length | A+T | A | C | AT skew | GC skew |
| --- | --- | --- | --- | --- | --- | --- | --- | --- | --- |
| Non-isopod Malacostraca and *L. polyphemus* |  |  |  |  |  |  |  |  |  |
| *Atergatis floridus* | Decapoda | Pleocyemata | NC_037201 | 16180 | 69.3 | 33 | 20.3 | -0.036 | -0.319 |
| *Penaeus vannamei* | Decapoda | Dendrobranchiata | NC_009626 | 15990 | 67.7 | 33 | 19.2 | -0.026 | -0.192 |
| *Typhlatya miravetensis* | Decapoda | Pleocyemata | NC_036335 | 15865 | 66.2 | 36 | 22.5 | 0.076 | -0.332 |
| *Squilla mantis* | Stomatopoda | Unipeltata | NC_006081 | 15994 | 70.2 | 35 | 16.8 | -0.001 | -0.13 |
| *Lysiosquillina maculata* | Stomatopoda | Unipeltata | NC_007443 | 16325 | 63.9 | 33 | 21.4 | 0.026 | -0.185 |
| *Metacrangonyx repens* | Amphipoda | Senticaudata | NC_019653 | 14355 | 76.9 | 38 | 11.7 | -0.025 | -0.014 |
| *Neomysis japonica* | Mysida | NA | NC_027510 | 17652 | 74.5 | 37 | 13.8 | -0.021 | -0.085 |
| *Eulimnogammarus cyaneus* | Amphipoda | Senticaudata | NC_033360 | 14370 | 67.6 | 33 | 20.3 | -0.019 | -0.251 |
| *Euphausia pacifica* | Euphausiacea | NA | NC_016184 | 16898 | 72 | 36 | 16 | 0.004 | -0.145 |
| *Limulus polyphemus* | Xiphosurida | NA | NC_003057 | 14985 | 67.6 | 38 | 22.7 | 0.111 | -0.399 |

### Compositional heterogeneity tests

Best-fit model for the non-partitioned NUC dataset was GTR+F+R6. In the partitioned dataset, three *nad* family genes (*nad1*+*2*+*4*) and *atp8* were assigned the TVM+I+G model, and *nad6* the HKY+I+G model, whereas all nine remaining genes were assigned the GTR+I+G model (File S1). In the χ^2^ compositional homogeneity test of the non-partitioned NUC dataset, where a sequence was denoted ‘failed’ if its nucleotide composition significantly deviated from the average composition of the alignment, only an outgroup species *Penaeus vannamei* passed the test (30 sequences failed). However, only 11 species failed the test in the AAs dataset, indicating that the use of amino acids can attenuate compositional heterogeneity.

### Methodological approaches to infer phylogeny

We tested the performance of two phylogenetic methods relying on standard homogenous models, maximum likelihood (ML) and Bayesian inference (BI), on non-partitioned and partitioned data, and non-standard heterogeneous (CAT-GTR) and heterotachous (GHOST) models on our datasets. The latter two models require the input data to be non-partitioned. The CAT-GTR site mixture model implemented in Phylobayes-MPI 1.7a (Lartillot et al., 2007) allows for site-specific rates of mutation, which is considered to be a more realistic model of amino acid evolution, especially for large multi-gene alignments (Maddock et al., 2016). GHOST model is an edge-unlinked mixture model consisting of several site classes with separate sets of model parameters and edge lengths on the same tree topology, thus naturally accounting for heterotachous evolution (Crotty et al., 2017).

### NUC dataset

GHOST, BI and ML analyses (both partitioned and non-partitioned) of the NUC dataset produced highly congruent topologies (referred to as the ‘NUC-consensus’ topology henceforth). Statistical support values were very high in BI (Fig. 1; all inferred topologies available in the Supporting information: File S2), relatively high in GHOST, and intermediate in ML analyses (low to high). Decapoda were rendered paraphyletic by the Euphausiacea nested within the clade (not in the ML-partitioned tree), and Mysida+Amphipoda were resolved as the sister-clade to Isopoda. In the isopod clade, Cymothoidae+Corallanidae families (Cymothoida) formed the basal clade, followed by Asellota and Phreatoicidea branches. The remaining isopods were divided into two sister-clades: Oniscidea and a ‘catch-all’ clade comprising Limnoriidea, Valvifera, Sphaeromatidea, *L. oceanica* (a ‘rogue’ Oniscidea species), and the remaining three Cymothoida species (*Gyge ovalis*, *Bathynomus* sp. and *Eurydice pulchra*), which did not cluster together (Fig. 1). Parsimony analysis produced a slightly rearranged topology, but crucial features were identical: Mysida+Amphipoda were the sister-clade to Isopoda, and Cymothoidae+Corallanidae were basal isopods. Notable differences were: Asellota forming a sister-clade with G. ovalis + *L. quadripunctata*, and Phreatoicidea at the base of the catch-all clade. The PB analysis produced a notably different overall topology, with all non-isopod lineages forming a sister-clade to the isopods (monophyletic Decapoda), but the topology of the isopod clade was relatively similar to the NUC-consensus, apart from the rogue *Gyge ovalis* (Fig. 2).

**Figure 1.**
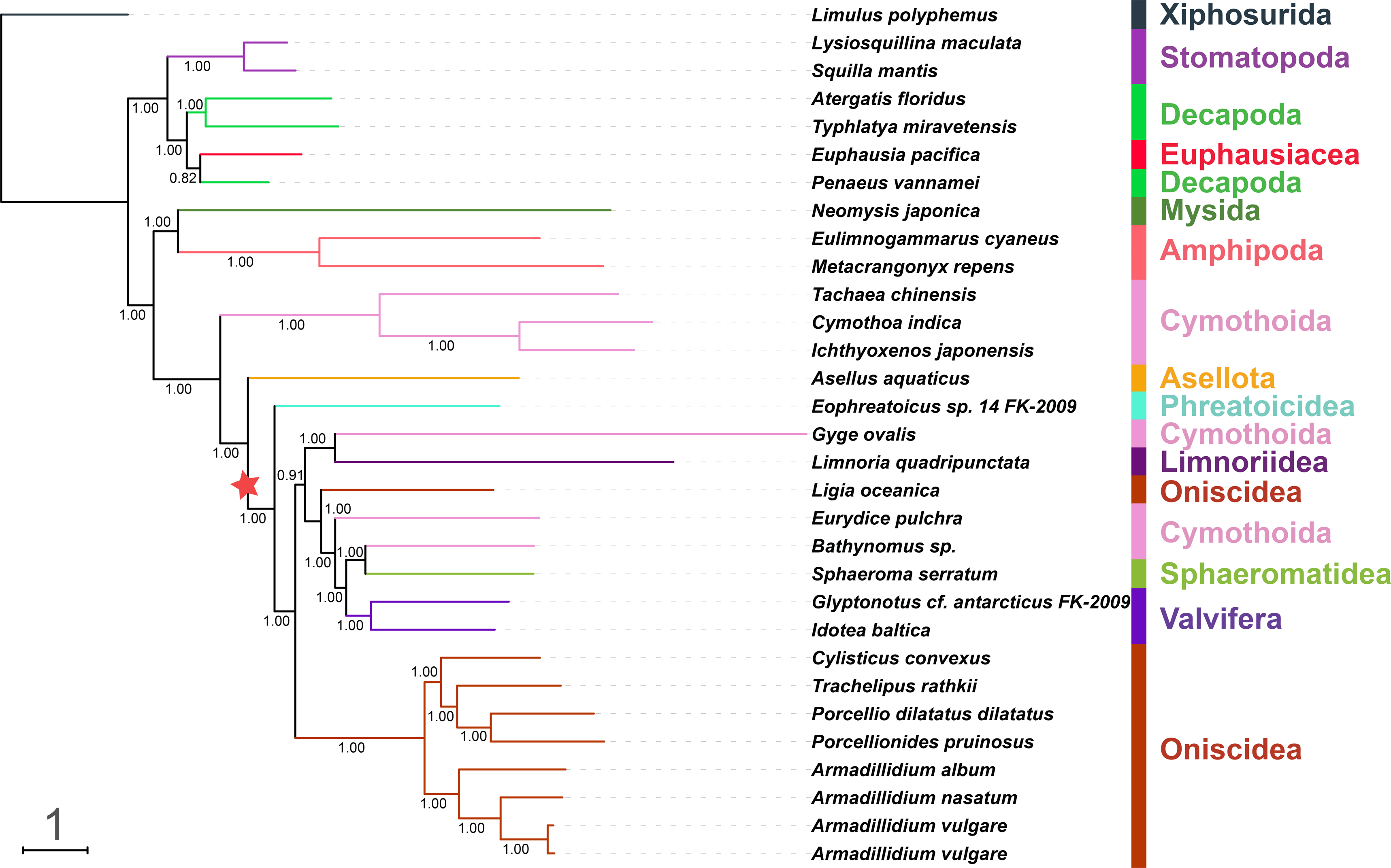
Mitochondrial phylogenomics of Isopoda (suborder information shown) reconstructed using partitioned nucleotide sequences of PCGs and rRNAs (NUC dataset) and BI algorithm. A set of nine non-isopod Malacostraca species and *Limulus polyphemus* were used as outgroups (order information shown). The scale bar corresponds to the estimated number of substitutions per site. Bayesian posterior support values are shown next to corresponding nodes. Star sign indicates a putative origin of replication inversion scenario implied by the topology (see Discussion).

**Figure 2.**
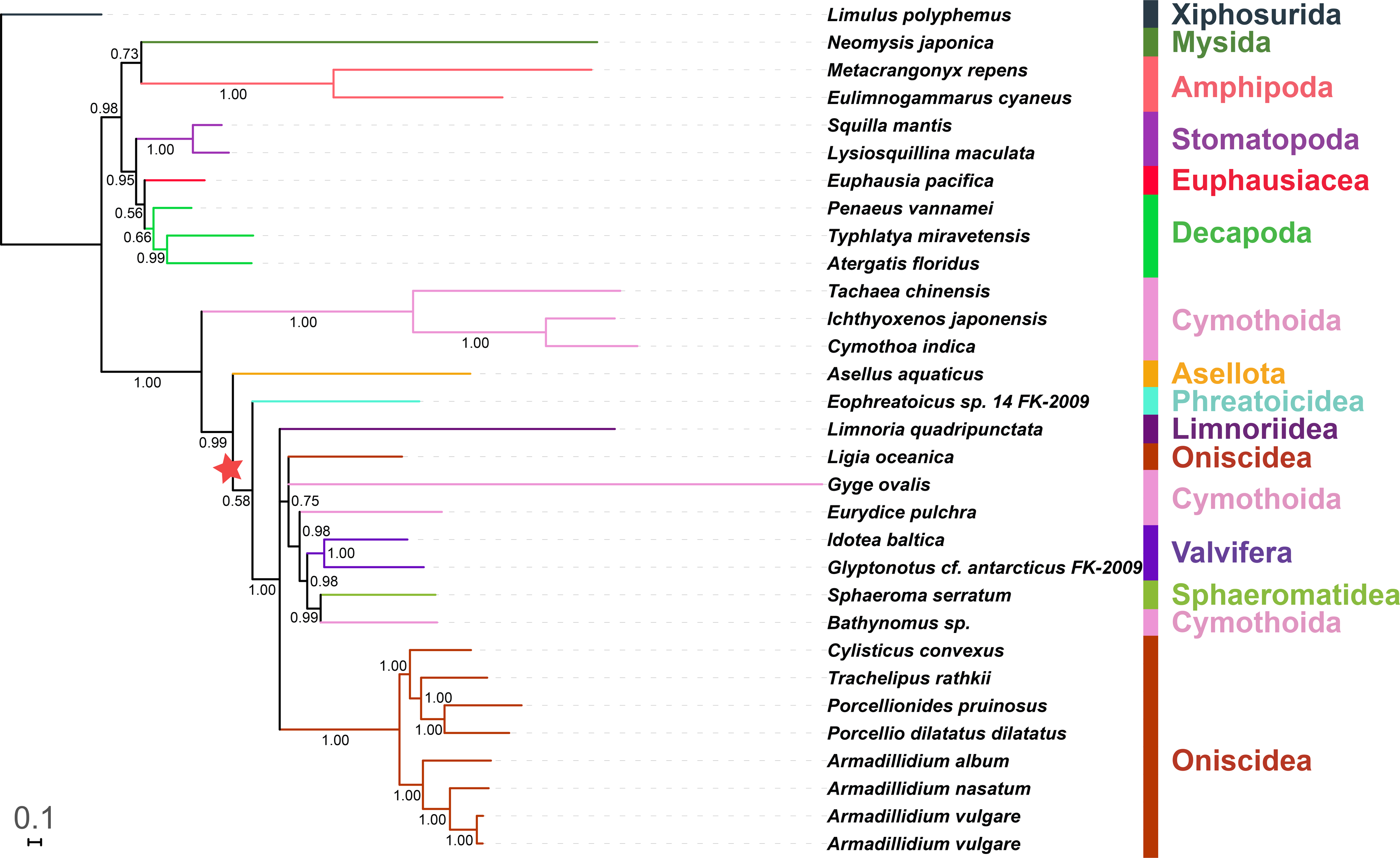
A phylogram reconstructed using nonpartitioned NUC dataset and an algorithm designed to address compositional heterogeneity: CAT-GTR (PB). Posterior Bayesian support values are shown. See Figure 1 for other details.

### Single-gene and concatenated gene family datasets

As we hypothesised that the topological instability may be driven by conflicting signals produced by different genes (Meiklejohn et al., 2014), we conducted ML phylogenetic analyses on 15 single-gene datasets (13 PCGs + 2 rRNAs). Surprisingly, all these produced unique topologies (File S2). *Atp6* resolved Asellota+Cymothoidae+Corallanidae as the basal isopod clade; *atp8* (a very small gene) produced an almost nonsensical topology (defined as: in stark disagreement with any reasonable phylogenetic hypothesis); *nad1* resolved Asellota at the base; *nad2* produced a slightly rearranged NUC-consensus topology; *nad3* (also small, ≈350bp) a nonsensical topology; *nad4* produced a rearranged topology (compared to NUC-consensus), but Cymothoidae+Corralanidae were the basal isopod clade; *nad4L* (small gene) produced a highly rearranged isopod clade; despite its large size (≈1700bp); *nad5* produced a non-canonical isopod topology; *nad6* produced a nonsensical topology (paraphyletic Isopoda, rogue *G. ovalis*); *cox1* produced a highly rearranged topology, including both the non-isopod malacostraca and isopod clades, with Phreatoicidea+Cymothoidae+Corralanidae as the basal isopod clade, and non-canonical Valvifera position; *cox2* produced a slightly rearranged NUC-consensus topology; *cox3* produced a highly rearranged topology, with a unique Asellota+Corallanidae clade at the isopod base and *L. quadripunctata* nested within the Oniscidea; *cytb* produced basal Phreatoicidea, and the remaining taxa divided into Oniscidea and ‘catch all’ sister-clades, where the rogue Cymothoidae+Corallanidae (along with Asellota) were on a long branch in the derived part of the clade; *rrnS* (or *12S*) topology was in some aspects similar to *cytb*, but with Phreatoicidea+Limnoriidea as the basal isopod clade; *rrnL* (or *16S*) produced an almost nonsensical topology, with paraphyletic Isopoda and rogue *G. ovalis*.

As some of the previous studies (see Introduction) used datasets combined of several genes, and as some of our optimal partitioning strategy analyses indicated that different gene families might evolve at somewhat congruent rates, we divided the PCG dataset into two concatenated gene families: nad_atp (*nad1-6* and *atp6-8*) and cox_cytb (*cox1-3* and *cytb*). ML analyses of both datasets also produced unique topologies. In comparison to NUC-consensus tree, nad_atp dataset resolved Mysida as the sister-group to all other Malacostraca, and rearranged Phreatoicidea, Limnoriidea and *G. ovalis*. Cox_cytb dataset produced somewhat rearranged non-isopod Malacostraca, a minor discrepancy in the position of *E. pulchra*, and slightly rearranged Oniscidea. Notably, none of the single-gene topologies corresponded to the two family topologies.

### AAs dataset: amino acids of 13 PCGs

Amino acids produced topologies that differed from the NUC-consensus one, with instable topology of the non-isopod Malacostraca, including paraphyletic Eucarida (Decapoda+Euphausiacea) in five (BI/ML, non-partitioned/partitioned, and GHOST; Fig. 3) out of six (PB; Fig. 4) analyses. Sister-clade to Isopoda also varied: Mysida+Amphipoda in Parsimony and both BI analyses, and Mysida in the remaining four analyses. In the isopod clade, partitioning had a major effect on the BI analysis: non-partitioned dataset resolved Cymothoidae+Corallanidae at the base, followed by Phreatoicidea+Asellota. Parsimony analysis produced a similar topology, but Phreatoicidea and *G. ovalis* + *L. quadripunctata* clades switched places. The partitioned BI dataset, as well the remaining five analyses, all resolved Phreatoicidea as the basal clade. In these, the sister-group to the remaining isopods (minus Phreatoicidea) was Asellota+Cymothoidae+Corallanidae in four analyses, and Asellota in PB. Five analyses (minus PB) produced a topology of the remainder of the isopod clade that was partially congruent with the NUC-consensus topology, but instead of it being divided into Oniscidea + all other taxa, here *G. ovalis* + *L. quadripunctata* were at the base (except in Parsimony: Phreatoicidea). Oniscidea (rendered paraphyletic by *L. oceanica*) clade topology was stable in all six, but the topology of the catch-all clade exhibited a number of permutations, none of which were identical to the NUC-consensus topology.

**Figure 3.**
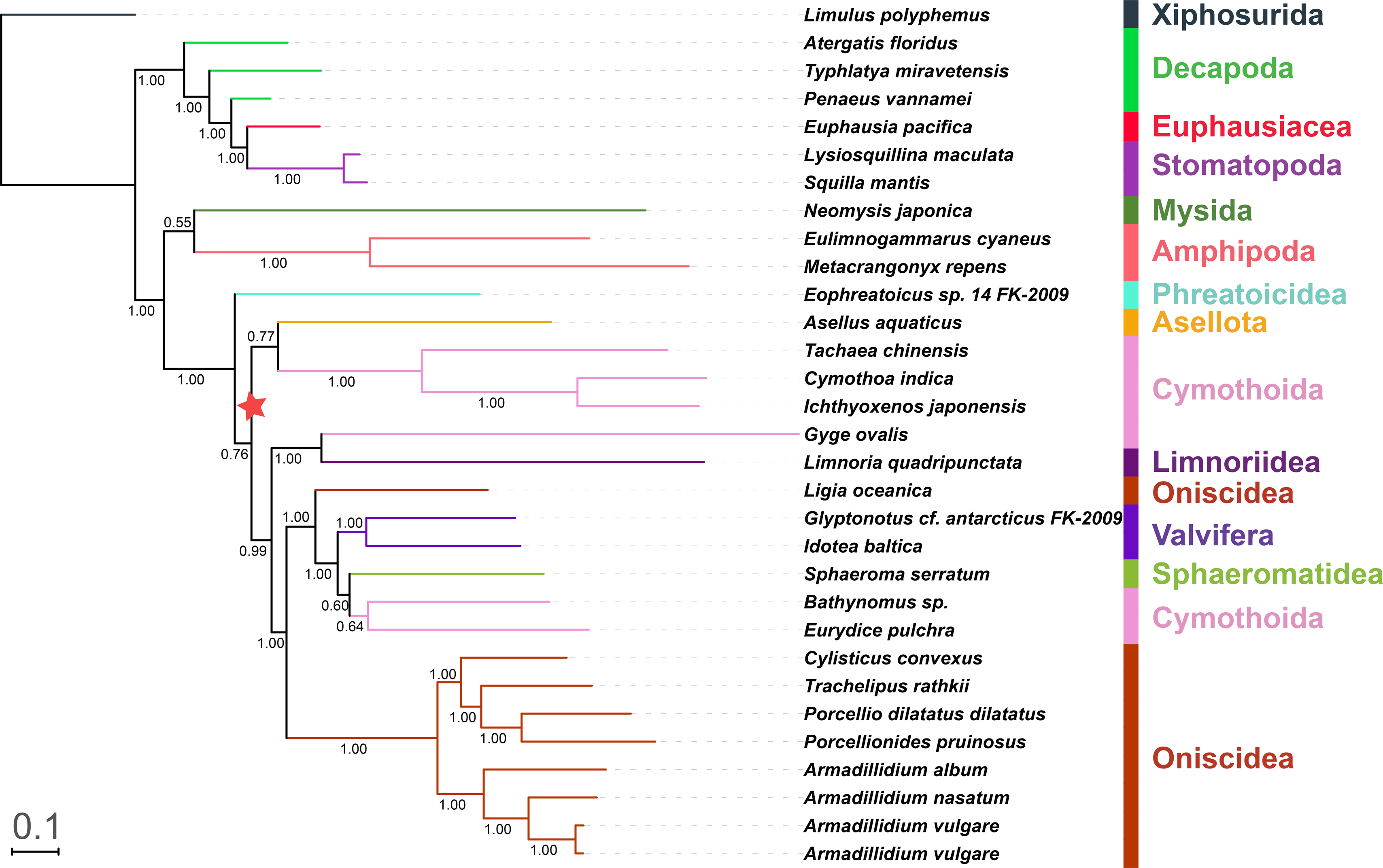
A phylogram reconstructed using amino acid dataset (AAs; 13 PCGs) in combination with data partitioning strategy and BI algorithm. See Figure 1 for other details.

**Figure 4.**
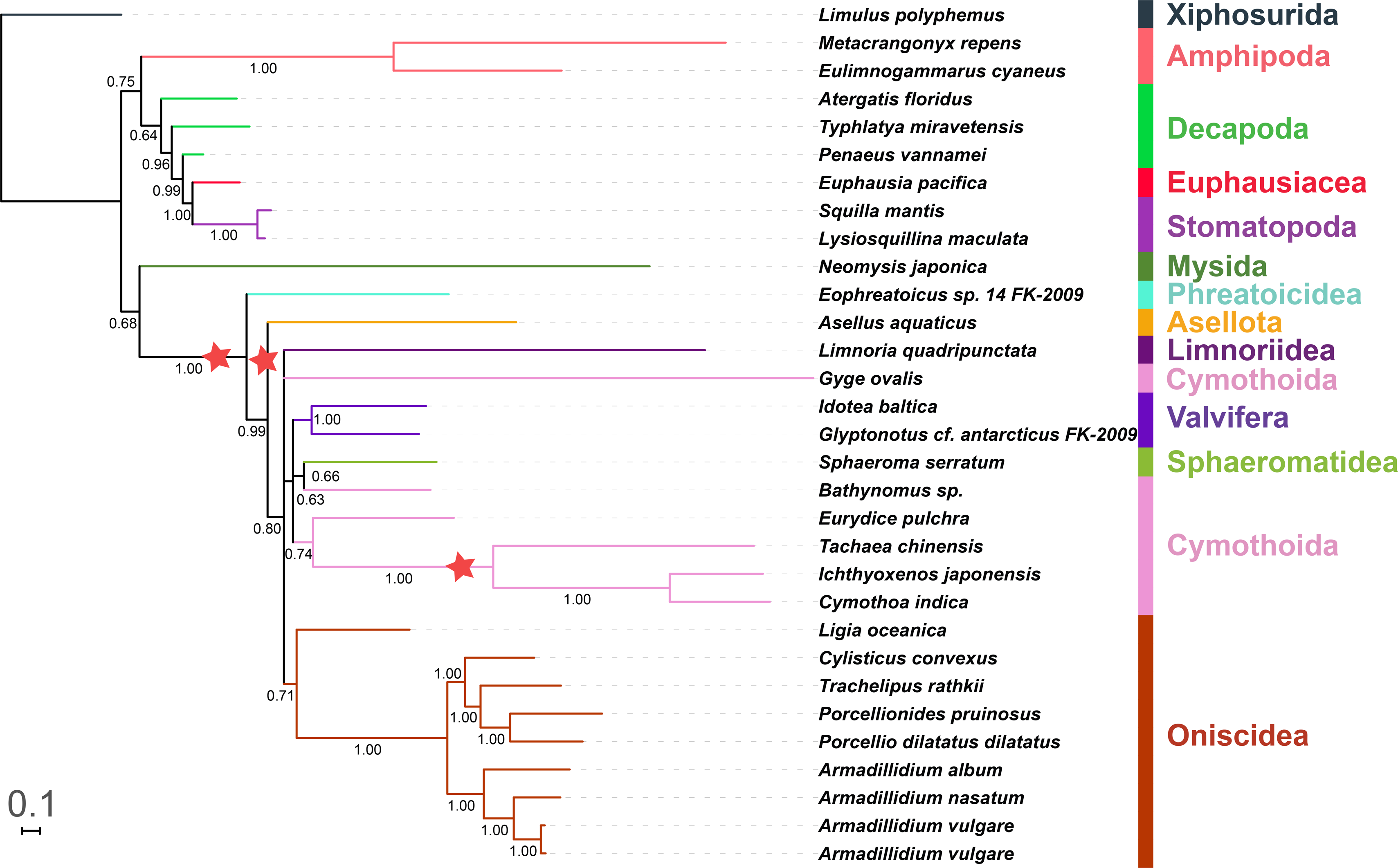
A phylogram reconstructed using AAs dataset and CAT-GTR algorithm designed for heterogeneous datasets (PB). See Figure 1 for other details.

PB produced the only topology (Fig. 4) with monophyletic Oniscidea, with the rogue *L. oceanica* at the base of the clade. Also importantly, Cymothoidae+Corallanidae clade was placed on a long branch within the catch-all clade, together with *Eurydice pulchra* (all Cymothoida). The suborder was still rendered paraphyletic by the positions of *Bathynomus* sp. and rogue (polytomy) *G. ovalis*.

### Gene order

As gene orders are unlikely to exhibit homoplasy, it has been hypothesised that they may be able to resolve difficult (deep) phylogenies in some cases (Boore, 2006), so we tested this approach. As many tRNA genes were missing or we suspected that they may be misannotated, to test for the presence of false signals, we used two datasets: one excluding all tRNAs (PCGs+rRNAs) and one (PCGs+rRNAs+tRNAs) excluding only the missing and ambiguously annotated tRNAs (such as tRNAL1 and 2). The two datasets produced incongruent topologies; different runs of the PCGs+rRNAs+tRNAs dataset produced identical topologies (Fig. 5), whereas different runs of PCGs+rRNAs did not (File S2: GO_PCGs+rRNAs_1 and 2). Both dataset produced a number of paraphyletic major clades and largely nonsensical topologies and largely low support values; for example, the more stable, PCGs+rRNAs+tRNAs, dataset produced paraphyletic Decapoda, Stomatopoda, Isopoda, Oniscidea and Cymothoida.

**Figure 5.**
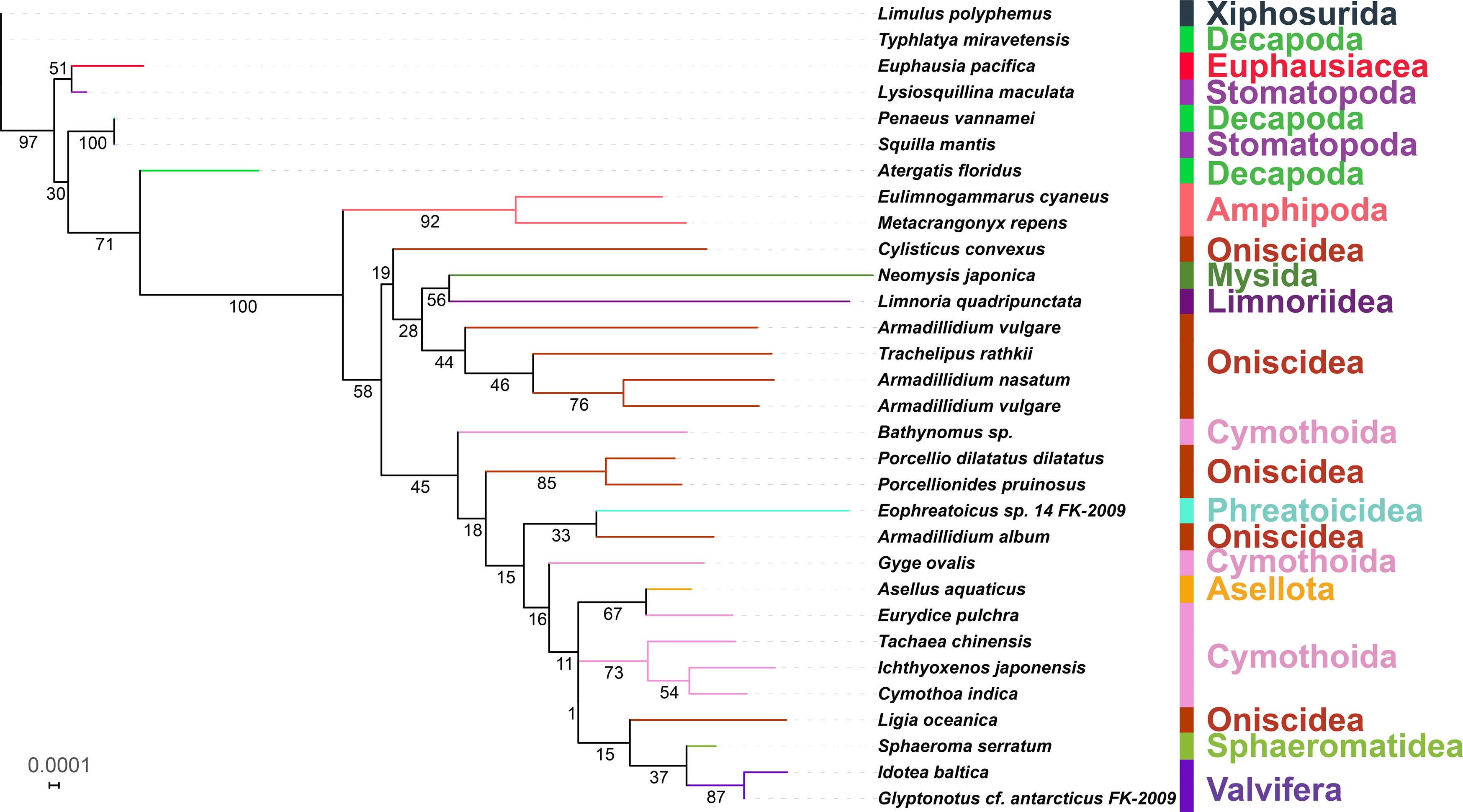
A phylogram reconstructed using mitochondrial gene orders (PCGs+rRNAs+tRNAs). Bootstrap support values are shown next to corresponding nodes. See Figure 1 for other details.

### Nuclear marker-based phylogeny (*18S*)

To view the evolutionary history of Isopoda from a non-mitochondrial perspective, we used the nuclear *18S* gene. This approach can also help us test whether the underlying reason for the conflicting phylogenetic signals between different studies might be mitochondrial introgression, which, recent evidence shows, is more widespread than previously thought (Edwards et al., 2016; Jakovlić, Wu, Treer, Šprem, & Gui, 2013; Mallet, Besansky, & Hahn, 2016). Some of the species used for mitogenomic analyses were not available, but we made sure to include representatives of all major taxa in the mitogenomic dataset (File S3). To obtain a more comparable dataset, we sequenced the *18S* gene of three parasitic Cymothoidae/Corallanidae species from the mtDNA dataset: *Cymothoa indica* (Cymothoidae; GenBank accession number MK079664), *Ichthyoxenos japonensis* (Cymothoidae; MK542857), and *Tachaea chinensis* (Corallanidae; MK542858); and an additional Cymothoidae species, *Asotana magnifica* (MK542856). We conducted a phylogenetic analysis using 55 malacostracan orthologues, with SYM+R4 selected as the best-fit model. Only one sequence (non-isopod: *Gammarus troglophilus*) failed the χ^2^ compositional homogeneity test. Results produced by this dataset were unstable, i.e. different runs and datasets (addition and removal of some taxa) would usually produce different topologies. However, some important features were rather constant among most analyses: Asellota at the base (Limnoriidea in Parsimony, but with very low support); highly derived Cymothoida, rendered paraphyletic by the nested Limnoriidea (monophyletic in Parsimony); and Oniscidea rendered paraphyletic by the rogue Ligiidae clade (*Ligia* sp. and *Ligidium* sp.). ML and BI analyses produced relatively congruent topologies, with Isopoda rendered paraphyletic by the Amphipoda clade nested within the large Cymothoida clade (File S2). As PB and Parsimony analyses produced monophyletic Isopoda, with Amphipoda as the sister-clade (Fig. 6), we can confidently reject this as a compositional heterogeneity artefact. Phreatoicidea largely clustered with the rogue Ligiidae clade, but this was not supported by the PB analysis. Cymothoida were monophyletic only in the Parsimony analysis, and divided into two clades in all topologies: a stable monophyletic clade comprising Dajidae and Bopyridae; and a large instable (exhibiting pervasive paraphyly) clade comprising Cirolanidae, Corallanidae and Cymothoidae families, and aforementioned intruders (Limnoriidea and Amphipoda). Corallanidae and Cirolanidae were mostly paraphyletic (somewhat erratic behaviour of *Eurydice pulchra*), whereas Cymothoidae (monophyletic) were highly derived and exhibited disproportionately long branches.

**Figure 6.**
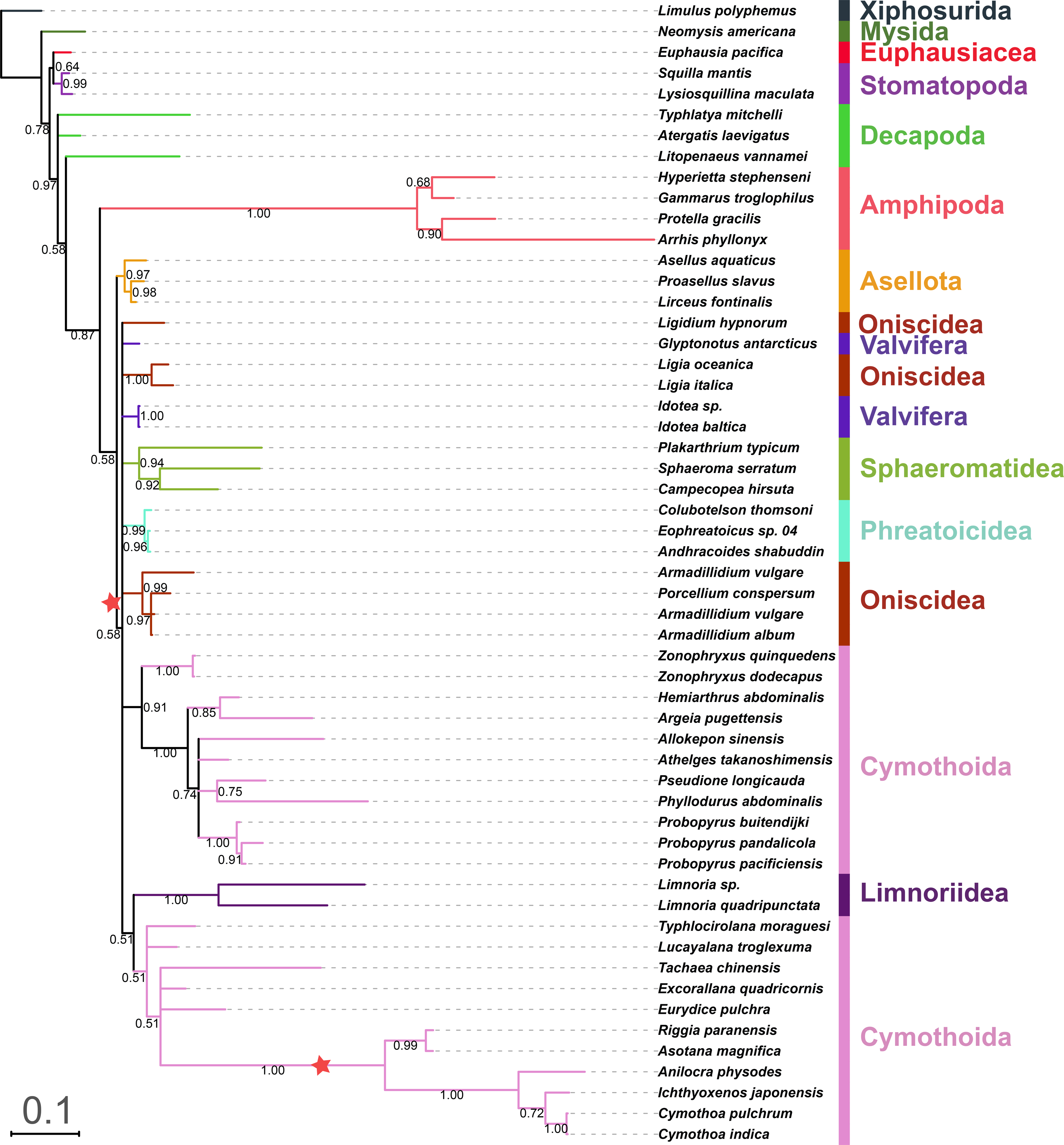
A phylogram inferred using the nuclear *18S* gene and CAT-GTR algorithm (PB). See Figure 1 for other details.

## Discussion

Most phylogenetic reconstruction algorithms assume homogeneity in nucleotide base compositions, but mitochondrial genomes of some groups of Arthropoda exhibit non-constant equilibrium nucleotide frequencies across different lineages, or compositional heterogeneity, which can produce artificial clustering in phylogenetic analysis (Hassanin, 2006; Lartillot et al., 2007; Morgan et al., 2013; Talavera et al., 2011). Having observed pervasive topological instability among previous attempts to resolve the phylogeny of Isopoda, we hypothesised that their remarkably different life histories may have produced asymmetrical adaptive evolutionary pressures on their (mito)genomes, which in turn resulted in compositional heterogeneity and heterotachy that interfere with the reconstruction of their evolutionary history.

We tested the performance of several methodologies commonly used to account for these compositional imbalances: the use of amino acid sequences (instead of nucleotide sequences), data partitioning, and phylogenetic algorithms designed for non-homogeneous data (heterogeneous CAT-GTR and heterotachous IQ-GHOST). We also tested commonly used algorithms (ML, BI and Parsimony), and a large number of different datasets. Apart from the heterotachous GHOST model, which tended to produce results identical to the common ML analysis, all other variables produced notable impacts on the topology, with unique topologies by far outnumbering identical topologies.

As regards the major unresolved issue of the isopod phylogeny, the sister-clade to all other isopods (basal clade) and the monophyly of Cymothoida, mitochondrial nucleotides (NUC) quite consistently produced Cymothoidae and Corallanidae as the basal clade, first followed by Asellota, then by Phreatoicidea, whereas other Cymothoida tended to be scattered throughout the central ‘catch-all’ clade. AAs, however, largely resolved Phreatoicidea as the basal isopod clade, with the remaining isopods split into two sister-clades: 1) Asellota + Cymothoidae/Corallanidae and 2) all remaining taxa. The Parsimony method produced Cymothoidae+Corallanidae at the base, followed by Asellota + (*G. ovalis* + *L. quadripunctata*) using both datasets. PB analysis of AAs dataset produced a remarkably different topology, with Phreatoicidea at the base, followed by Asellota, but Cymothoidae+Corallanidae were relatively derived, and Cymothoida not so deeply paraphyletic. Nuclear (*18S* gene) topology was also strongly affected by the methodology, but consistently resolved Asellota as the basal isopod clade (Limnoriidea in Parsimony), and Cymothoida as paraphyletic (nested Limnoriidea, not in Parsimony), but highly derived. Even though it appears that we did not manage to reach a conclusion, as we failed to infer a stable topology, we did manage to identify another feature of isopod mitogenomes that may help us in this quest.

### Strand compositional bias: AT and GC skews

Organellar genomes often exhibit a phenomenon known as strand asymmetry, or strand compositional bias, where positive AT skew values indicate more A than T on the strand, positive GC skews indicate more G than C, and vice versa (Reyes, Gissi, Pesole, & Saccone, 1998; Wei et al., 2010). A likely cause for this is hydrolytic deamination of bases on the leading strand when it is single stranded, i.e. during replication and transcription (Bernt, Braband, Schierwater, & Stadler, 2013; Fonseca, Harris, & Posada, 2014; Reyes et al., 1998). Whereas other crustacean taxa usually exhibit positive overall AT skews for genes located on the plus strand (or majority strand) and negative GC skews for genes on the minus strand (minority strand) (Hassanin, 2006; Wei et al., 2010), isopod mitogenomes usually exhibit negative overall AT skews and positive GC skews of the majority strand (Kilpert et al., 2012; Kilpert & Podsiadlowski, 2006; Yu et al., 2018). This is believed to be a consequence of an inversion of the replication origin (RO), where the changed replication order of two mitochondrial DNA strands consequently resulted in an inversed strand asymmetry (Bernt et al., 2013; Hassanin et al., 2005; Kilpert et al., 2012; Kilpert & Podsiadlowski, 2006; Wei et al., 2010).

It is known from before that *Asellus aquaticus* (Asellota) possesses an inversed skew in comparison to other isopod taxa (Kilpert & Podsiadlowski, 2006), but we found that the three available Cymothoidae and Corallanidae species (*C. indica*, *T. chinensis* and *I. japonensis*) also exhibit inversed skew patterns (Fig. 7, Table 1). As regards other studied non-isopod Malacostraca, they exhibit GC skews very similar to the isopod outliers, from −0.014 in *Metacrangonyx repens* to −0.332 in *Typhlatya miravetensis*, and mixed AT skews (negative in Mysida and Amphipoda, and mixed positive and negative in Stomatopoda and Decapoda). As intra-genomic and inter-specific variations in base composition have a strong power to bias phylogenetic analyses (Romiguier & Roux, 2017), and skew-driven LBA phylogenetic artefacts have been reported in arthropods (Hassanin, 2006; Hassanin et al., 2005) and other metazoans (Sun, Li, Kong, & Yu, 2018), we suspect that inversed skews in some isopods may produce such artefacts among the branches exhibiting similar skews.

**Figure 7.**
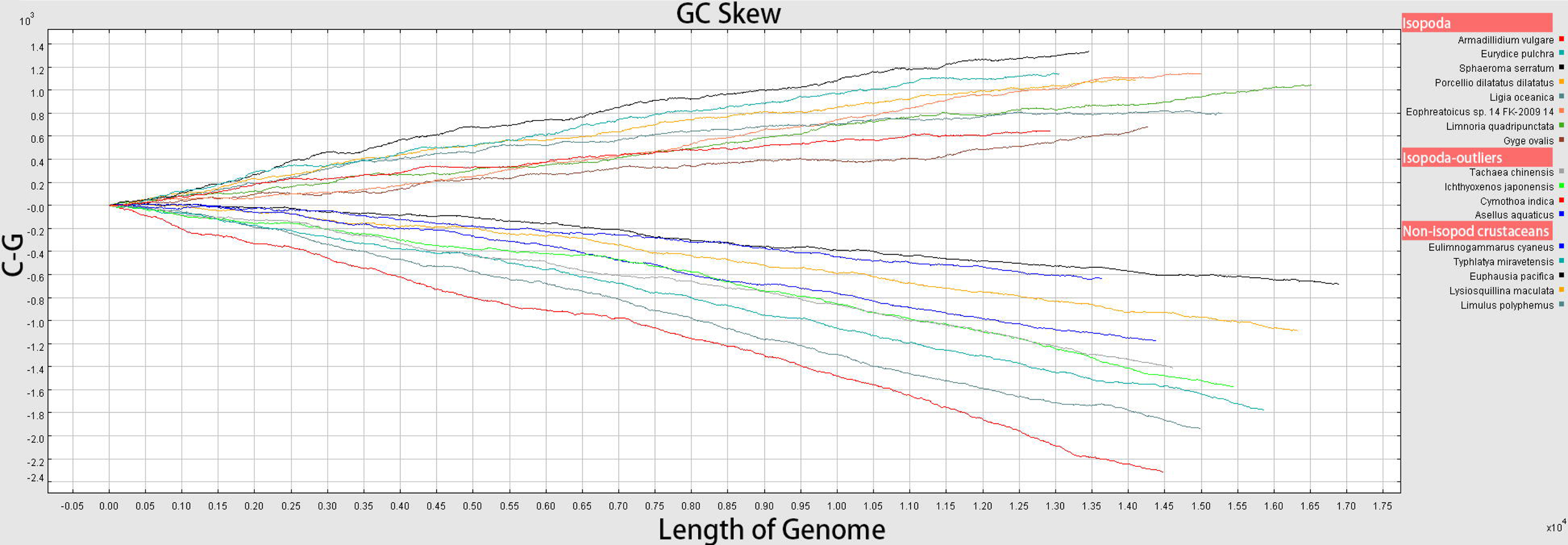
Cumulative GC skews of the majority strands of a selected subset of mitogenomes used for phylogenetic analyses.

### Collation of evolutionary hypotheses

The inversed skew in *Asellus aquaticus* led Kilpert and Podsiadlowski (Kilpert & Podsiadlowski, 2006) to speculate that this clearly suggests that this taxon branched off first in the isopod phylogeny, but a few years later Kilpert et al. (Kilpert et al., 2012) noticed that the basal position of Phreatoicidea causes a conflict in explaining the inversed isopod GC skew (present in *Eophreatoicus* sp.; Fig. 4 – red stars). Our NUC dataset would mostly support a modified version of the first scenario, the replication origin inversion occurred in isopods after Cymothoidae+Corallanidae and Asellota branched off from the main isopod lineage (Figs. 1 and 2 – red star signs), whereas our AAs analyses would mostly support a scenario where the RO inversion occurred only in the common ancestor of Asellota+(Cymothoidae+Corallanidae) clade (Fig. 3). However, there is no support for either of these scenarios from nuclear (*18S*) or morphological (G D F Wilson, 2009) data, which relatively consistently indicate that Asellota (*18S*) or Asellota+Phreatoicidea (morphology) are the basal clade. Parsimony analyses also complicate these scenarios by Asellota forming a sister-clade with *G. ovalis* and *L. quadripunctata*. Importantly, our PB (heterogeneous algorithm) analysis of the AAs dataset produced an mtDNA topology that exhibited notable similarity to the *18S* and morphology-based topologies, where Cymothoidae+Corallanidae clustered with *E. pulchra* (Cirolanidae) in the relatively derived part of the isopod clade. From that, we can conclude that a combination of AAs dataset (which is expected to be less affected by skews than nucleotides) and a heterogeneous CAT-GTR model was most successful in attenuating the phylogenetic artefacts caused by compositional biases. We can therefore reject the above scenarios, and conclude that inversed skews of Asellota and Cymothoidae+Corallanidae are non-synapomorphic. The inversed skew of highly derived Cymothoidae+Corallanidae produces an LBA artefact of them clustering at the base of the isopod clade, phylogenetically close to other taxa with similar (homoplastic) skew patterns (Asellota and non-isopod Malacostraca).

Having established this, now we can use these results to infer the most parsimonious hypothesis for the course of events in the evolutionary history of Isopoda. First, we can reject with confidence the basal position of Cymothoidae+Corallanidae as an artefact. This indicates that the basal isopod taxon is either Asellota (G D F Wilson, 2009), Phreatoicidea (R. C. Brusca & Wilson, 1991), or Asellota+Phreatoicidea sister-clade (Dreyer & Wgele, 2001; Kilpert et al., 2012; George D.F. Wilson, 1999). The latter two scenarios are less parsimonious, as they would require at least three independent RO inversions in the evolutionary history of Isopoda (in the ancestral isopod, in Asellota, and in Cymothoidae+Corallanidae), whereas the first scenario is more parsimonious, as it requires only two (in the ancestral isopod after the split of Asellota and in Cymothoidae+Corallanidae; Fig. 6 red stars). Additionally, the *18S* dataset relatively consistently resolved Asellota as the basal branch (disregarding paraphyletic Isopoda and Parsimony analysis). Therefore, we can tentatively conclude that multiple evidence supports the original hypothesis of Kilpert and Podsiadlowski (Kilpert & Podsiadlowski, 2006): Asellota is the oldest isopod branch and RO inversion in isopods occurred after the Asellota branched off. This scenario implies either a homoplastic nature of the inverted skews in Asellota and Cymothoidae+Corallanidae, or an introgression event from Asellota. Although the latter scenario would directly explain the phylogenetic affinity between the two taxa, the fact that PB (and especially PB+AAs) analyses managed to attenuate this artefact is a strong indication that we can reject this hypothesis, and conclude that homoplastic skews in these taxa are driven by architectural rearrangements.

Although this resolves the issue of the deep paraphyly of Cymothoida, i.e., places the rogue Cymothoidae+Corallanidae clade back within the remaining Cymothoida, *18S* data still resolve the Cymothoida as divided into two clades (Dajidae+Bopyridae and Cirolanidae+Corallanidae+Cymothoidae), and rendered paraphyletic by the nested Limnoriidea. However, as Limnoriidea were resolved as the basal isopod taxon in the *18S* Parsimony analysis, we suspect that this is an LBA between two rogue taxa exhibiting elevated evolutionary rates, and thus erratic phylogenetic behaviour: Corallanidae+Cymothoidae and Limnoriidea. The monophyly of Corallanidae is unsupported by our *18S* analyses, so it will be needed to sequence further mitogenomic (to identify skews) and nuclear molecular data for these three families. This combination of skews and nuclear data would enable us to identify the exact point in evolutionary history where the RO inversion occurred in these taxa, and infer the most parsimonious topology and/or taxonomy, i.e., the one that supports a single RO inversion (or introgression event), as opposed to those that would require multiple events.

As regards other unresolved issues in the phylogeny of Isopoda, our analyses further corroborate the existence of several rogue taxa that exhibit somewhat erratic topological behaviour. The position of *Ligia oceanica* (nominally Oniscidea: Ligiidae), a recognized rogue taxon (Luana S.F. Lins et al., 2017; Shen et al., 2017; G D F Wilson, 2009; Yu et al., 2018), was mostly resolved at the base of the catch-all clade in mtDNA analyses, but in the (putatively) most reliable mitochondrial topology AAs+PB, it was resolved as the basal Oniscidea species. Although this is in perfect agreement with morphological data, which resolve Ligiidae as the most primitive Oniscidea clade (Schmidt, 2008), we cannot claim that this issue is fully resolved, because the entire Ligiidae family exhibited rogue behaviour in the *18S* dataset as well. Three Cymothoida taxa in the mtDNA dataset that exhibit standard isopod skews, *Bathynomus* sp., *E. pulchra* (both Cirolanidae) and *G. ovalis* (Bopyridae), also exhibited rather instable topological behaviour, and we did not find support for their monophyly using the mtDNA data. Although the Cirolanidae are believed to be ancient (Wetzer, 2002) and highly plesiomorphic within this suborder (Brandt & Poore, 2003), this is not supported by the *18S* dataset. A putatively relevant observation is that all three species exhibit unique, highly rearranged, gene orders (Fig. 8). As another rogue species, *L. quadripunctata* (Limnoriidea) (Luana S.F. Lins et al., 2017; G D F Wilson, 2009; Zou et al., 2018), also exhibits a highly rearranged gene order (Lloyd et al., 2015), and as there is evidence of a close positive correlation between the mitogenomic architectural instability and the mutation rate (Hassanin, 2006; Shao, Dowton, Murrell, & Barker, 2003; Xu, Jameson, Tang, & Higgs, 2006), we hypothesise that frequent genome rearrangements may have resulted in an accelerated mutational rate in these species. In agreement with this hypothesis, *E. pulchra* and *L. quadripunctata* exhibited the highest evolutionary rates of all studied isopod (and decapod) mitogenomes (Shen et al., 2017). However, as *L. quadripunctata* was resolved as the basal isopod clade in one study (mitogenomic AAs dataset + BI analysis; Cymothoidae+Corallanidae unavailable at the time) (Luana S.F. Lins et al., 2017), and Limnoriidea exhibited rather erratic behaviour in our *18S* dataset, we can safely argue that mitogenomic rearrangements are unsatisfactory explanation for some of the observed phenomena. Accelerated rates of substitution in some Arthropoda were previously explained by three main factors: genomic rearrangements, including duplication of the control region and gene translocation, parasitic lifestyle, and small body size (Hassanin, 2006). We therefore hypothesise that a proportion of these phylogenetic artefacts are driven by adaptive evolution, which also produces compositional biases that are interfering with phylogenetic reconstruction by producing disproportionately long branches. Furthermore, inverted mitogenomic skews also do not explain the disproportionately long branch of Cymothoidae in the *18S* dataset. We hypothesise that frequent life-history innovations in Cymothoida (Hata et al., 2017; Luana S.F. Lins et al., 2017) may be producing accelerated substitution rates in some of these taxa. Therefore, we can tentatively conclude that mitochondrial evolution in isopods is a result of interplay between adaptive and nonadaptive evolutionary pressures, where non-adaptive outweigh the adaptive in some taxa, such as Cymothoidae and Corallanidae.

**Figure 8.**
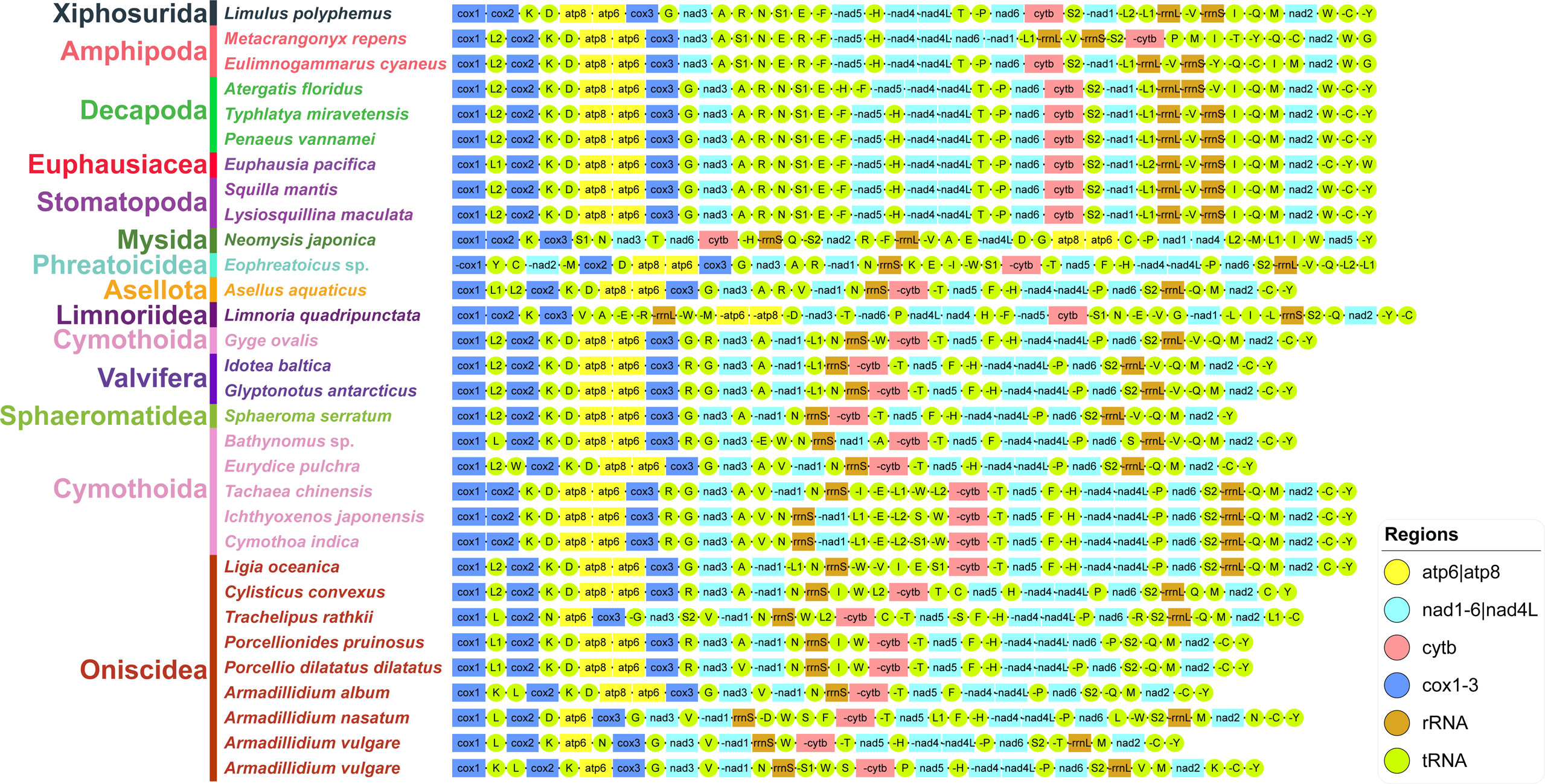
Gene orders in the mitogenomes of Isopoda (and selected Malacostraca).

The topology of non-isopod Malacostraca was rather instable as well, with PB producing notably different topologies from other analyses. As non-isopod Malacostraca also exhibit notable variability in skews (Table 1), we conclude that compositional heterogeneity also interfered with phylogenetic reconstruction. The most relevant question for this study is that of the sister-group to Isopoda: Mysida+Amphipoda in most of our mtNUC analyses (except PB, a unique topology), BI+AAs, and all Parsimony topologies; Mysida in most AAs analyses (ML+GHOST+PB); and Amphipoda in the PB *18S* analysis (Amphipoda were nested within the Isopoda in ML and BI *18S* topologies). As Amphipoda are considered to be the most prominent contender for this position (G D F Wilson, 2009), this again indicates that a much smaller marker in combination with the CAT-GTR model may be producing more reliable results than complete mitogenomes.

### Methodological implications

Our findings indicate that phylogenetic reconstruction using mitochondrial data in Isopoda is severely hampered by compositional biases. This corroborates an earlier observation that in isopods mitochondrial sequence data are prone to producing artefactual LBA relationships, and thus a poor tool for phylogenetic reconstruction in this taxon (Wetzer, 2002). Intriguingly, these did not affect only the nucleotide dataset, but also the amino acid dataset. Amino acid datasets should be less affected by nonadaptive compositional biases, as nonsynonymous mutations are likely to be affected by the purifying selection (as opposed to synonymous mutations). However, it has been shown that mitochondrial strand asymmetry (skews) can have very large effects on the composition of the encoded proteins (Botero-Castro et al., 2018; Min & Hickey, 2007). Some groups of residues share similar physico-chemical properties and can be conservatively exchanged without significant functional impacts (Botero-Castro et al., 2018), and some parts of protein chains may evolve under lesser functional constraints, so many non-synonymous mutations may not affect the protein functionally. Finally, as argued above, adaptive evolutionary pressures are also likely to affect the composition of proteins in isopods.

All individual genes produced unique topologies, which has important implications for the interpretation of previous results inferred using single-gene datasets. This phenomenon has been observed in isopods before on a much smaller scale (Wetzer, 2002), and it is in agreement with the proposed mosaic nature of (mitochondrial) genomes, where different loci often produce conflicting phylogenetic signals (Degnan & Rosenberg, 2009; Pollard, Iyer, Moses, & Eisen, 2006; Romiguier & Roux, 2017). It should be noted that some genes produced less pronounced phylogenetic artefacts, notably *cytb* and *12S* (File S2), which indicates that these two genes might be evolving under a very strong purifying selection. The latter gene (*12S*) produced a topology that exhibited remarkable congruence with the nuclear *18S* topology, especially in placing Cymothoidae+Corallanidae in the derived part of the clade. Although this corroborates the observation that some genes produce less biased phylogenies than others (Romiguier, Ranwez, Delsuc, Galtier, & Douzery, 2013), the artefact of Asellota clustering within the derived Cymothoida clade shows that compositional biases affected this dataset as well, and that we can safely conclude that single-gene mitochondrial markers are not a suitable tool for this task. Similarly, gene orders produced topological instability, very low support, and almost nonsensical topologies, with paraphyletic Isopoda. As highly rearranged gene orders were at the base of the isopod clade, we hypothesise that the discontinuous evolution of mitogenomic architecture evolution (Zou et al., 2017) produces phylogenetic artefacts, such as LBA. This corroborates the hypothesis that gene orders are not a useful phylogenetic marker in lineages exhibiting destabilised mitogenomic architecture (Zhang et al., 2017; Zou et al., 2017).

All of the tested standard models (BI, ML and Parsimony) were very sensitive to compositional biases, and produced highly misleading artefacts. We also tested the performance of a new (experimental) GHOST heterotachous model (Crotty et al., 2017), but we found that it produces results almost identical to the common ML algorithm, so we conclude that on this dataset the model appears to be largely useless. Importantly, as regards the aforementioned unresolved feud about the most suitable methodological approach to account for compositional heterogeneity (Feuda et al., 2017; Whelan & Halanych, 2017), our results indicate that (in isopods) the CAT-GTR model by far outperforms the partitioning (assigning different evolutionary models to different partitions). Although we discourage the use of mitochondrial data as a tool for phylogenetic reconstruction in isopods, there are other available methodological approaches designed to account for this problem (Hassanin, 2006; Richards, Brown, Barley, Chong, & Thomson, 2018; Sheffield et al., 2009; Yang, Li, Dang, & Bu, 2018), so future studies may attempt to test their performance.

## Conclusions

With respect to our working hypothesis, we can accept the first part of it: asymmetrical mutational pressures generate compositional heterogeneity in isopod mitogenomes and interfere with phylogenetic reconstruction. However, we were mistaken in assuming that these mutational pressures are primarily adaptive, i.e., caused by their radically diverse life histories. Our results imply that mitochondrial evolution in isopods is a result of interplay between adaptive and non-adaptive evolutionary pressures, where non-adaptive outweigh the adaptive in some taxa (Cymothoidae and Corallanidae). This is in agreement with a recent observation that mitogenomes in isopods mutate at a rate independent of life history traits (Saclier et al., 2018), and contributes to our understanding of the interplay of adaptive and nonadaptive processes in shaping the mitochondrial genomes of Metazoa (Bernt et al., 2013; Smith, 2016). We can conclude that mitogenomic architectural instability (comprising RO inversions) generates strong compositional biases that render mitogenomic sequence data a very poor tool for phylogenetic reconstruction in Isopoda. The deeply contrasting phylogenetic signals that we identified, not only between nuclear and mitochondrial datasets, but also among different mitogenomic datasets, have important implications both for the interpretation of past studies and for scientists who plan to study the phylogeny of these taxa in the future. Regardless of this, simply by rejecting the previous contradictory hypotheses inferred using mitochondrial data, we managed to at least partially resolve several contentious issues in the phylogeny of Isopoda. As regards the methodological approaches used here to account for compositional heterogeneity, we can conclude that none of the tools managed to fully revolve these biases, but CAT-GTR algorithm outperformed partitioning, and best results were achieved by combining it with the amino acids dataset. Although we discourage the use of mitochondrial data for this purpose, any future study that would aim to rely (even if partially) on mitogenomic data would have to first unambiguously prove that they used tools that successfully account for it. As mtDNA data have played a major role in our current understanding of the evolutionary history of life on Earth (Rubinoff et al., 2005), implications of this study are much broader than its original scope. As our findings show that architectural rearrangements can produce major compositional biases even on short evolutionary timescales, the implications of this study are that proving the suitability of data via GC and AT skew analyses should be a prerequisite for every study that aims to use mitochondrial data for phylogenetic reconstruction, even among closely related taxa. These findings should not discourage scientists from sequencing further isopod mitogenomes, as their architectural hypervariability still makes them a useful tool for unravelling the conundrums of evolution of mitochondrial architecture, and as mitochondrial skews can be used as an additional phylogenetic tool to infer the most parsimonious phylogenetic hypotheses.

## Acknowledgements

This work was supported by the Earmarked Fund for China Agriculture Research System (CARS-45-15); the National Natural Science Foundation of China (31872604, 31572658); and the Deanship of Scientific Research at King Saud University (RGP 1435-012). The funders had no role in the design of the study, collection, analysis and interpretation of data, and in writing the manuscript. The authors declare that they have no conflict of interest. The authors would like to express their sincere appreciation to Chen Rong (Bio-Transduction Lab) for helping us to conduct some of the wet lab experiments, and the Deanship of Scientific Research at King Saud University for funding this research.

## Data Accessibility

DNA sequences: Genbank accessions MK079664, MK542856, MK542857 and MK542858. The remaining data are included within the article and its supplementary files

## Author Contributions

IJ, DZ, HZ, and GTW designed research. DZ, HZ, CJH, WXL, KAAG, FAM, and SM performed research. DZ contributed analytical tools; IJ, DZ, HZ, CJH, WXL, KAAG, FAM, SM and GTW analyzed data. IJ and DZ wrote the paper, and all authors revised it critically for important intellectual content.

## Supporting Information

File S1: best partitioning scheme and model selection

File S2: Phylograms produced by all analyses.

File S3: The 18S dataset with taxonomy details.

File S4: Alignments used in this study.

