## Supplementary material for "Mitochondrial architecture rearrangements produce asymmetrical nonadaptive mutational pressures that subvert the phylogenetic reconstruction in Isopoda": File S2

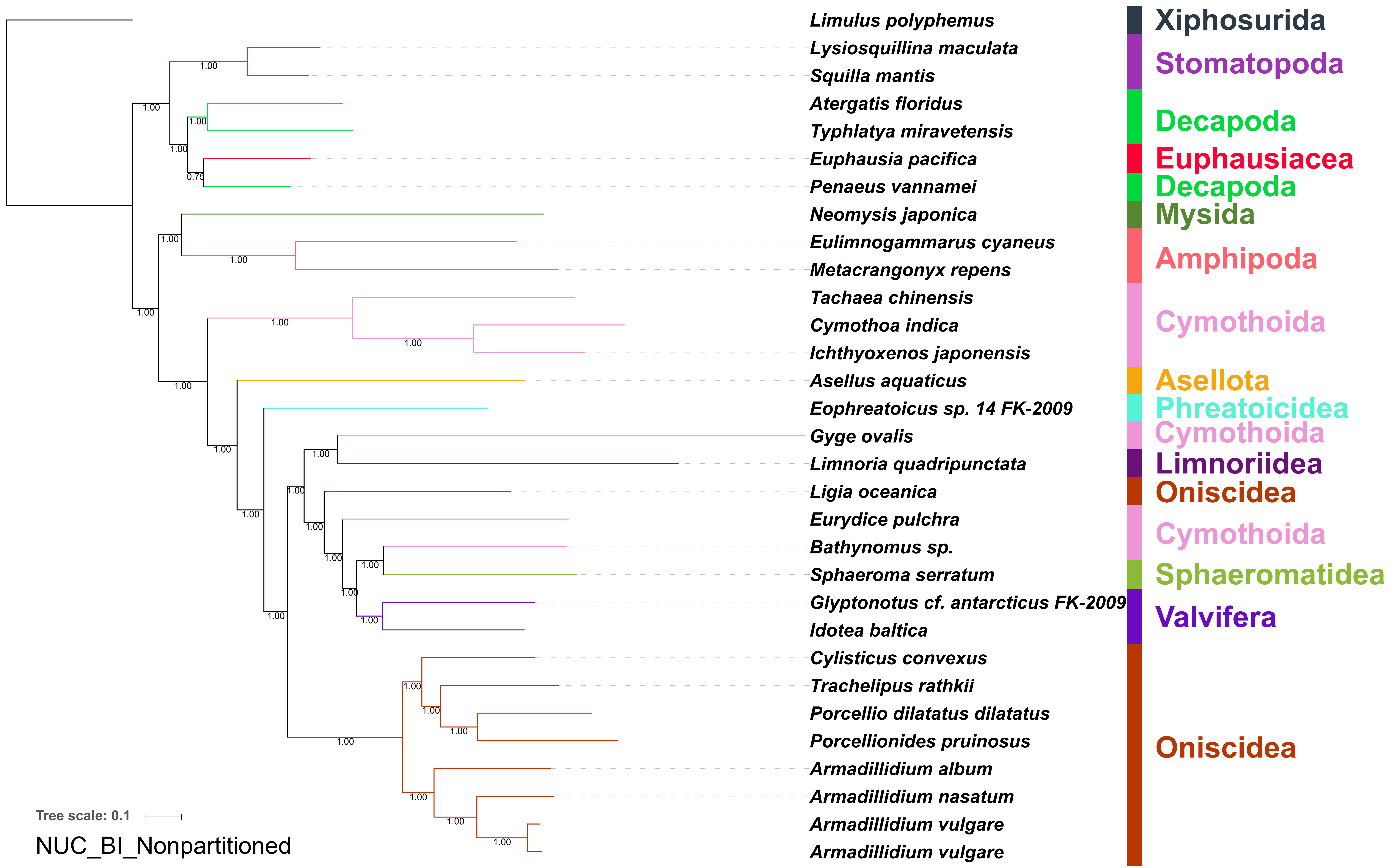
Limulus polyphemus
Lysiosquillina maculata
Squilla mantis
Atergatis floridus
Typhlatya miravetensis
Euphausia pacifica
Penaeus vannamei
Neomysis japonica
Eulimnogammarus cyaneus
Metacrangonyx repens
Tachaea chinensis
Cymothoa indica
Ichthyoxenos japonensis
Asellus aquaticus
Eophreatoicus sp. 14 FK-2009
Gyge ovalis
Limnoria quadripunctata
Ligia oceanica
Eurydice pulchra
Bathynomus sp.
Sphaeroma serratum
Glyptonotus cf. antarcticus FK-2009
Idotea baltica
Cylisticus convexus
Trachelipus rathkii
Porcellio dilatatus dilatatus
Porcellionides pruinosus
Armadillidium album
Armadillidium nasatum
Armadillidium vulgare
Armadillidium vulgare
Xiphosurida
Stomatopoda
Decapoda
Euphausiacea
Decapoda
Mysida
Amphipoda
Cymothoida
Asellota
Phreatoicidea
Cymothoida
Limnoriidea
Oniscidea
Cymothoida
Sphaeromatidea
Valvifera
Oniscidea
Tree scale: 0.1
NUC_BI_Nonpartitioned

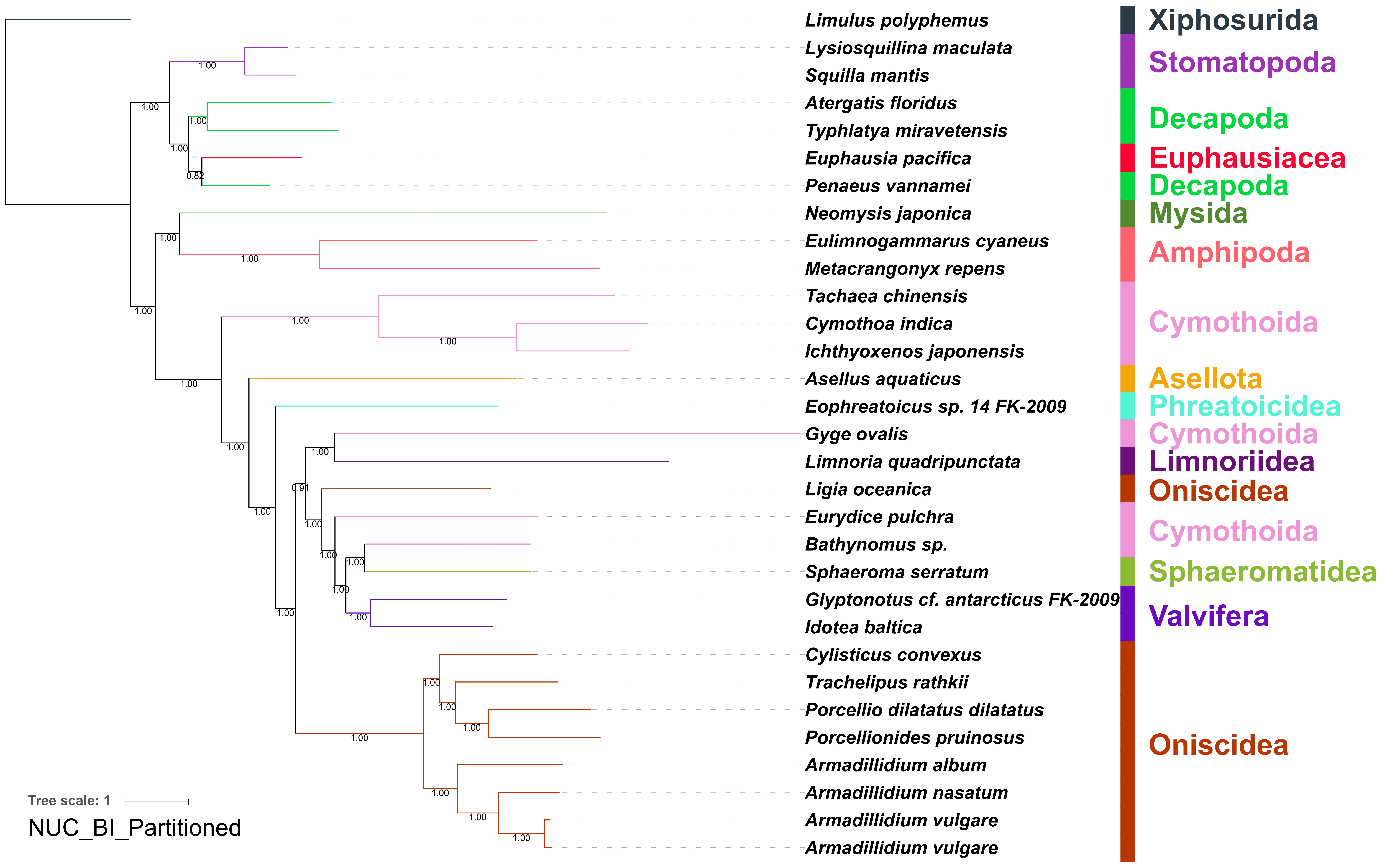

Limulus polyphemus
Lysiosquillina maculata
Squilla mantis
Atergatis floridus
Typhlatya miravetensis
Euphausia pacifica
Penaeus vannamei
Neomysis japonica
Eulimnogammarus cyaneus
Metacrangonyx repens
Tachaea chinensis
Cymothoa indica
Ichthyoxenos japonensis
Asellus aquaticus
Eophreatoicus sp. 14 FK-2009
Gyge ovalis
Limnoria quadripunctata
Ligia oceanica
Eurydice pulchra
Bathynomus sp.
Sphaeroma serratum
Glyptonotus cf. antarcticus FK-2009
Idotea baltica
Cylisticus convexus
Trachelipus rathkii
Porcellio dilatatus dilatatus
Porcellionides pruinosus
Armadillidium album
Armadillidium nasatum
Armadillidium vulgare
Armadillidium vulgare
Xiphosurida
Stomatopoda
Decapoda
Euphausiacea
Decapoda
Mysida
Amphipoda
Cymothoida
Asellota
Phreatoicidea
Cymothoida
Limnoriidea
Oniscidea
Cymothoida
Sphaeromatidea
Valvifera
Oniscidea
Tree scale: 1
NUC_BI_Partitioned

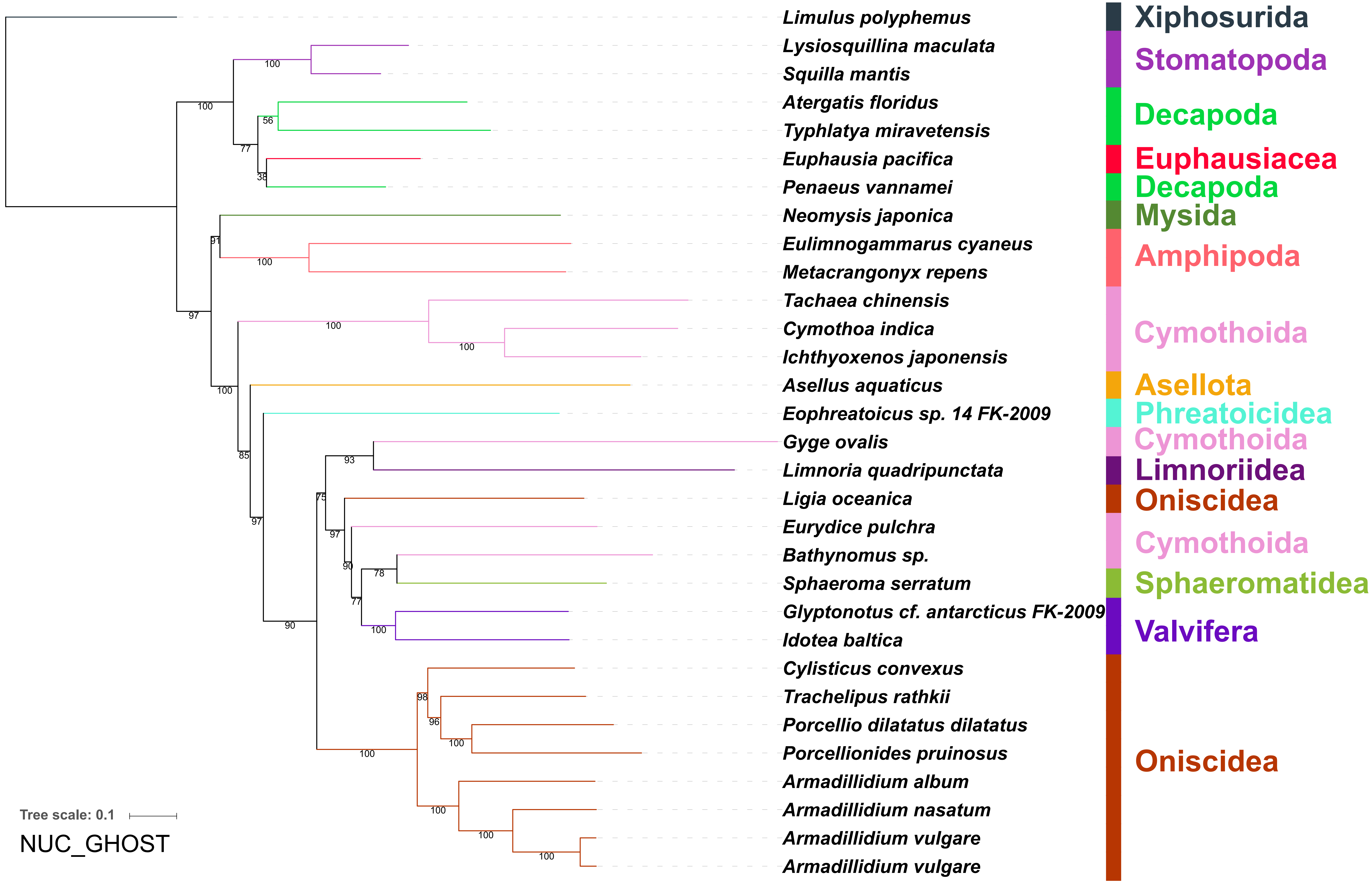
Limulus polyphemus
Lysiosquillina maculata
Squilla mantis
Atergatis floridus
Typhlatya miravetensis
Euphausia pacifica
Penaeus vannamei
Neomysis japonica
Eulimnogammarus cyaneus
Metacrangonyx repens
Tachaea chinensis
Cymothoa indica
Ichthyoxenos japonensis
Asellus aquaticus
Eophreatoicus sp. 14 FK-2009
Gyge ovalis
Limnoria quadripunctata
Ligia oceanica
Eurydice pulchra
Bathynomus sp.
Sphaeroma serratum
Glyptonotus cf. antarcticus FK-2009
Idotea baltica
Cylisticus convexus
Trachelipus rathkii
Porcellio dilatatus dilatatus
Porcellionides pruinosus
Armadillidium album
Armadillidium nasatum
Armadillidium vulgare
Armadillidium vulgare
Xiphosurida
Stomatopoda
Decapoda
Euphausiacea
Decapoda
Mysida
Amphipoda
Cymothoida
Asellota
Phreatoicidea
Cymothoida
Limnoriidea
Oniscidea
Cymothoida
Sphaeromatidea
Valvifera
Oniscidea
Tree scale: 0.1
NUC_GHOST

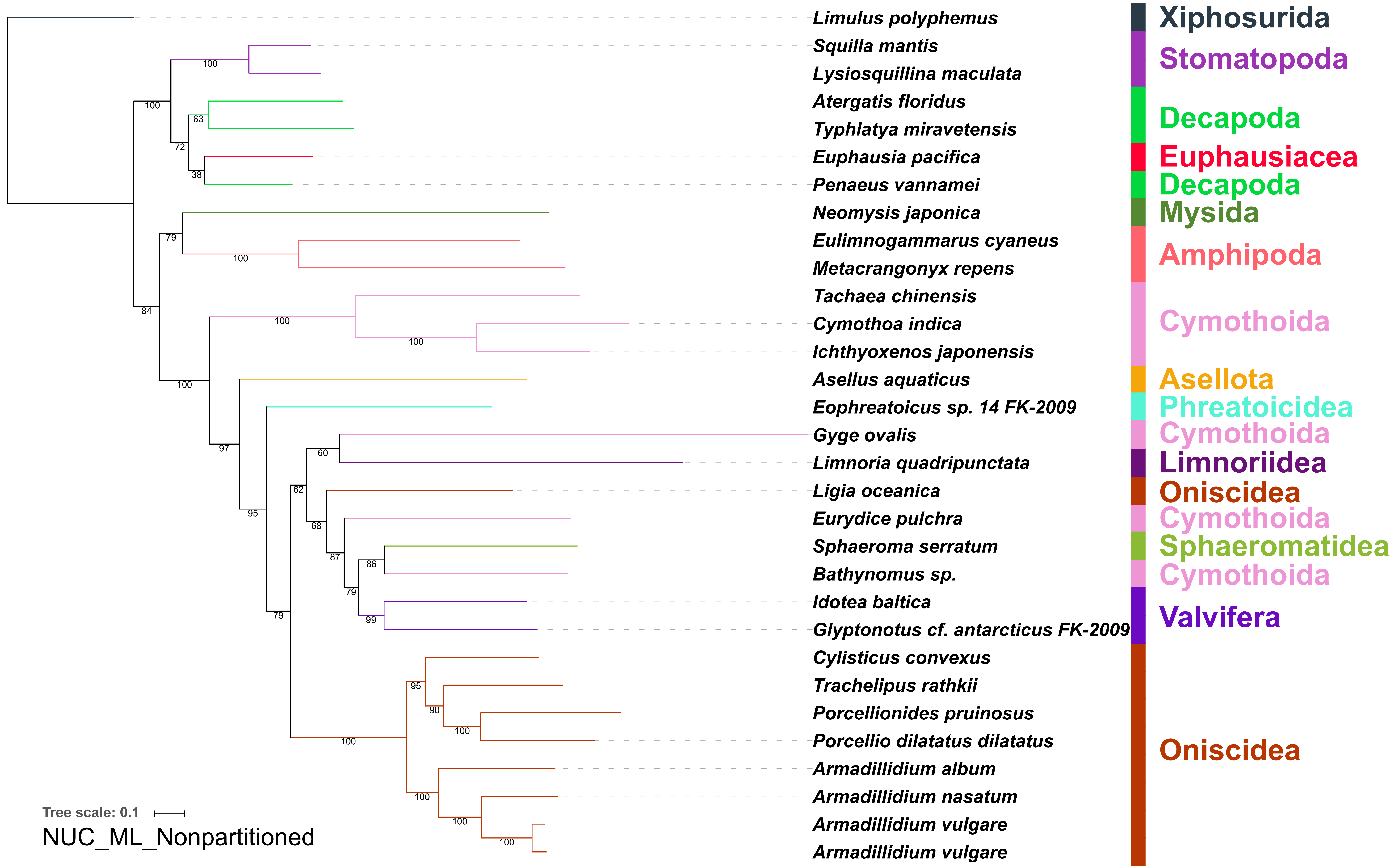

Limulus polyphemus
Squilla mantis
Lysiosquillina maculata
Atergatis floridus
Typhlatya miravetensis
Euphausia pacifica
Penaeus vannamei
Neomysis japonica
Eulimnogammarus cyaneus
Metacrangonyx repens
Tachaea chinensis
Cymothoa indica
Ichthyoxenos japonensis
Asellus aquaticus
Eophreatoicus sp. 14 FK-2009
Gyge ovalis
Limnoria quadripunctata
Ligia oceanica
Eurydice pulchra
Sphaeroma serratum
Bathynomus sp.
Idotea baltica
Glyptonotus cf. antarcticus FK-2009
Cylisticus convexus
Trachelipus rathkii
Porcellionides pruinosus
Porcellio dilatatus dilatatus
Armadillidium album
Armadillidium nasatum
Armadillidium vulgare
Armadillidium vulgare
Xiphosurida
Stomatopoda
Decapoda
Euphausiacea
Decapoda
Mysida
Amphipoda
Cymothoida
Asellota
Phreatoicidea
Cymothoida
Limnoriidea
Oniscidea
Cymothoida
Sphaeromatidea
Cymothoida
Valvifera
Oniscidea
Tree scale: 0.1
NUC_ML_Nonpartitioned

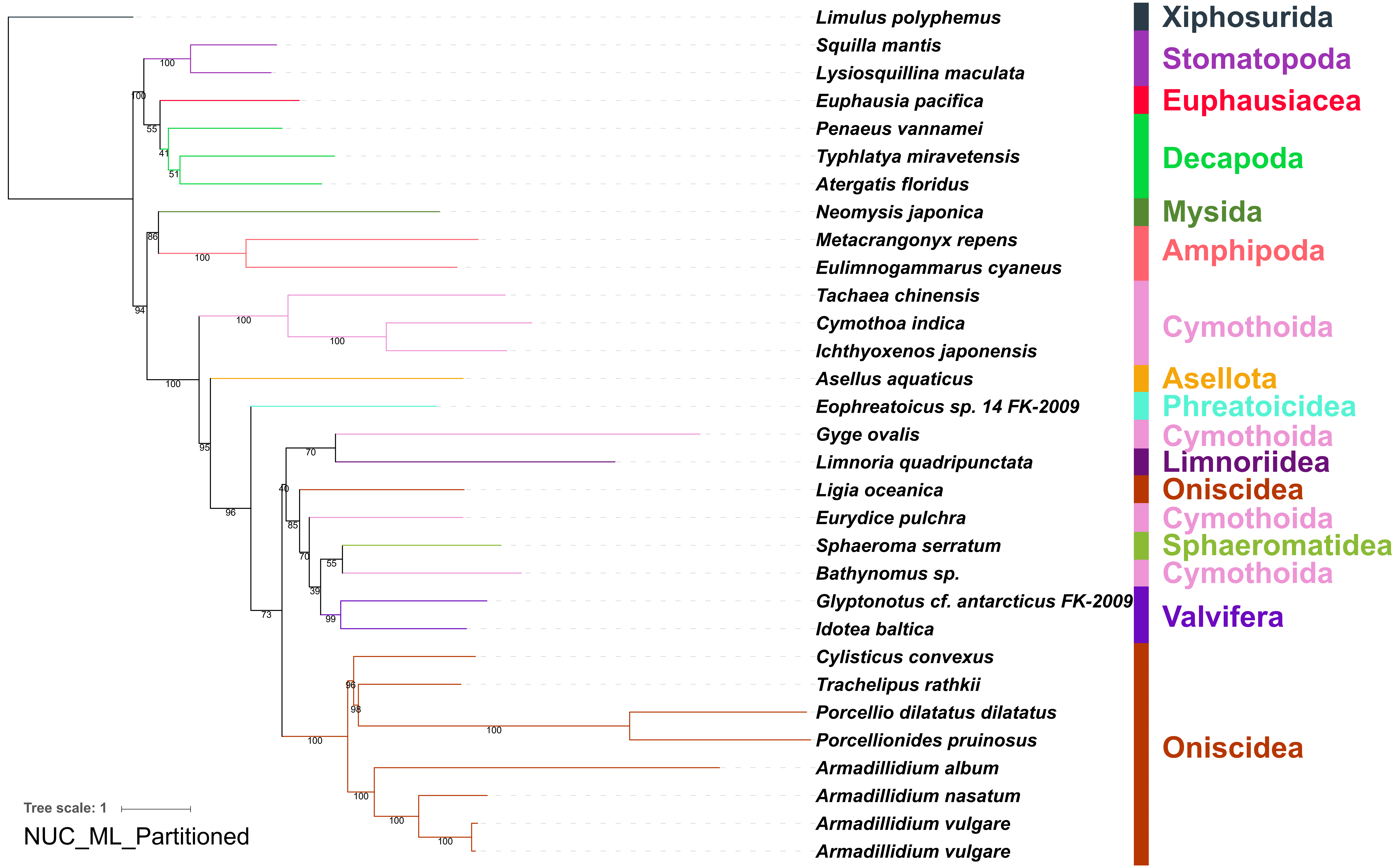
Limulus polyphemus
Squilla mantis
Lysiosquillina maculata
Euphausia pacifica
Penaeus vannamei
Typhlatya miravetensis
Atergatis floridus
Neomysis japonica
Metacrangonyx repens
Eulimnogammarus cyaneus
Tachaea chinensis
Cymothoa indica
Ichthyoxenos japonensis
Asellus aquaticus
Eophreatoicus sp. 14 FK-2009
Gyge ovalis
Limnoria quadripunctata
Ligia oceanica
Eurydice pulchra
Sphaeroma serratum
Bathynomus sp.
Glyptonotus cf. antarcticus FK-2009
Idotea baltica
Cylisticus convexus
Trachelipus rathkii
Porcellio dilatatus dilatatus
Porcellionides pruinosus
Armadillidium album
Armadillidium nasatum
Armadillidium vulgare
Armadillidium vulgare
Xiphosurida
Stomatopoda
Euphausiacea
Decapoda
Mysida
Amphipoda
Cymothoida
Asellota
Phreatoicidea
Cymothoida
Limnoriidea
Oniscidea
Cymothoida
Sphaeromatidea
Cymothoida
Valvifera
Oniscidea
Tree scale: 1
NUC_ML_Partitioned

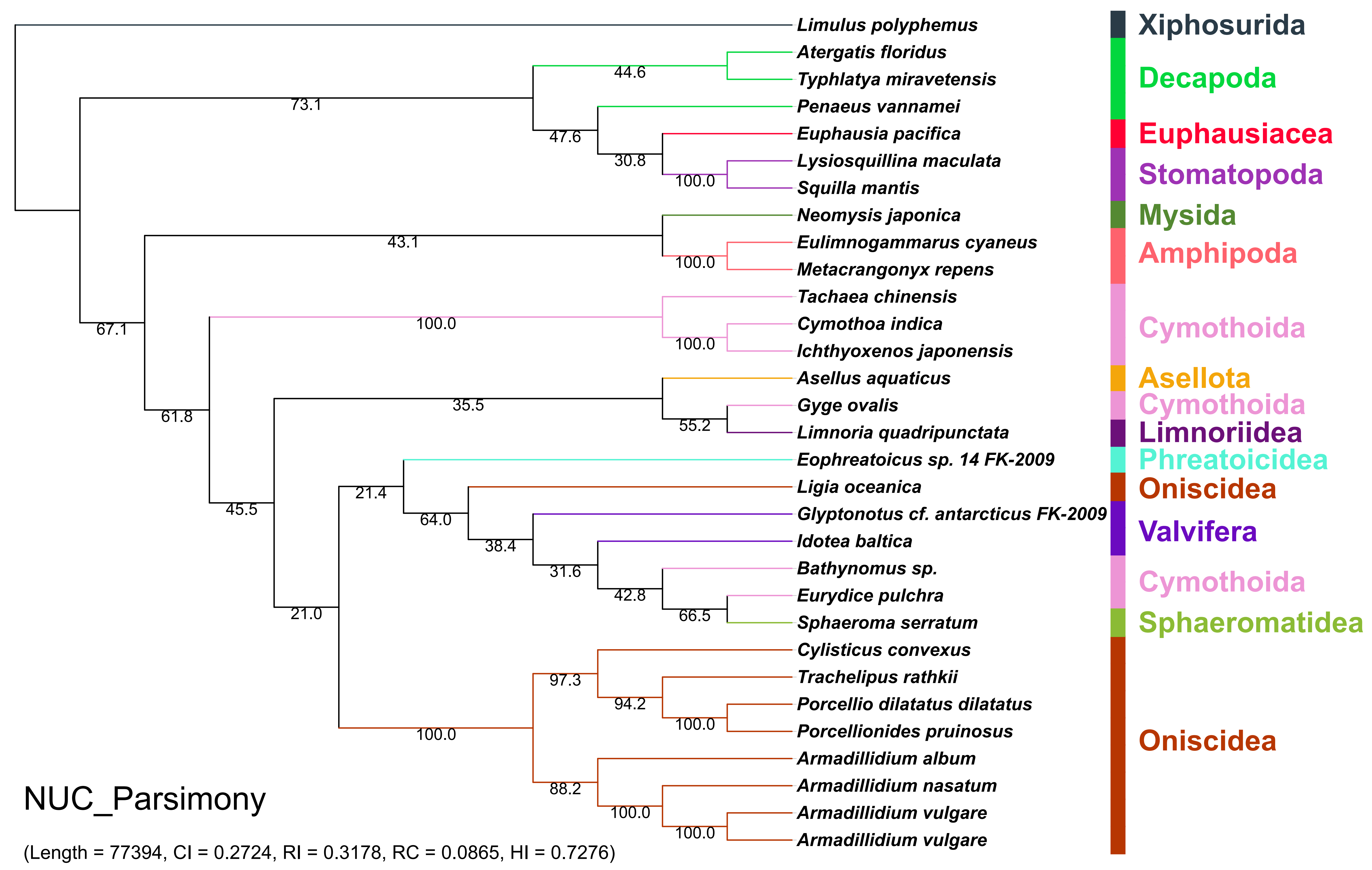
Limulus polyphemus
Atergatis floridus
Typhlatya miravetensis
Penaeus vannamei
Euphausia pacifica
Lysiosquillina maculata
Squilla mantis
Neomysis japonica
Eulimnogammarus cyaneus
Metacrangonyx repens
Tachaea chinensis
Cymothoa indica
Ichthyoxenos japonensis
Asellus aquaticus
Gyge ovalis
Limnoria quadripunctata
Eophreatoicus sp. 14 FK-2009
Ligia oceanica
Glyptonotus cf. antarcticus FK-2009
Idotea baltica
Bathynomus sp.
Eurydice pulchra
Sphaeroma serratum
Cylisticus convexus
Trachelipus rathkii
Porcellio dilatatus dilatatus
Porcellionides pruinosus
Armadillidium album
Armadillidium nasatum
Armadillidium vulgare
Armadillidium vulgare
Xiphosurida
Decapoda
Euphausiacea
Stomatopoda
Mysida
Amphipoda
Cymothoida
Asellota
Cymothoida
Limnoriidea
Phreatoicidea
Oniscidea
Valvifera
Cymothoida
Sphaeromatidea
Oniscidea
44.6
73.1
47.6
30.8
100.0
43.1
100.0
100.0
67.1
100.0
35.5
55.2
61.8
21.4
45.5
64.0
38.4
31.6
42.8
66.5
21.0
97.3
94.2
100.0
100.0
88.2
100.0
100.0
NUC_Parsimony
(Length = 77394, CI = 0.2724, RI = 0.3178, RC = 0.0865, HI = 0.7276)

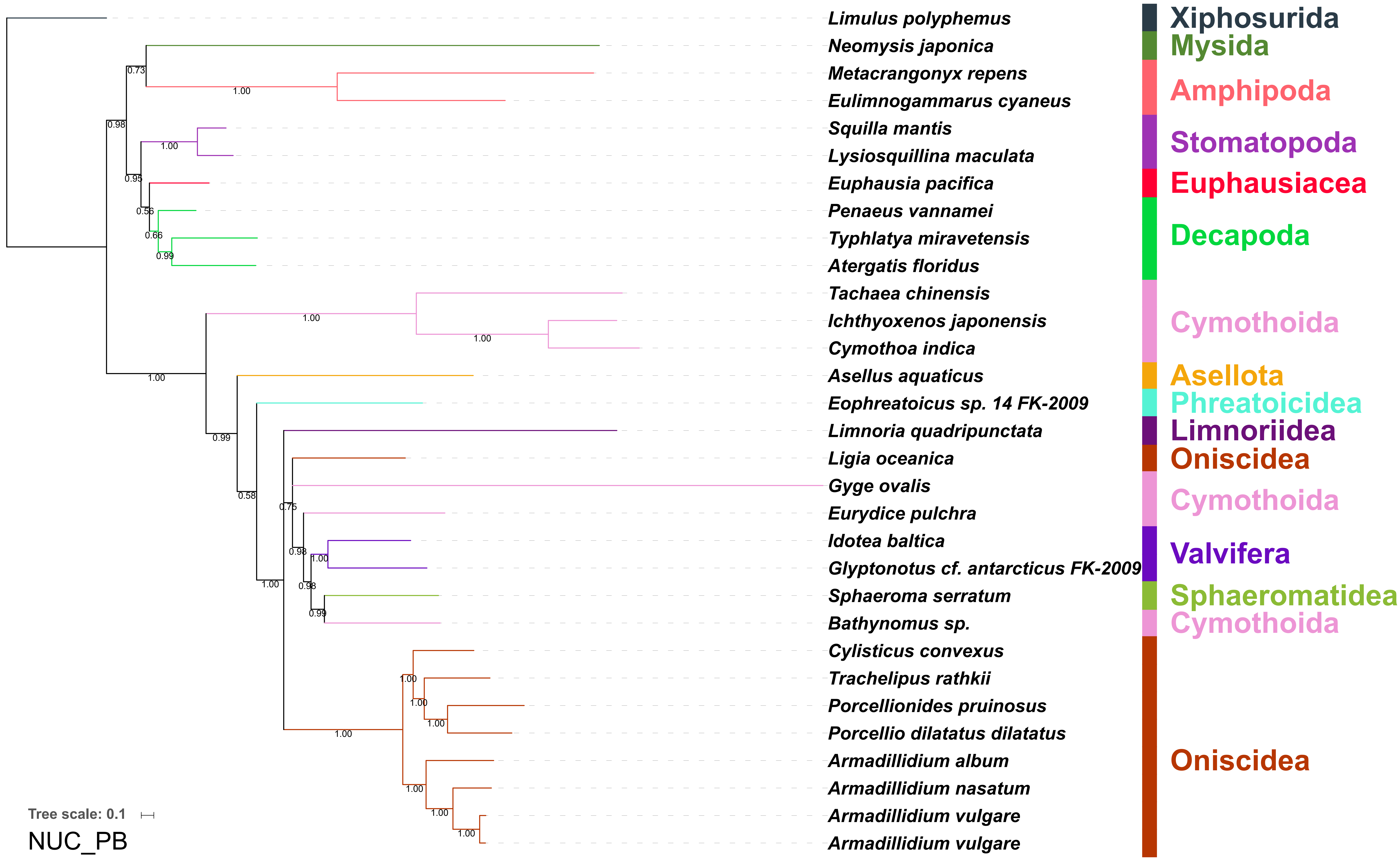
Limulus polyphemus
Neomysis japonica
Metacrangonyx repens
Eulimnogammarus cyaneus
Squilla mantis
Lysiosquillina maculata
Euphausia pacifica
Penaeus vannamei
Typhlatya miravetensis
Atergatis floridus
Tachaea chinensis
Ichthyoxenos japonensis
Cymothoa indica
Asellus aquaticus
Eophreatoicus sp. 14 FK-2009
Limnoria quadripunctata
Ligia oceanica
Gyge ovalis
Eurydice pulchra
Idotea baltica
Glyptonotus cf. antarcticus FK-2009
Sphaeroma serratum
Bathynomus sp.
Cylisticus convexus
Trachelipus rathkii
Porcellionides pruinosus
Porcellio dilatatus dilatatus
Armadillidium album
Armadillidium nasatum
Armadillidium vulgare
Armadillidium vulgare
Xiphosurida
Mysida
Amphipoda
Stomatopoda
Euphausiacea
Decapoda
Cymothoida
Asellota
Phreatoicidea
Limnoriidea
Oniscidea
Cymothoida
Valvifera
Sphaeromatidea
Cymothoida
Oniscidea
Tree scale: 0.1
NUC_PB

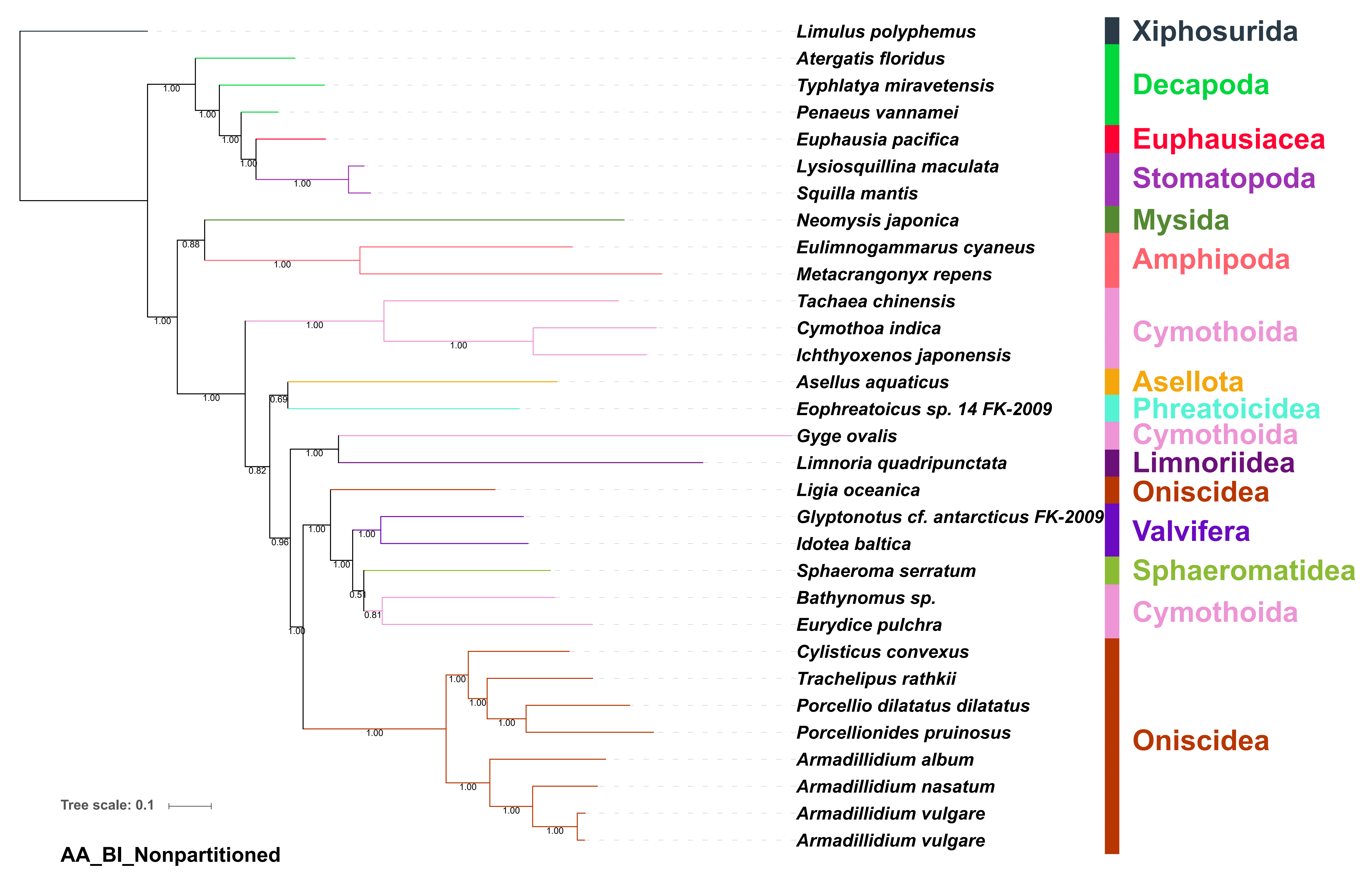
Limulus polyphemus
Atergatis floridus
Typhlatya miravetensis
Penaeus vannamei
Euphausia pacifica
Lysiosquillina maculata
Squilla mantis
Neomysis japonica
Eulimnogammarus cyaneus
Metacrangonyx repens
Tachaea chinensis
Cymothoa indica
Ichthyoxenos japonensis
Asellus aquaticus
Eophreatoicus sp. 14 FK-2009
Gyge ovalis
Limnoria quadripunctata
Ligia oceanica
Glyptonotus cf. antarcticus FK-2009
Idotea baltica
Sphaeroma serratum
Bathynomus sp.
Eurydice pulchra
Cylisticus convexus
Trachelipus rathkii
Porcellio dilatatus dilatatus
Porcellionides pruinosus
Armadillidium album
Armadillidium nasatum
Armadillidium vulgare
Armadillidium vulgare
Xiphosurida
Decapoda
Euphausiacea
Stomatopoda
Mysida
Amphipoda
Cymothoida
Asellota
Phreatoicidea
Cymothoida
Limnoriidea
Oniscidea
Valvifera
Sphaeromatidea
Cymothoida
Oniscidea
Tree scale: 0.1
AA_BI_Nonpartitioned

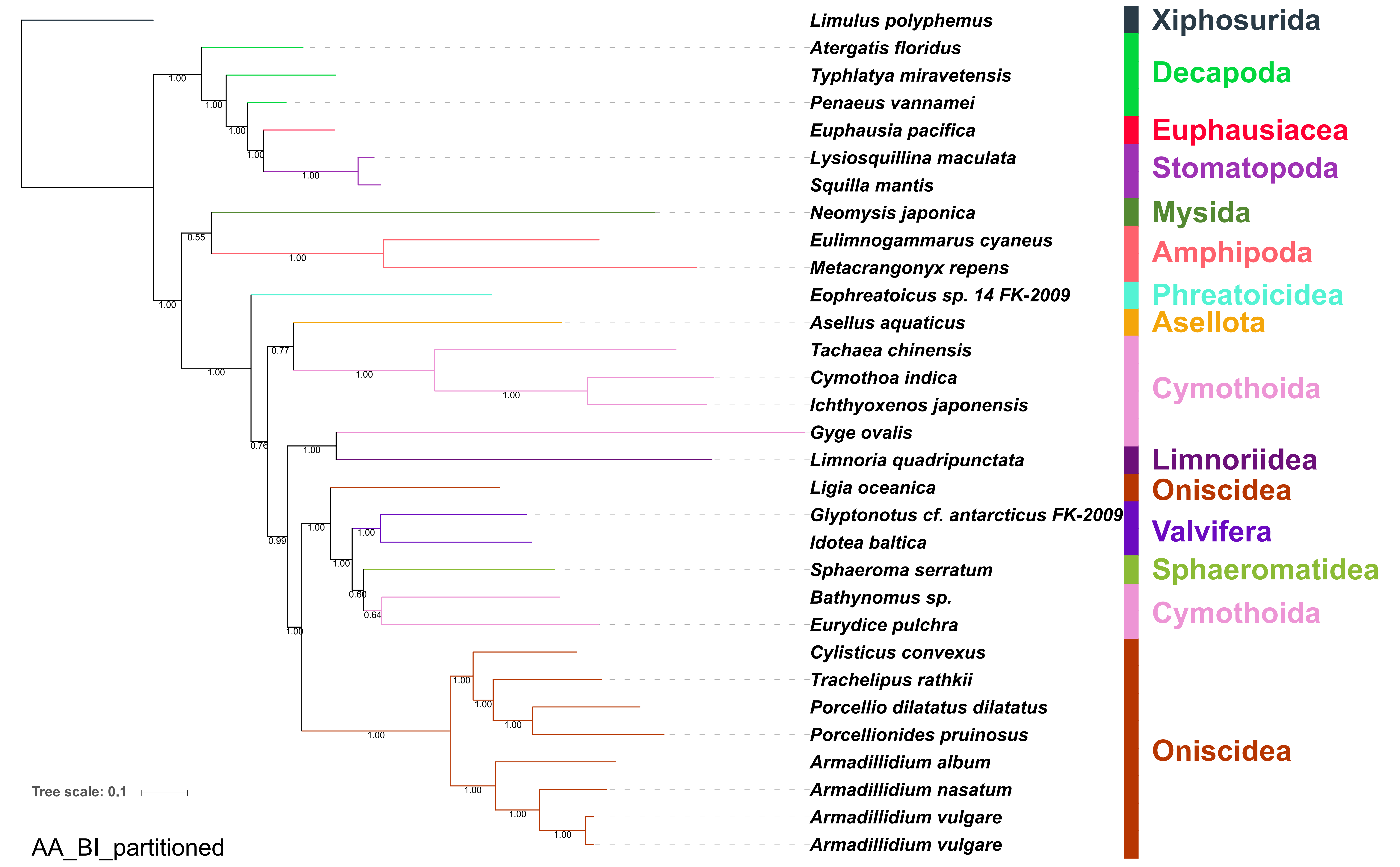

Limulus polyphemus
Atergatis floridus
Typhlatya miravetensis
Penaeus vannamei
Euphausia pacifica
Lysiosquillina maculata
Squilla mantis
Neomysis japonica
Eulimnogammarus cyaneus
Metacrangonyx repens
Eophreatoicus sp. 14 FK-2009
Asellus aquaticus
Tachaea chinensis
Cymothoa indica
Ichthyoxenos japonensis
Gyge ovalis
Limnoria quadripunctata
Ligia oceanica
Glyptonotus cf. antarcticus FK-2009
Idotea baltica
Sphaeroma serratum
Bathynomus sp.
Eurydice pulchra
Cylisticus convexus
Trachelipus rathkii
Porcellio dilatatus dilatatus
Porcellionides pruinosus
Armadillidium album
Armadillidium nasatum
Armadillidium vulgare
Armadillidium vulgare
Xiphosurida
Decapoda
Euphausiacea
Stomatopoda
Mysida
Amphipoda
Phreatoicidea
Asellota
Cymothoida
Limnoriidea
Oniscidea
Valvifera
Sphaeromatidea
Cymothoida
Oniscidea
Tree scale: 0.1
AA_BI_partitioned

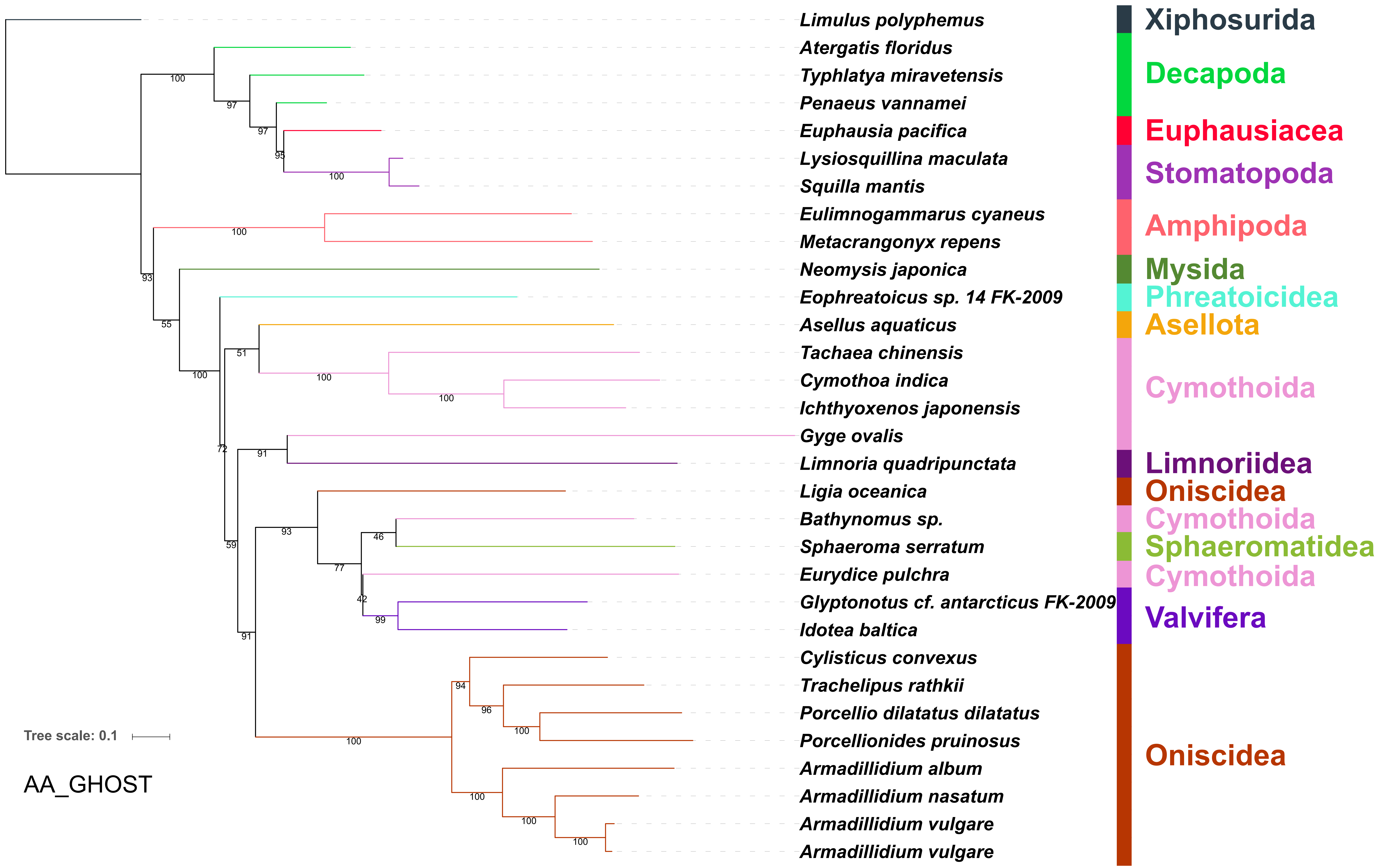

Limulus polyphemus
Atergatis floridus
Typhlatya miravetensis
Penaeus vannamei
Euphausia pacifica
Lysiosquillina maculata
Squilla mantis
Eulimnogammarus cyaneus
Metacrangonyx repens
Neomysis japonica
Eophreatoicus sp. 14 FK-2009
Asellus aquaticus
Tachaea chinensis
Cymothoa indica
Ichthyoxenos japonensis
Gyge ovalis
Limnoria quadripunctata
Ligia oceanica
Bathynomus sp.
Sphaeroma serratum
Eurydice pulchra
Glyptonotus cf. antarcticus FK-2009
Idotea baltica
Cylisticus convexus
Trachelipus rathkii
Porcellio dilatatus dilatatus
Porcellionides pruinosus
Armadillidium album
Armadillidium nasatum
Armadillidium vulgare
Armadillidium vulgare
Xiphosurida
Decapoda
Euphausiacea
Stomatopoda
Amphipoda
Mysida
Phreatoicidea
Asellota
Cymothoida
Limnoriidea
Oniscidea
Cymothoida
Sphaeromatidea
Cymothoida
Valvifera
Oniscidea
Tree scale: 0.1
AA_GHOST

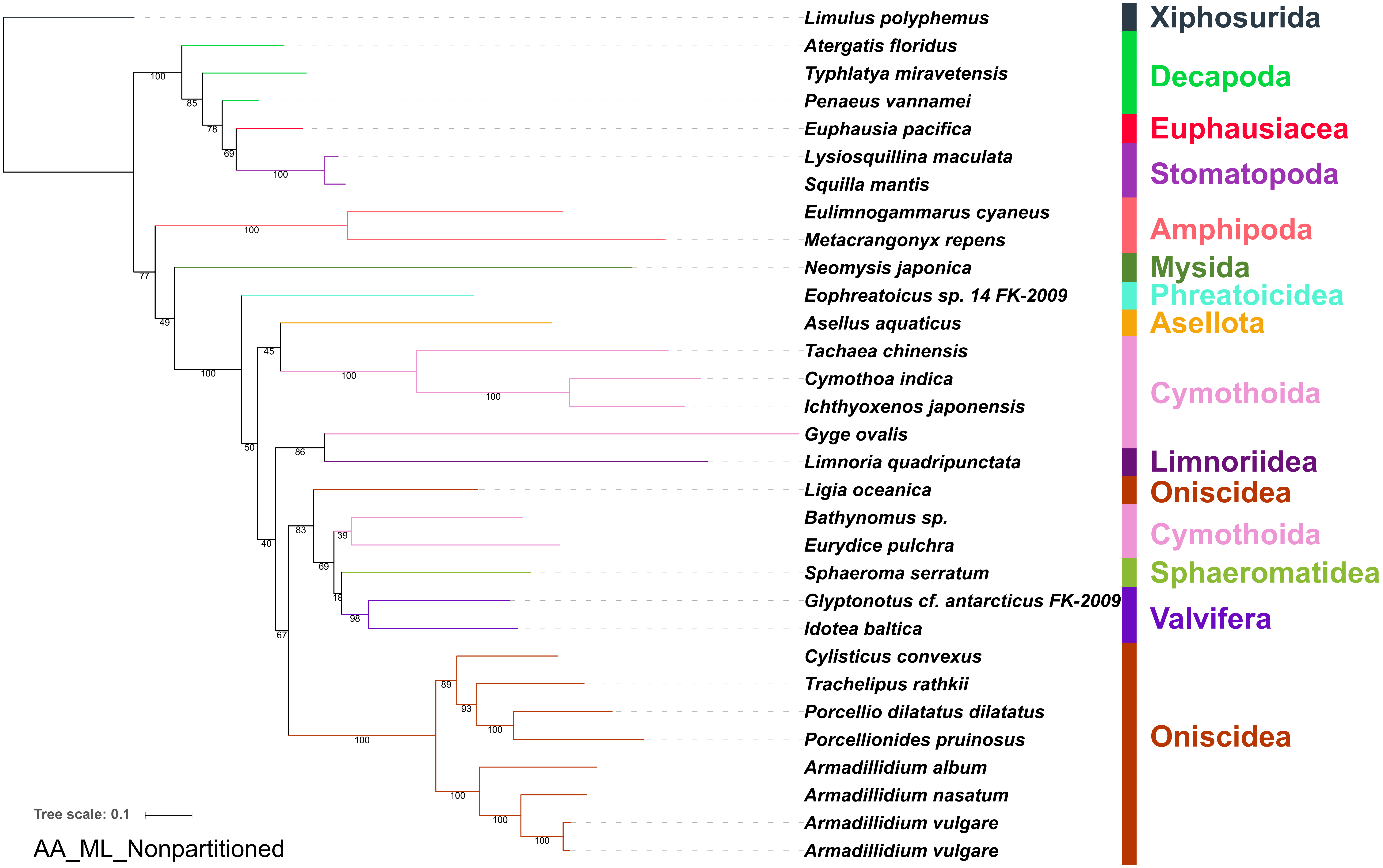
Limulus polyphemus
Atergatis floridus
Typhlatya miravetensis
Penaeus vannamei
Euphausia pacifica
Lysiosquillina maculata
Squilla mantis
Eulimnogammarus cyaneus
Metacrangonyx repens
Neomysis japonica
Eophreatoicus sp. 14 FK-2009
Asellus aquaticus
Tachaea chinensis
Cymothoa indica
Ichthyoxenos japonensis
Gyge ovalis
Limnoria quadripunctata
Ligia oceanica
Bathynomus sp.
Eurydice pulchra
Sphaeroma serratum
Glyptonotus cf. antarcticus FK-2009
Idotea baltica
Cylisticus convexus
Trachelipus rathkii
Porcellio dilatatus dilatatus
Porcellionides pruinosus
Armadillidium album
Armadillidium nasatum
Armadillidium vulgare
Armadillidium vulgare
Xiphosurida
Decapoda
Euphausiacea
Stomatopoda
Amphipoda
Mysida
Phreatoicidea
Asellota
Cymothoida
Limnoriidea
Oniscidea
Cymothoida
Sphaeromatidea
Valvifera
Oniscidea
Tree scale: 0.1
AA_ML_Nonpartitioned

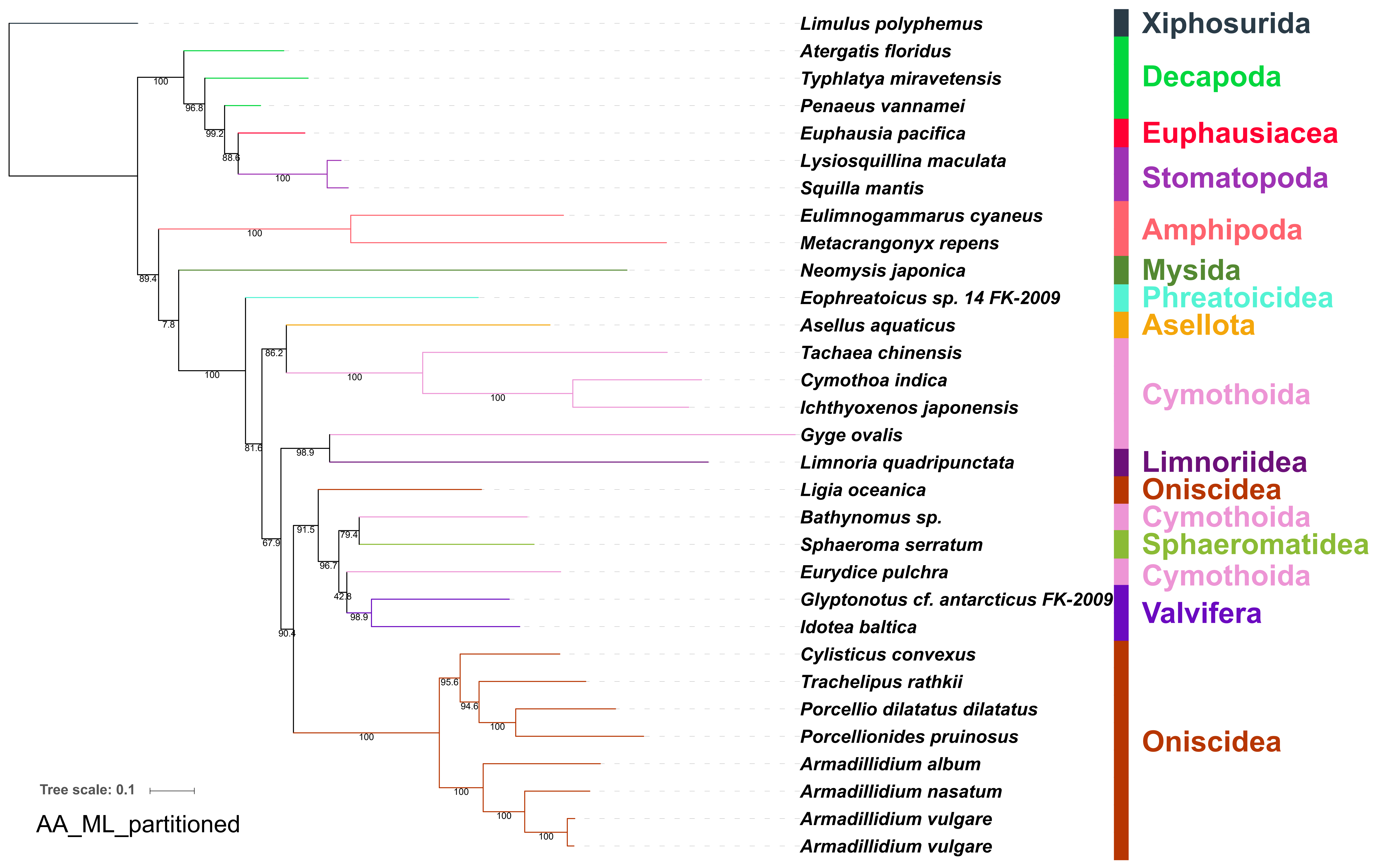
Limulus polyphemus
Atergatis floridus
Typhlatya miravetensis
Penaeus vannamei
Euphausia pacifica
Lysiosquillina maculata
Squilla mantis
Eulimnogammarus cyaneus
Metacrangonyx repens
Neomysis japonica
Eophreatoicus sp. 14 FK-2009
Asellus aquaticus
Tachaea chinensis
Cymothoa indica
Ichthyoxenos japonensis
Gyge ovalis
Limnoria quadripunctata
Ligia oceanica
Bathynomus sp.
Sphaeroma serratum
Eurydice pulchra
Glyptonotus cf. antarcticus FK-2009
Idotea baltica
Cylisticus convexus
Trachelipus rathkii
Porcellio dilatatus dilatatus
Porcellionides pruinosus
Armadillidium album
Armadillidium nasatum
Armadillidium vulgare
Armadillidium vulgare
Xiphosurida
Decapoda
Euphausiacea
Stomatopoda
Amphipoda
Mysida
Phreatoicidea
Asellota
Cymothoida
Limnoriidea
Oniscidea
Cymothoida
Sphaeromatidea
Cymothoida
Valvifera
Oniscidea
100
96.8
99.2
88.6
100
100
89.4
7.8
100
86.2
100
100
81.6
98.9
67.9
91.5
79.4
96.7
42.8
98.9
90.4
95.6
94.6
100
100
100
100
100
Tree scale: 0.1
AA_ML_partitioned

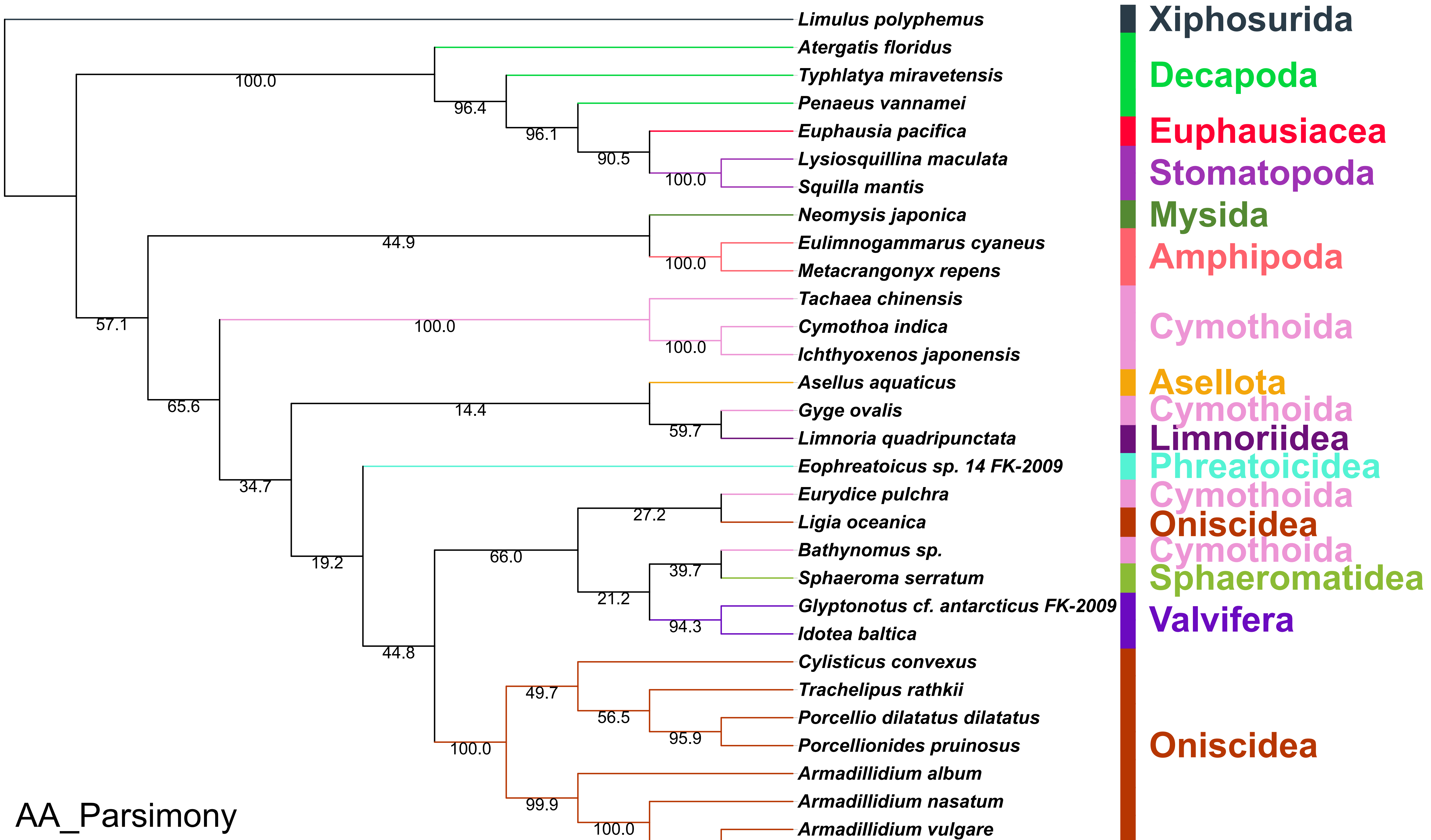


(Length = 19208, CI = 0.5037, RI = 0.4299, RC = 0.2165, HI = 0.4963)

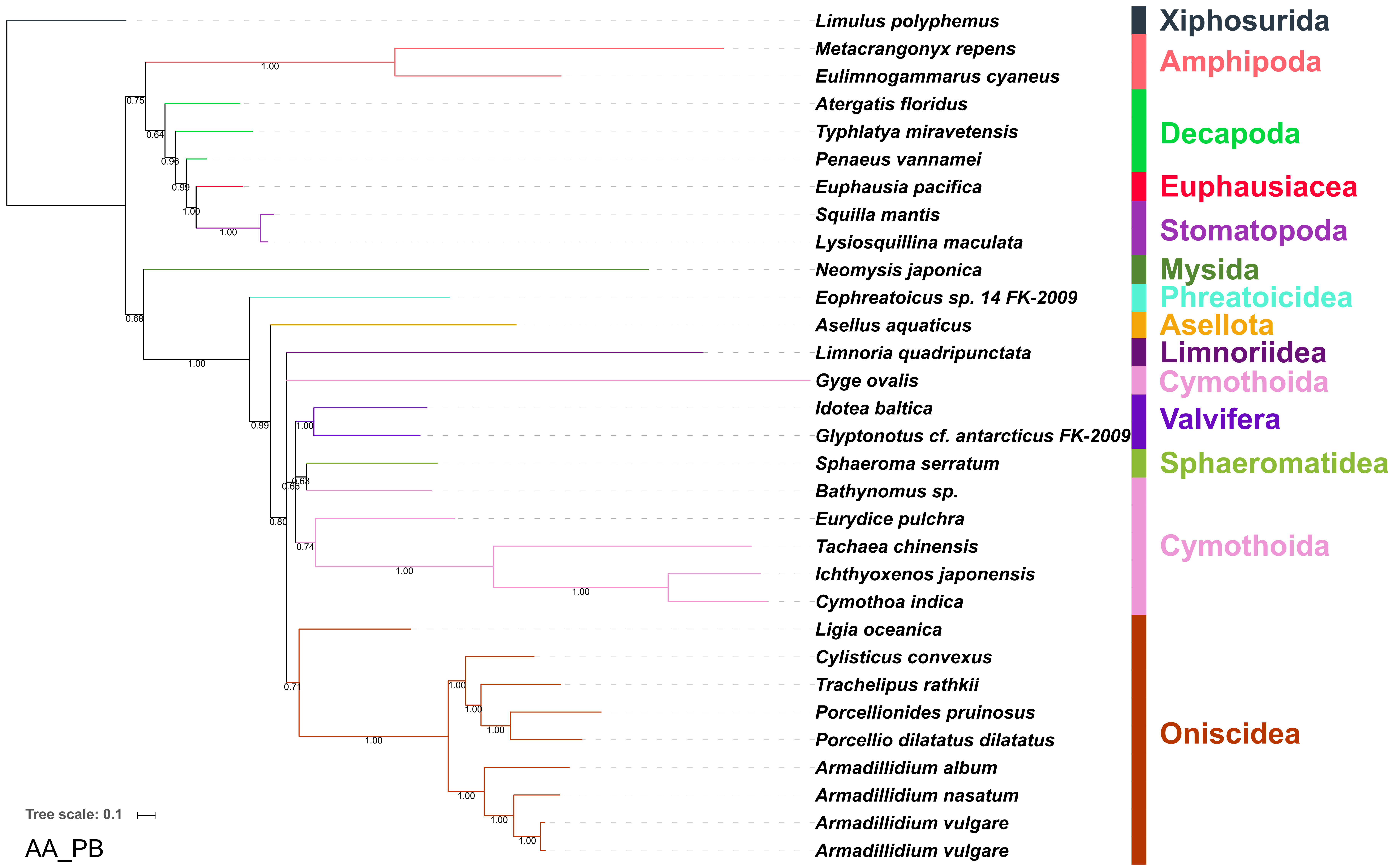

Limulus polyphemus
Metacrangonyx repens
Eulimnogammarus cyaneus
Atergatis floridus
Typhlatya miravetensis
Penaeus vannamei
Euphausia pacifica
Squilla mantis
Lysiosquillina maculata
Neomysis japonica
Eophreatoicus sp. 14 FK-2009
Asellus aquaticus
Limnoria quadripunctata
Gyge ovalis
Idotea baltica
Glyptonotus cf. antarcticus FK-2009
Sphaeroma serratum
Bathynomus sp.
Eurydice pulchra
Tachaea chinensis
Ichthyoxenos japonensis
Cymothoa indica
Ligia oceanica
Cylisticus convexus
Trachelipus rathkii
Porcellionides pruinosus
Porcellio dilatatus dilatatus
Armadillidium album
Armadillidium nasatum
Armadillidium vulgare
Armadillidium vulgare
Xiphosurida
Amphipoda
Decapoda
Euphausiacea
Stomatopoda
Mysida
Phreatoicidea
Asellota
Limnoriidea
Cymothoida
Valvifera
Sphaeromatidea
Cymothoida
Oniscidea
Tree scale: 0.1
AA_PB

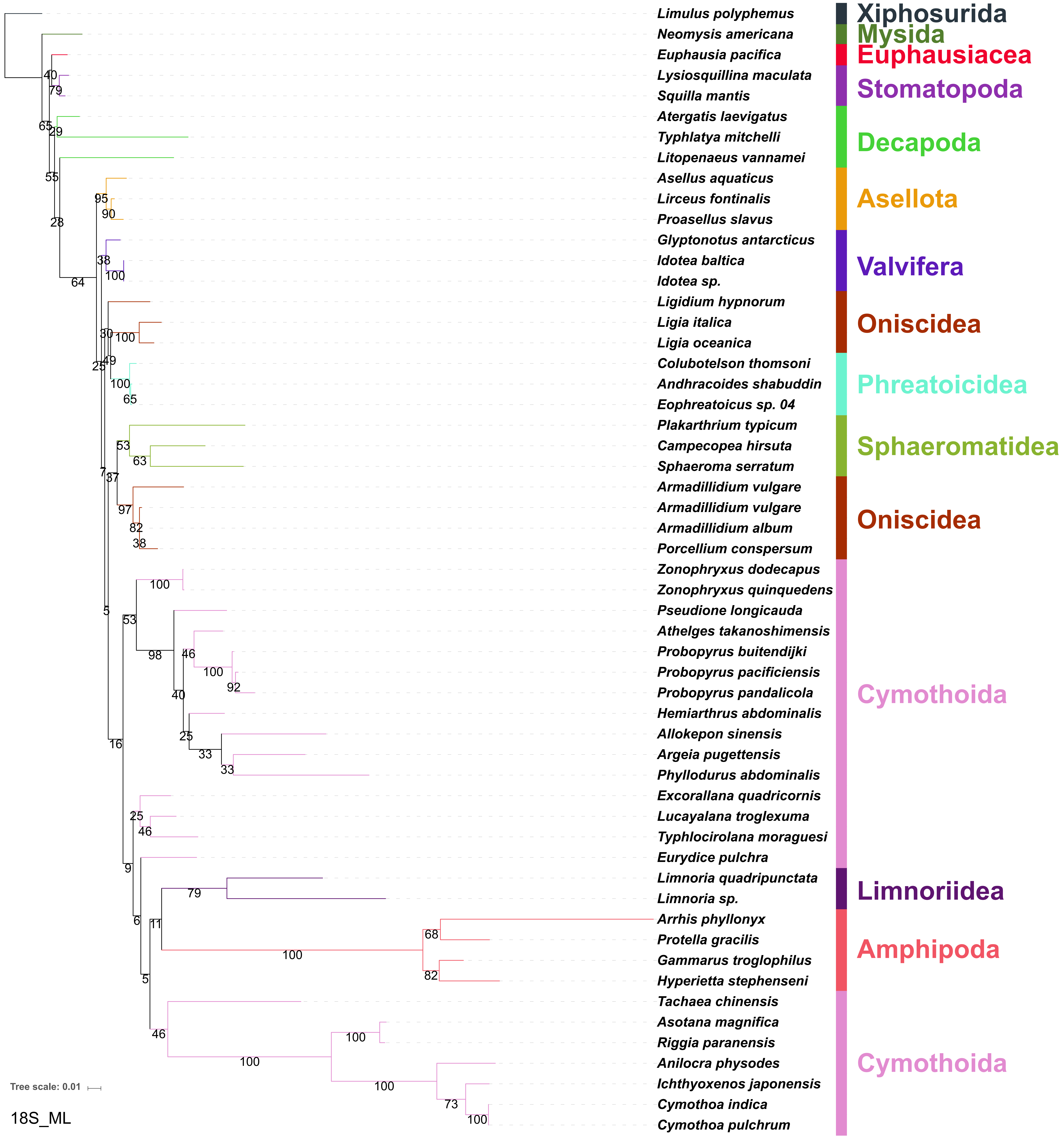

Limulus polyphemus
Neomysis americana
Euphausia pacifica
Lysiosquillina maculata
Squilla mantis
Atergatis laevigatus
Typhlatya mitchelli
Litopenaeus vannamei
Asellus aquaticus
Lirceus fontinalis
Proasellus slavus
Glyptonotus antarcticus
Idotea baltica
Idotea sp.
Ligidium hypnorum
Ligia italica
Ligia oceanica
Colubotelson thomsoni
Andhracoides shabuddin
Eophreatoicus sp. 04
Plakarthrium typicum
Campecopea hirsuta
Sphaeroma serratum
Armadillidium vulgare
Armadillidium vulgare
Armadillidium album
Porcellium conspersum
Zonophryxus dodecapus
Zonophryxus quinquedens
Pseudione longicauda
Athelges takanoshimensis
Probopyrus buitendijki
Probopyrus pacificiensis
Probopyrus pandalicola
Hemiarthrus abdominalis
Allokepon sinensis
Argeia pugettensis
Phyllodurus abdominalis
Excorallana quadricornis
Lucayalana troglexuma
Typhlocirolana moraguesi
Eurydice pulchra
Limnoria quadripunctata
Limnoria sp.
Arrhis phyllonyx
Protella gracilis
Gammarus troglophilus
Hyperietta stephenseni
Tachaea chinensis
Asotana magnifica
Riggia paranensis
Anilocra physodes
Ichthyoxenos japonensis
Cymothoa indica
Cymothoa pulchrum
Xiphosurida
Mysida
Euphausiacea
Stomatopoda
Decapoda
Asellota
Valvifera
Oniscidea
Phreatoicidea
Sphaeromatidea
Oniscidea
Cymothoida
Limnoriidea
Amphipoda
Cymothoida
Tree scale: 0.01
18S_ML

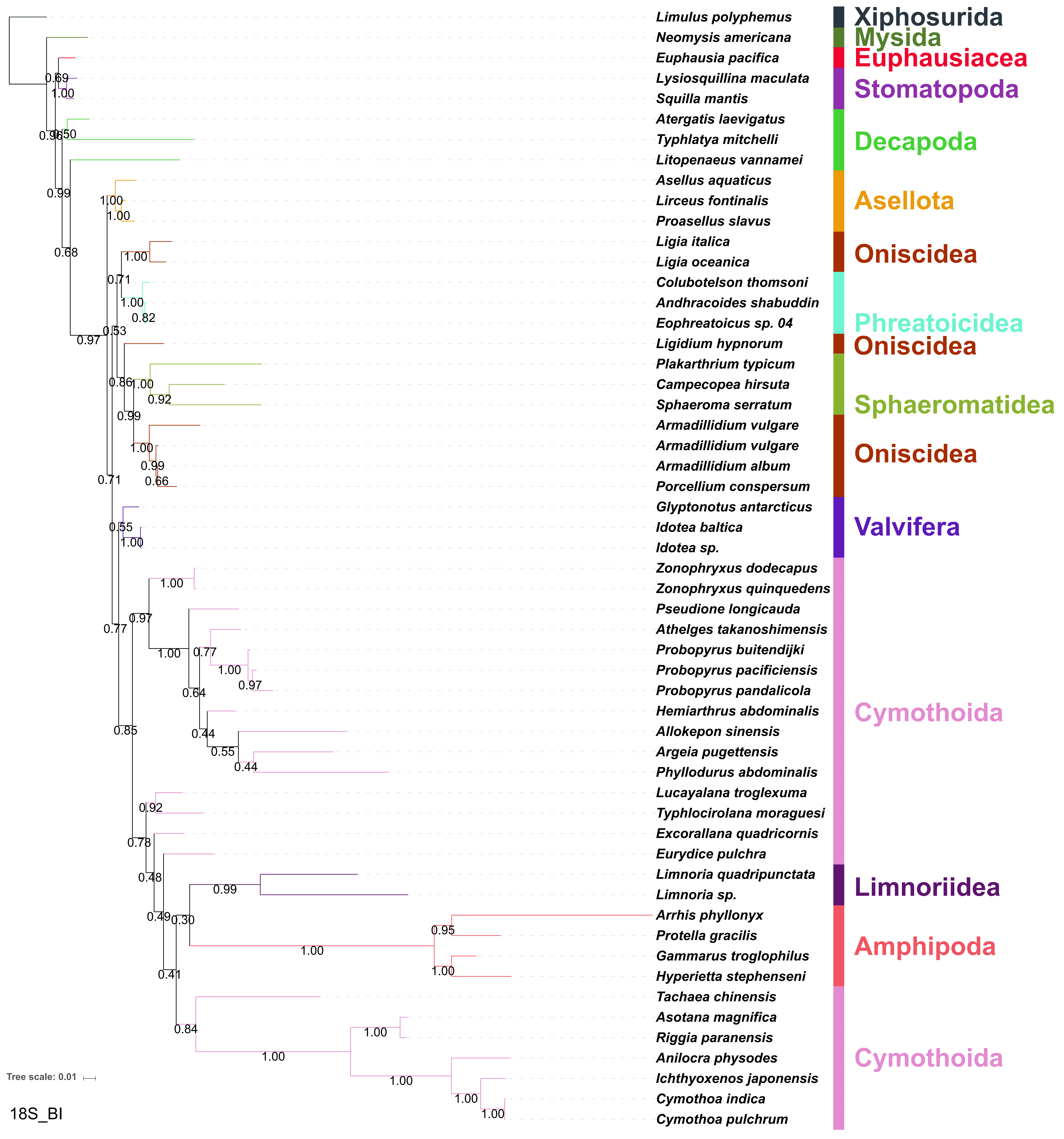

Limulus polyphemus
Neomysis americana
Euphausia pacifica
Lysiosquillina maculata
Squilla mantis
Atergatis laevigatus
Typhlatya mitchelli
Litopenaeus vannamei
Asellus aquaticus
Lirceus fontinalis
Proasellus slavus
Ligia italica
Ligia oceanica
Colubotelson thomsoni
Andhracoides shabuddin
Eophreatoicus sp. 04
Ligidium hypnorum
Plakarthrium typicum
Campecopea hirsuta
Sphaeroma serratum
Armadillidium vulgare
Armadillidium vulgare
Armadillidium album
Porcellium conspersum
Glyptonotus antarcticus
Idotea baltica
Idotea sp.
Zonophryxus dodecapus
Zonophryxus quinquedens
Pseudione longicauda
Athelges takanoshimensis
Probopyrus buitendijki
Probopyrus pacificiensis
Probopyrus pandalicola
Hemiarthrus abdominalis
Allokepon sinensis
Argeia pugettensis
Phyllodurus abdominalis
Lucayalana troglexuma
Typhlocirolana moraguesi
Excorallana quadricornis
Eurydice pulchra
Limnoria quadripunctata
Limnoria sp.
Arrhis phyllonyx
Protella gracilis
Gammarus troglophilus
Hyperietta stephenseni
Tachaea chinensis
Asotana magnifica
Riggia paranensis
Anilocra physodes
Ichthyoxenos japonensis
Cymothoa indica
Cymothoa pulchrum
Xiphosurida
Mysida
Euphausiacea
Stomatopoda
Decapoda
Asellota
Oniscidea
Phreatoicidea
Oniscidea
Sphaeromatidea
Oniscidea
Valvifera
Cymothoida
Limnoriidea
Amphipoda
Cymothoida
Tree scale: 0.01
18S_BI

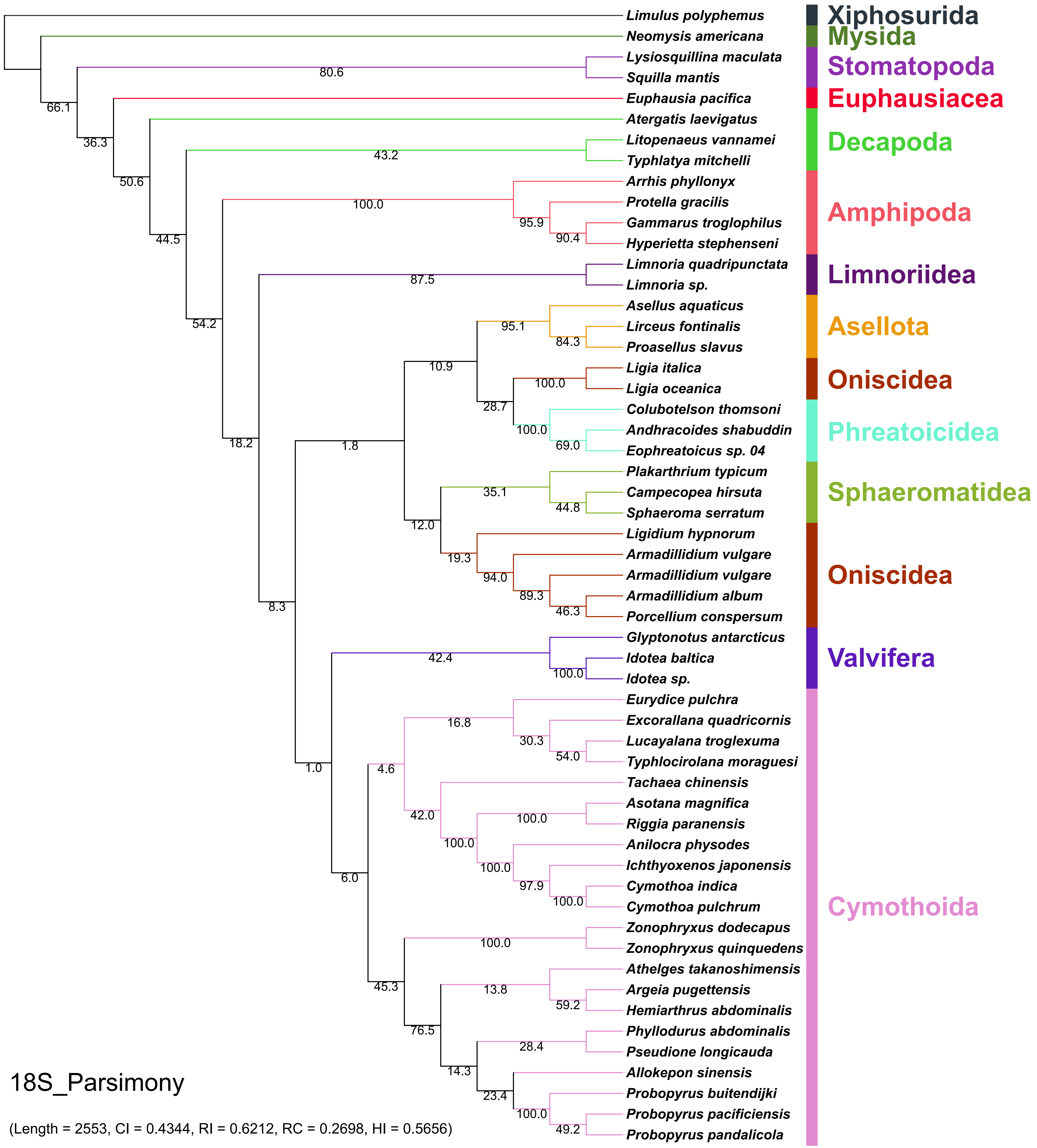

Limulus polyphemus
Neomysis americana
Lysiosquillina maculata
Squilla mantis
Euphausia pacifica
Atergatis laevigatus
Litopenaeus vannamei
Typhlatya mitchelli
Arrhis phyllonyx
Protella gracilis
Gammarus troglophilus
Hyperietta stephenseni
Limnoria quadripunctata
Limnoria sp.
Asellus aquaticus
Lirceus fontinalis
Proasellus slavus
Ligia italica
Ligia oceanica
Colubotelson thomsoni
Andhracoides shabuddin
Eophreatoicus sp. 04
Plakarthrium typicum
Campecopea hirsuta
Sphaeroma serratum
Ligidium hypnorum
Armadillidium vulgare
Armadillidium vulgare
Armadillidium album
Porcellium conspersum
Glyptonotus antarcticus
Idotea baltica
Idotea sp.
Eurydice pulchra
Excorallana quadricornis
Lucayalana troglexuma
Typhlocirolana moraguesi
Tachaea chinensis
Asotana magnifica
Riggia paranensis
Anilocra physodes
Ichthyoxenos japonensis
Cymothoa indica
Cymothoa pulchrum
Zonophryxus dodecapus
Zonophryxus quinquedens
Athelges takanoshimensis
Argeia pugettensis
Hemiarthrus abdominalis
Phyllodurus abdominalis
Pseudione longicauda
Allokepon sinensis
Probopyrus buitendijki
Probopyrus pacificiensis
Probopyrus pandalicola
Xiphosurida
Mysida
Stomatopoda
Euphausiacea
Decapoda
Amphipoda
Limnoriidea
Asellota
Oniscidea
Phreatoicidea
Sphaeromatidea
Oniscidea
Valvifera
Cymothoida
80.6
66.1
36.3
50.6
43.2
44.5
100.0
95.9
90.4
87.5
54.2
95.1
84.3
10.9
100.0
28.7
100.0
69.0
18.2
1.8
35.1
44.8
12.0
19.3
94.0
89.3
46.3
8.3
42.4
100.0
16.8
30.3
54.0
1.0
4.6
42.0
100.0
100.0
100.0
97.9
100.0
6.0
100.0
45.3
13.8
59.2
76.5
28.4
14.3
23.4
100.0
49.2
18S_Parsimony
(Length = 2553, CI = 0.4344, RI = 0.6212, RC = 0.2698, HI = 0.5656)

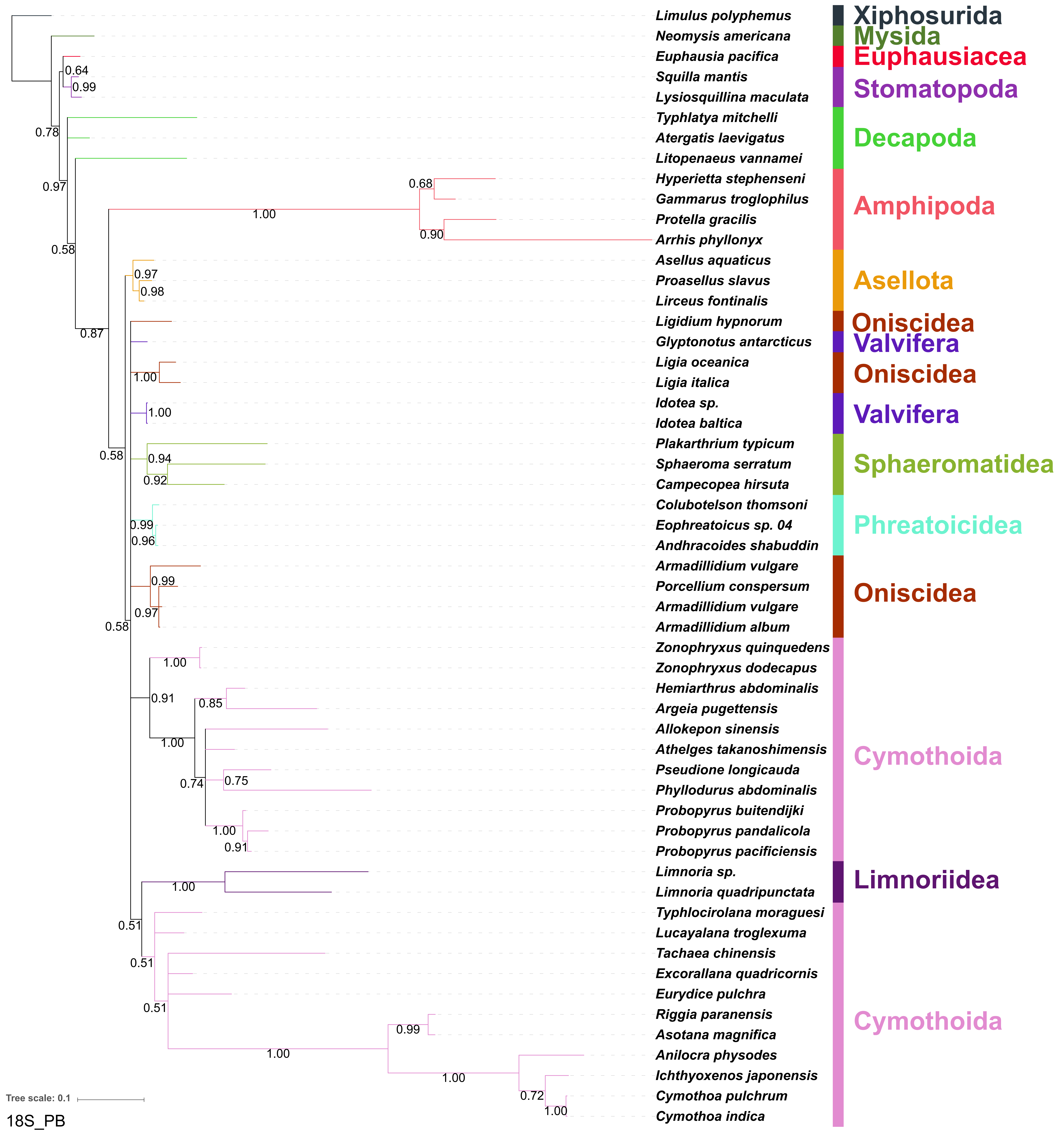

Limulus polyphemus
Neomysis americana
Euphausia pacifica
Squilla mantis
Lysiosquillina maculata
Typhlatya mitchelli
Atergatis laevigatus
Litopenaeus vannamei
Hyperietta stephenseni
Gammarus troglophilus
Protella gracilis
Arrhis phyllonyx
Asellus aquaticus
Proasellus slavus
Lirceus fontinalis
Ligidium hypnorum
Glyptonotus antarcticus
Ligia oceanica
Ligia italica
Idotea sp.
Idotea baltica
Plakarthrium typicum
Sphaeroma serratum
Campecopea hirsuta
Colubotelson thomsoni
Eophreatoicus sp. 04
Andhracoides shabuddin
Armadillidium vulgare
Porcellium conspersum
Armadillidium vulgare
Armadillidium album
Zonophryxus quinquedens
Zonophryxus dodecapus
Hemiarthrus abdominalis
Argeia pugettensis
Allokepon sinensis
Athelges takanoshimensis
Pseudione longicauda
Phyllodurus abdominalis
Probopyrus buitendijki
Probopyrus pandalicola
Probopyrus pacificiensis
Limnoria sp.
Limnoria quadripunctata
Typhlocirolana moraguesi
Lucayalana troglexuma
Tachaea chinensis
Excorallana quadricornis
Eurydice pulchra
Riggia paranensis
Asotana magnifica
Anilocra physodes
Ichthyoxenos japonensis
Cymothoa pulchrum
Cymothoa indica
Xiphosurida
Mysida
Euphausiacea
Stomatopoda
Decapoda
Amphipoda
Asellota
Oniscidea
Valvifera
Oniscidea
Valvifera
Sphaeromatidea
Phreatoicidea
Oniscidea
Cymothoida
Limnoriidea
Cymothoida
0.64
0.99
0.78
0.97
0.58
0.87
1.00
0.68
0.90
0.97
0.98
0.58
1.00
1.00
0.94
0.92
0.99
0.96
0.99
0.97
0.58
1.00
0.91
0.85
1.00
0.74
0.75
1.00
0.91
1.00
0.51
0.51
0.51
0.99
1.00
1.00
0.72
1.00
Tree scale: 0.1
18S_PB

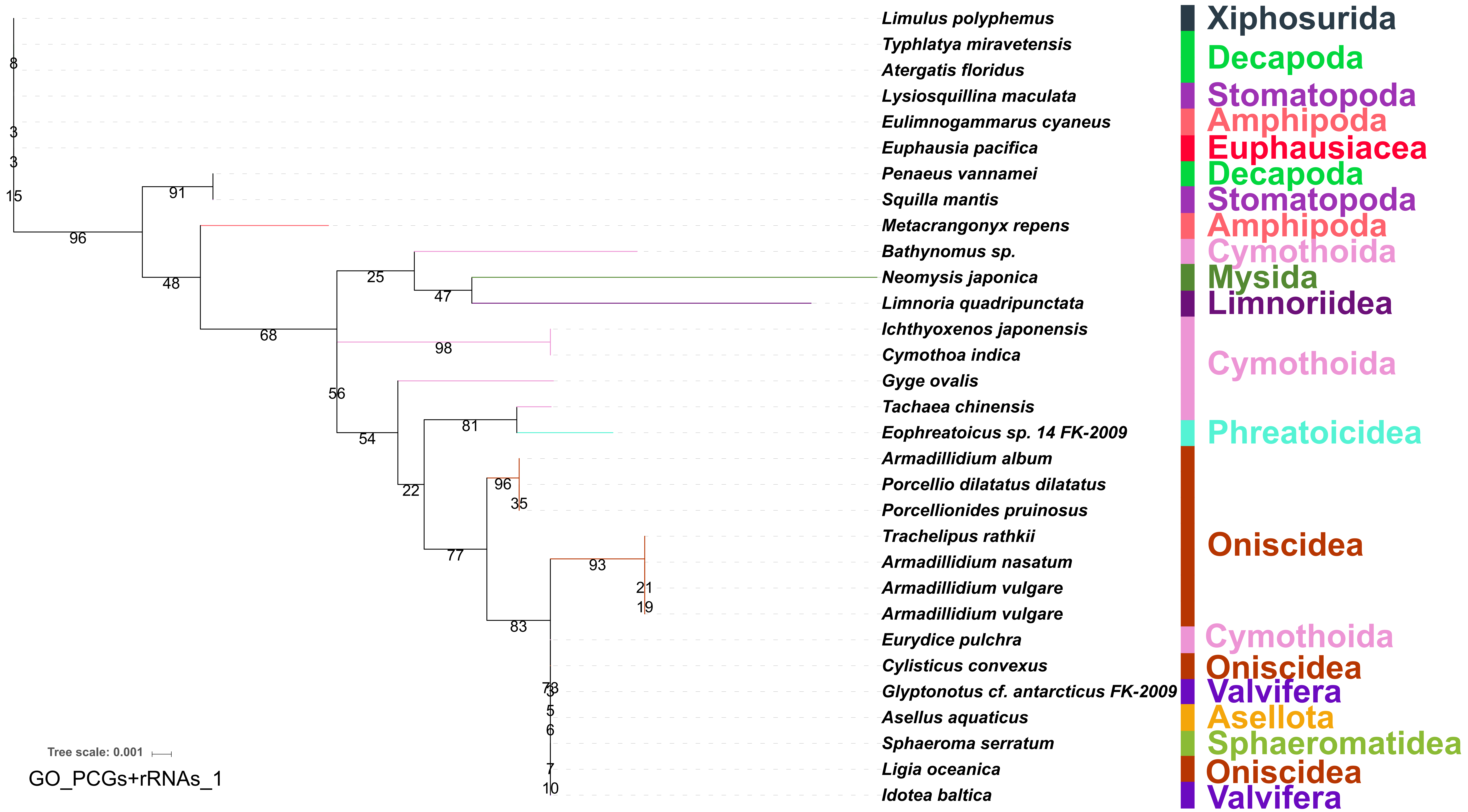
Limulus polyphemus
Typhlatya miravetensis
Atergatis floridus
Lysiosquillina maculata
Eulimnogammarus cyaneus
Euphausia pacifica
Penaeus vannamei
Squilla mantis
Metacrangonyx repens
Bathynomus sp.
Neomysis japonica
Limnoria quadripunctata
Ichthyoxenos japonensis
Cymothoa indica
Gyge ovalis
Tachaea chinensis
Eophreatoicus sp. 14 FK-2009
Armadillidium album
Porcellio dilatatus dilatatus
Porcellionides pruinosus
Trachelipus rathkii
Armadillidium nasatum
Armadillidium vulgare
Armadillidium vulgare
Eurydice pulchra
Cylisticus convexus
Glyptonotus cf. antarcticus FK-2009
Asellus aquaticus
Sphaeroma serratum
Ligia oceanica
Idotea baltica
Xiphosurida
Decapoda
Stomatopoda
Amphipoda
Euphausiacea
Decapoda
Stomatopoda
Amphipoda
Cymothoida
Mysida
Limnoriidea
Cymothoida
Phreatoicidea
Oniscidea
Cymothoida
Oniscidea
Valvifera
Asellota
Sphaeromatidea
Oniscidea
Valvifera
Tree scale: 0.001
GO_PCGs+rRNAs_1

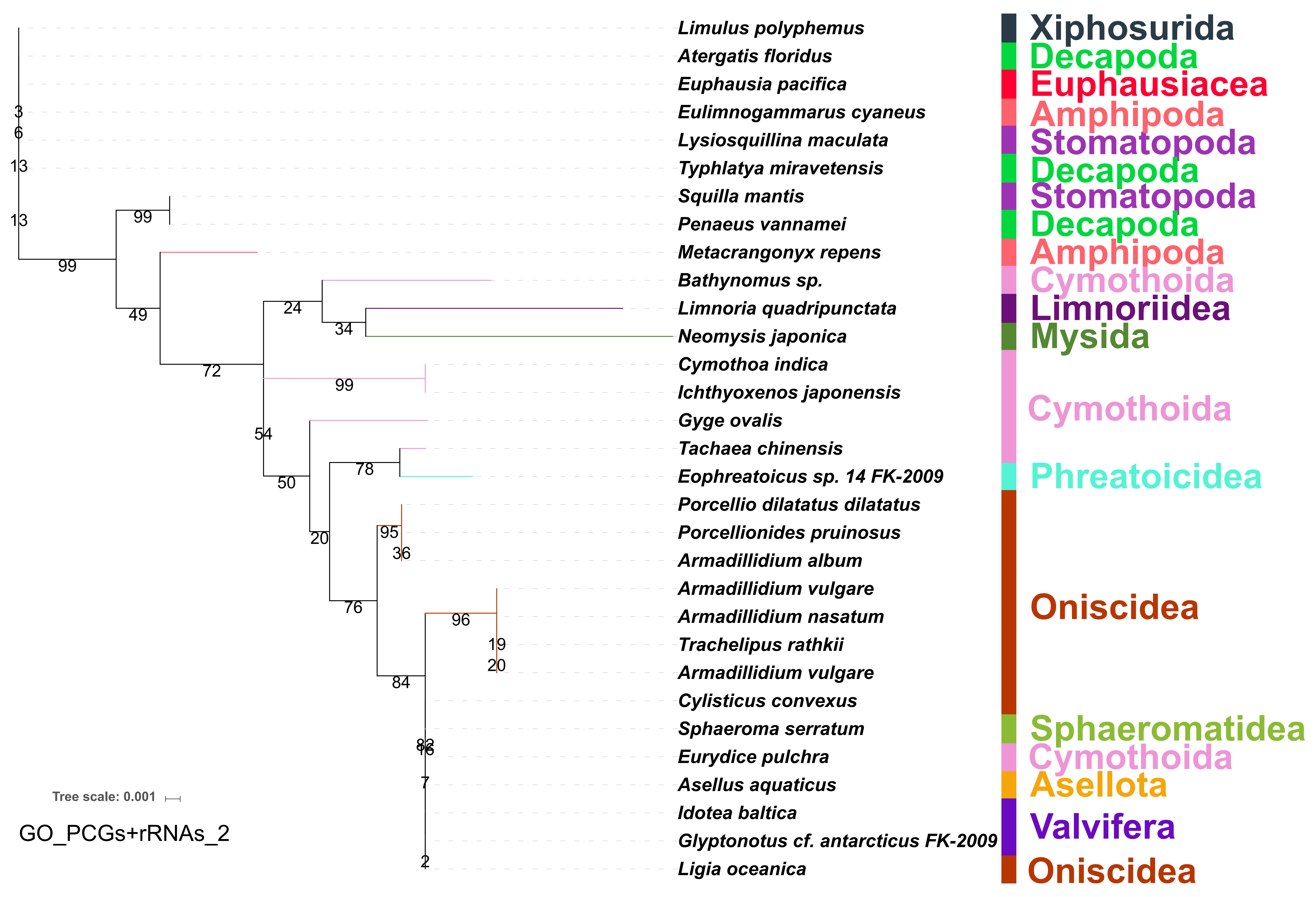

Limulus polyphemus
Atergatis floridus
Euphausia pacifica
Eulimnogammarus cyaneus
Lysiosquillina maculata
Typhlatya miravetensis
Squilla mantis
Penaeus vannamei
Metacrangonyx repens
Bathynomus sp.
Limnoria quadripunctata
Neomysis japonica
Cymothoa indica
Ichthyoxenos japonensis
Gyge ovalis
Tachaea chinensis
Eophreatoicus sp. 14 FK-2009
Porcellio dilatatus dilatatus
Porcellionides pruinosus
Armadillidium album
Armadillidium vulgare
Armadillidium nasatum
Trachelipus rathkii
Armadillidium vulgare
Cylisticus convexus
Sphaeroma serratum
Eurydice pulchra
Asellus aquaticus
Idotea baltica
Glyptonotus cf. antarcticus FK-2009
Ligia oceanica
Xiphosurida
Decapoda
Euphausiacea
Amphipoda
Stomatopoda
Decapoda
Stomatopoda
Decapoda
Amphipoda
Cymothoida
Limnoriidea
Mysida
Cymothoida
Phreatoicidea
Oniscidea
Sphaeromatidea
Cymothoida
Asellota
Valvifera
Oniscidea
Tree scale: 0.001
GO_PCGs+rRNAs_2

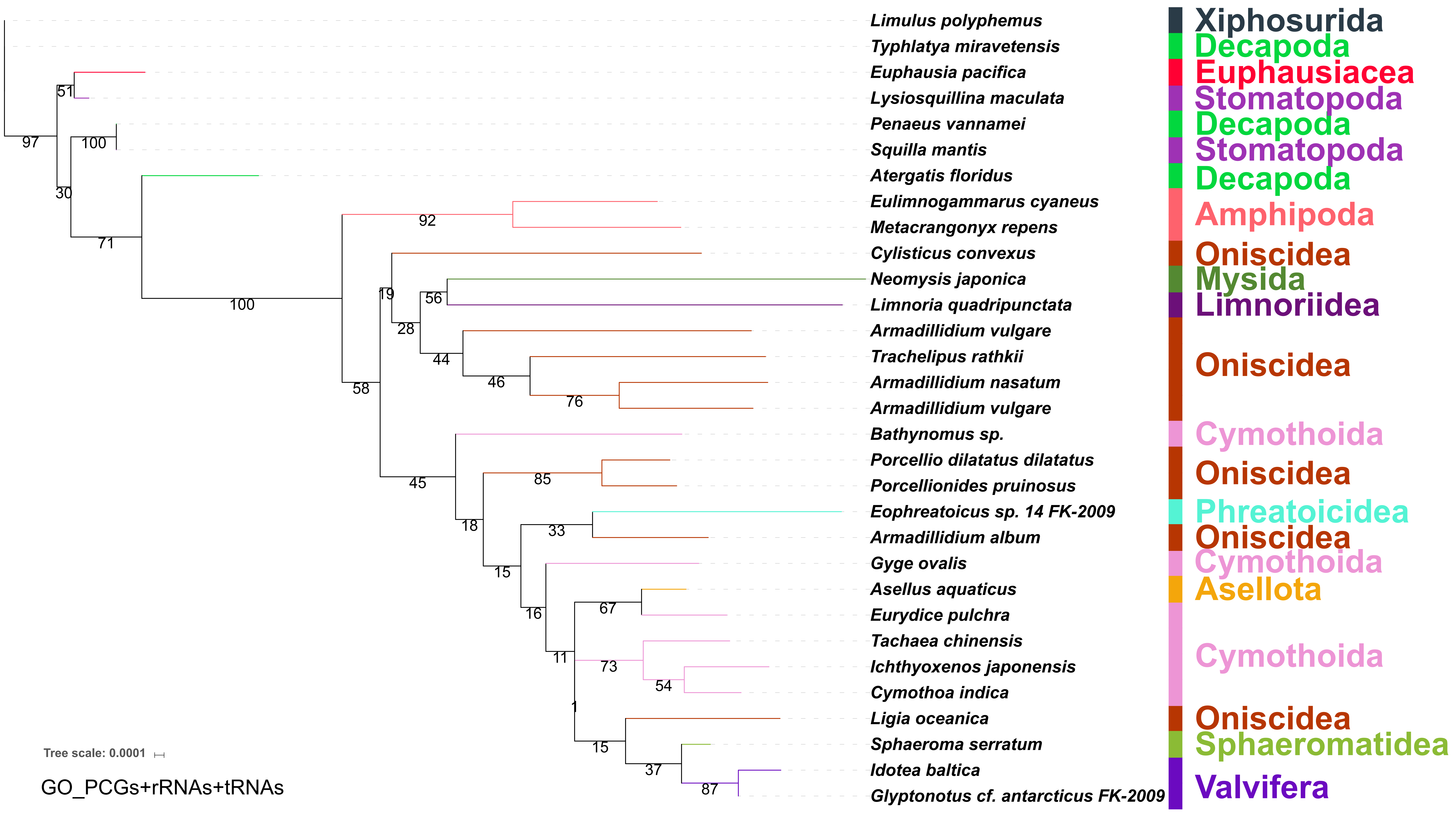

Limulus polyphemus
Typhlatya miravetensis
Euphausia pacifica
Lysiosquillina maculata
Penaeus vannamei
Squilla mantis
Atergatis floridus
Eulimnogammarus cyaneus
Metacrangonyx repens
Cylisticus convexus
Neomysis japonica
Limnoria quadripunctata
Armadillidium vulgare
Trachelipus rathkii
Armadillidium nasatum
Armadillidium vulgare
Bathynomus sp.
Porcellio dilatatus dilatatus
Porcellionides pruinosus
Eophreatoicus sp. 14 FK-2009
Armadillidium album
Gyge ovalis
Asellus aquaticus
Eurydice pulchra
Tachaea chinensis
Ichthyoxenos japonensis
Cymothoa indica
Ligia oceanica
Sphaeroma serratum
Idotea baltica
Glyptonotus cf. antarcticus FK-2009
Xiphosurida
Decapoda
Euphausiacea
Stomatopoda
Decapoda
Stomatopoda
Decapoda
Amphipoda
Oniscidea
Mysida
Limnoriidea
Oniscidea
Cymothoida
Oniscidea
Phreatoicidea
Oniscidea
Cymothoida
Asellota
Cymothoida
Oniscidea
Sphaeromatidea
Valvifera
97
51
100
30
71
100
92
19
56
28
44
46
76
58
45
85
18
33
15
16
67
11
73
54
1
15
37
87
Tree scale: 0.0001
GO_PCGs+rRNAs+tRNAs

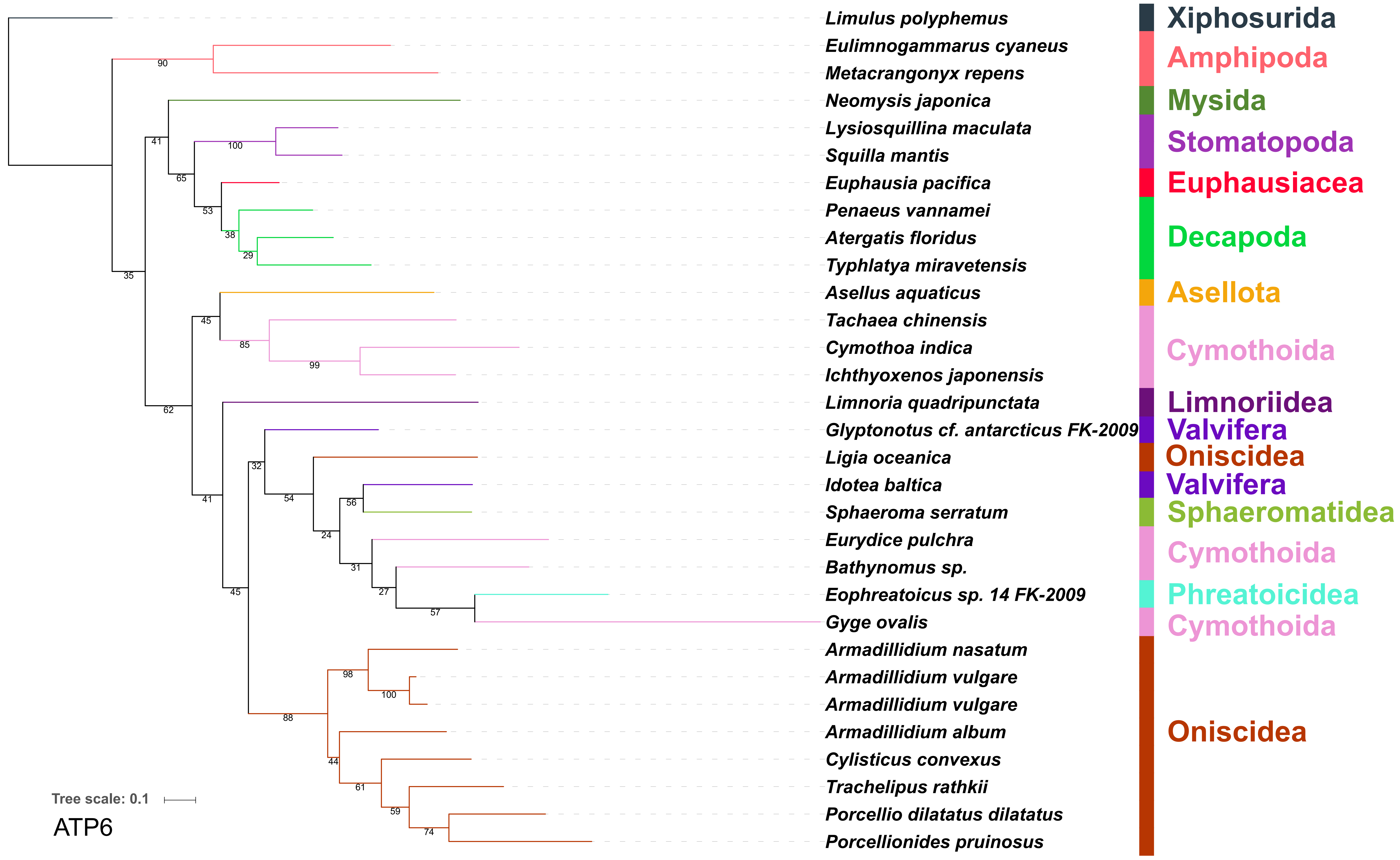

Limulus polyphemus
Eulimnogammarus cyaneus
Metacrangonyx repens
Neomysis japonica
Lysiosquillina maculata
Squilla mantis
Euphausia pacifica
Penaeus vannamei
Atergatis floridus
Typhlatya miravetensis
Asellus aquaticus
Tachaea chinensis
Cymothoa indica
Ichthyoxenos japonensis
Limnoria quadripunctata
Glyptonotus cf. antarcticus FK-2009
Ligia oceanica
Idotea baltica
Sphaeroma serratum
Eurydice pulchra
Bathynomus sp.
Eophreatoicus sp. 14 FK-2009
Gyge ovalis
Armadillidium nasatum
Armadillidium vulgare
Armadillidium vulgare
Armadillidium album
Cylisticus convexus
Trachelipus rathkii
Porcellio dilatatus dilatatus
Porcellionides pruinosus
Xiphosurida
Amphipoda
Mysida
Stomatopoda
Euphausiacea
Decapoda
Asellota
Cymothoida
Limnoriidea
Valvifera
Oniscidea
Valvifera
Sphaeromatidea
Cymothoida
Phreatoicidea
Cymothoida
Oniscidea
Tree scale: 0.1
ATP6

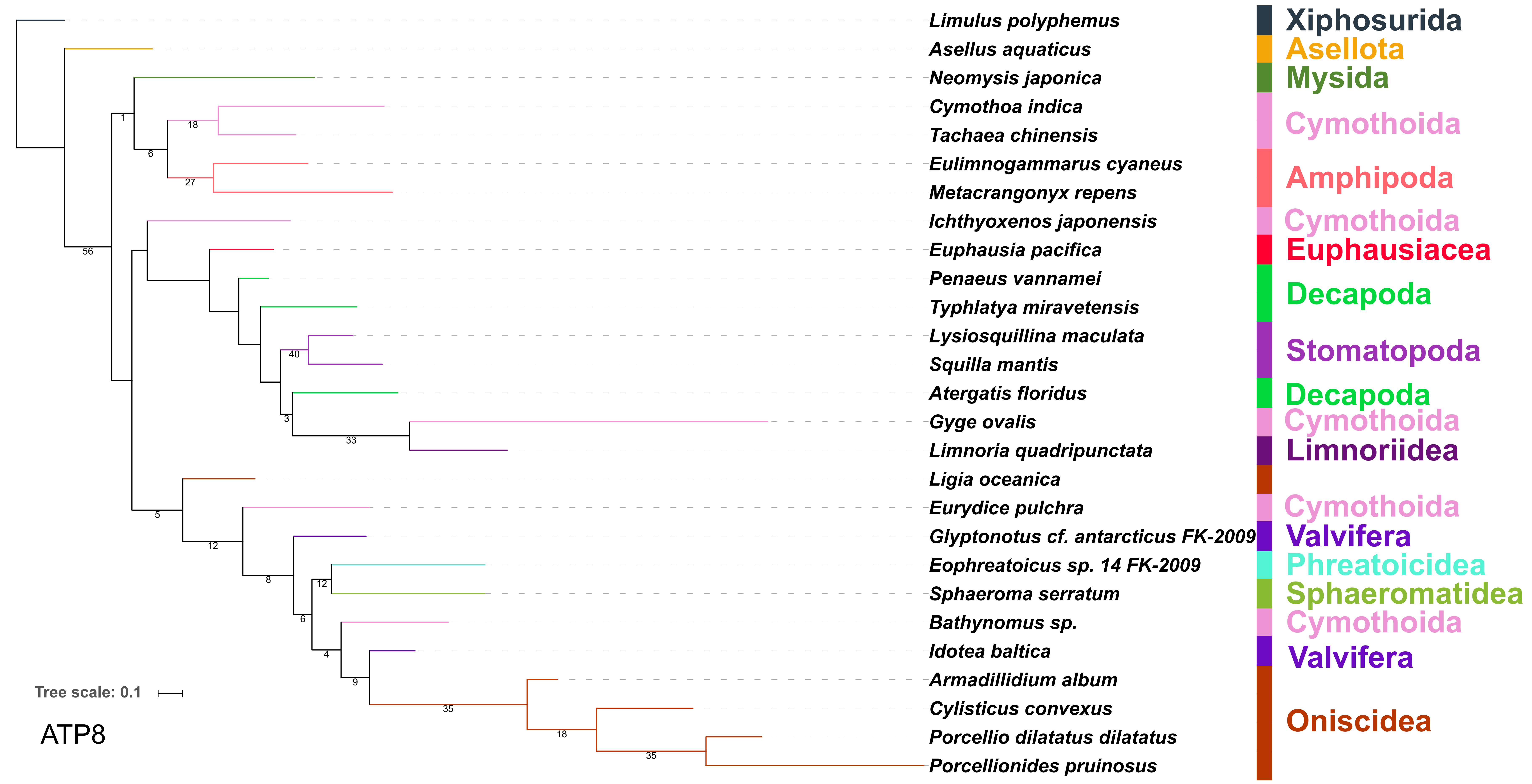
Limulus polyphemus
Asellus aquaticus
Neomysis japonica
Cymothoa indica
Tachaea chinensis
Eulimnogammarus cyaneus
Metacrangonyx repens
Ichthyoxenos japonensis
Euphausia pacifica
Penaeus vannamei
Typhlatya miravetensis
Lysiosquillina maculata
Squilla mantis
Atergatis floridus
Gyge ovalis
Limnoria quadripunctata
Ligia oceanica
Eurydice pulchra
Glyptonotus cf. antarcticus FK-2009
Eophreatoicus sp. 14 FK-2009
Sphaeroma serratum
Bathynomus sp.
Idotea baltica
Armadillidium album
Cylisticus convexus
Porcellio dilatatus dilatatus
Porcellionides pruinosus
Xiphosurida
Asellota
Mysida
Cymothoida
Amphipoda
Cymothoida
Euphausiacea
Decapoda
Stomatopoda
Decapoda
Cymothoida
Limnoriidea
Cymothoida
Valvifera
Phreatoicidea
Sphaeromatidea
Cymothoida
Valvifera
Oniscidea
Tree scale: 0.1
ATP8

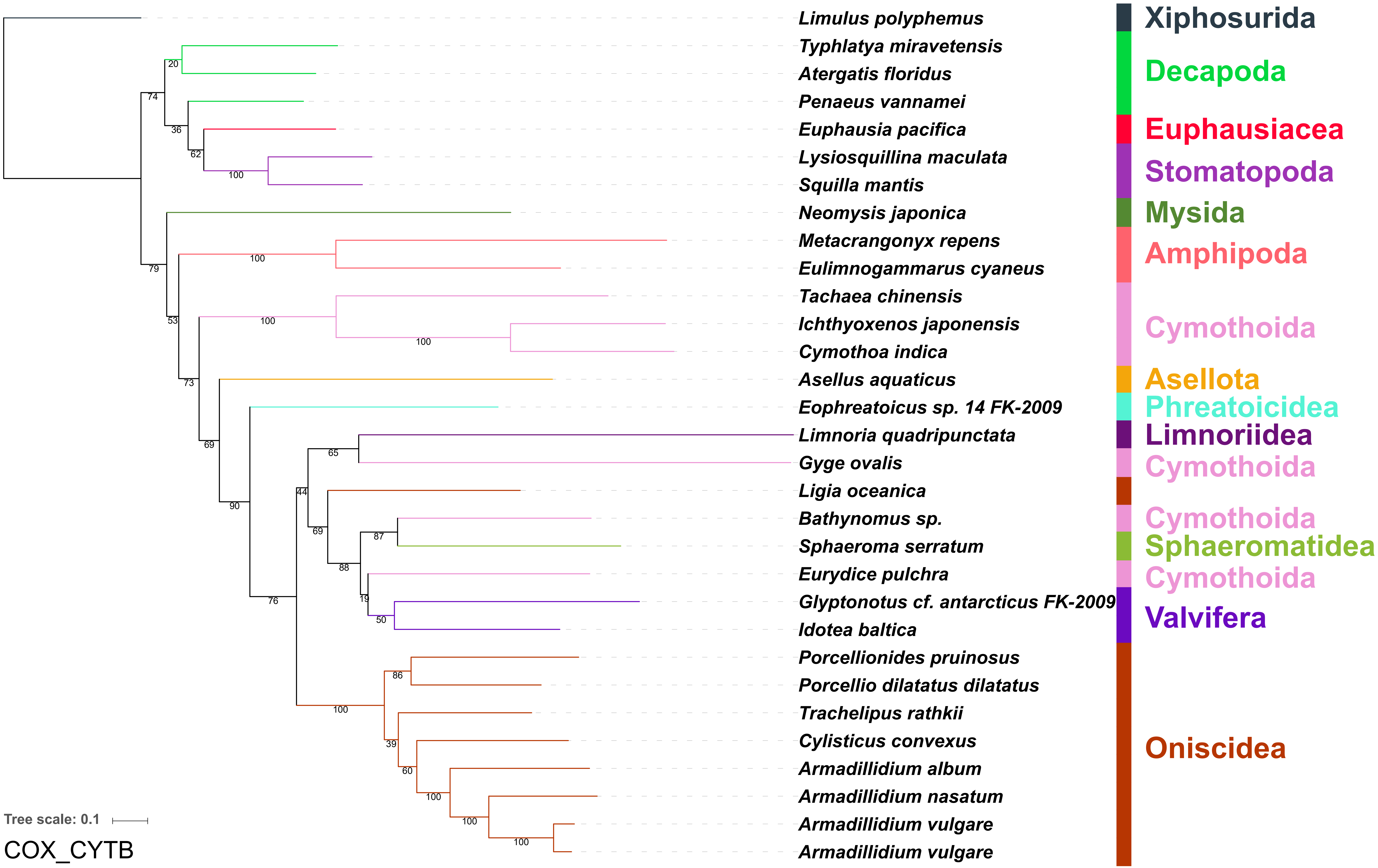

Limulus polyphemus
Typhlatya miravetensis
Atergatis floridus
Penaeus vannamei
Euphausia pacifica
Lysiosquillina maculata
Squilla mantis
Neomysis japonica
Metacrangonyx repens
Eulimnogammarus cyaneus
Tachaea chinensis
Ichthyoxenos japonensis
Cymothoa indica
Asellus aquaticus
Eophreatoicus sp. 14 FK-2009
Limnoria quadripunctata
Gyge ovalis
Ligia oceanica
Bathynomus sp.
Sphaeroma serratum
Eurydice pulchra
Glyptonotus cf. antarcticus FK-2009
Idotea baltica
Porcellionides pruinosus
Porcellio dilatatus dilatatus
Trachelipus rathkii
Cylisticus convexus
Armadillidium album
Armadillidium nasatum
Armadillidium vulgare
Armadillidium vulgare
Xiphosurida
Decapoda
Euphausiacea
Stomatopoda
Mysida
Amphipoda
Cymothoida
Asellota
Phreatoicidea
Limnoriidea
Cymothoida
Cymothoida
Sphaeromatidea
Cymothoida
Valvifera
Oniscidea
Tree scale: 0.1
COX_CYTB

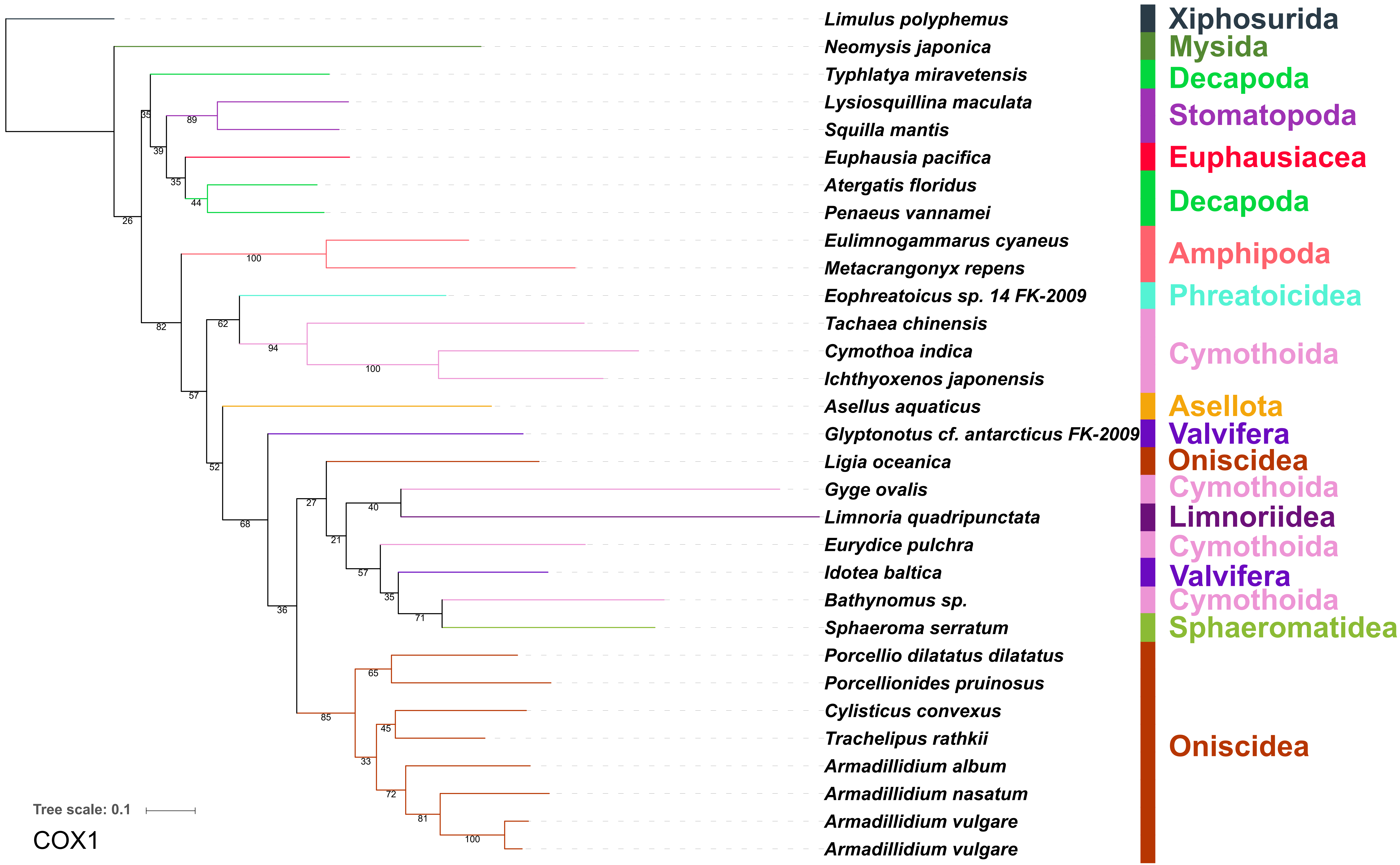

Limulus polyphemus
Neomysis japonica
Typhlatya miravetensis
Lysiosquillina maculata
Squilla mantis
Euphausia pacifica
Atergatis floridus
Penaeus vannamei
Eulimnogammarus cyaneus
Metacrangonyx repens
Eophreatoicus sp. 14 FK-2009
Tachaea chinensis
Cymothoa indica
Ichthyoxenos japonensis
Asellus aquaticus
Glyptonotus cf. antarcticus FK-2009
Ligia oceanica
Gyge ovalis
Limnoria quadripunctata
Eurydice pulchra
Idotea baltica
Bathynomus sp.
Sphaeroma serratum
Porcellio dilatatus dilatatus
Porcellionides pruinosus
Cylisticus convexus
Trachelipus rathkii
Armadillidium album
Armadillidium nasatum
Armadillidium vulgare
Armadillidium vulgare
Xiphosurida
Mysida
Decapoda
Stomatopoda
Euphausiacea
Decapoda
Amphipoda
Phreatoicidea
Cymothoida
Asellota
Valvifera
Oniscidea
Cymothoida
Limnoriidea
Cymothoida
Valvifera
Cymothoida
Sphaeromatidea
Oniscidea
Tree scale: 0.1
COX1

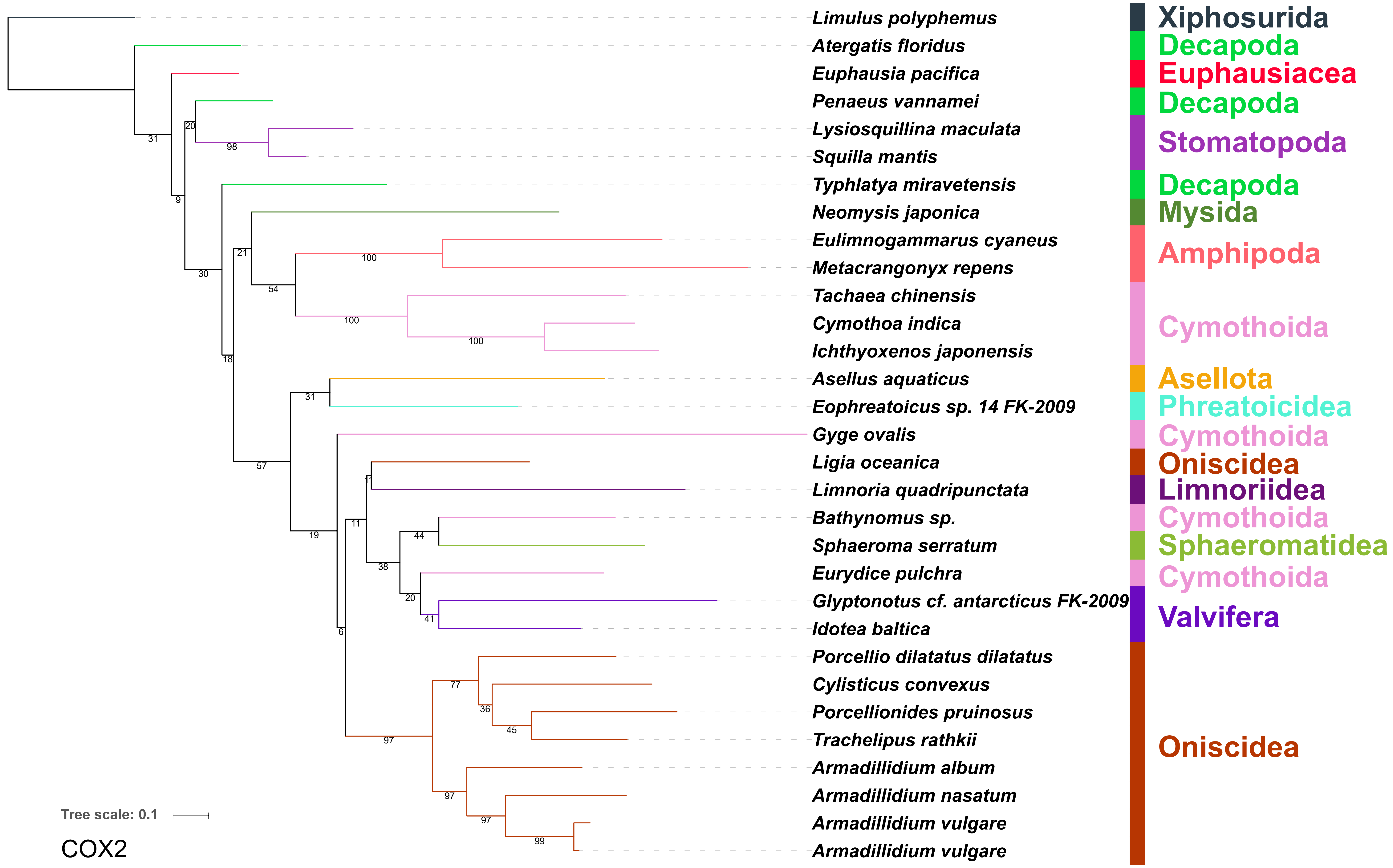

Limulus polyphemus
Atergatis floridus
Euphausia pacifica
Penaeus vannamei
Lysiosquillina maculata
Squilla mantis
Typhlatya miravetensis
Neomysis japonica
Eulimnogammarus cyaneus
Metacrangonyx repens
Tachaea chinensis
Cymothoa indica
Ichthyoxenos japonensis
Asellus aquaticus
Eophreatoicus sp. 14 FK-2009
Gyge ovalis
Ligia oceanica
Limnoria quadripunctata
Bathynomus sp.
Sphaeroma serratum
Eurydice pulchra
Glyptonotus cf. antarcticus FK-2009
Idotea baltica
Porcellio dilatatus dilatatus
Cylisticus convexus
Porcellionides pruinosus
Trachelipus rathkii
Armadillidium album
Armadillidium nasatum
Armadillidium vulgare
Armadillidium vulgare
Xiphosurida
Decapoda
Euphausiacea
Decapoda
Stomatopoda
Decapoda
Mysida
Amphipoda
Cymothoida
Asellota
Phreatoicidea
Cymothoida
Oniscidea
Limnoriidea
Cymothoida
Sphaeromatidea
Cymothoida
Valvifera
Oniscidea
Tree scale: 0.1
COX2

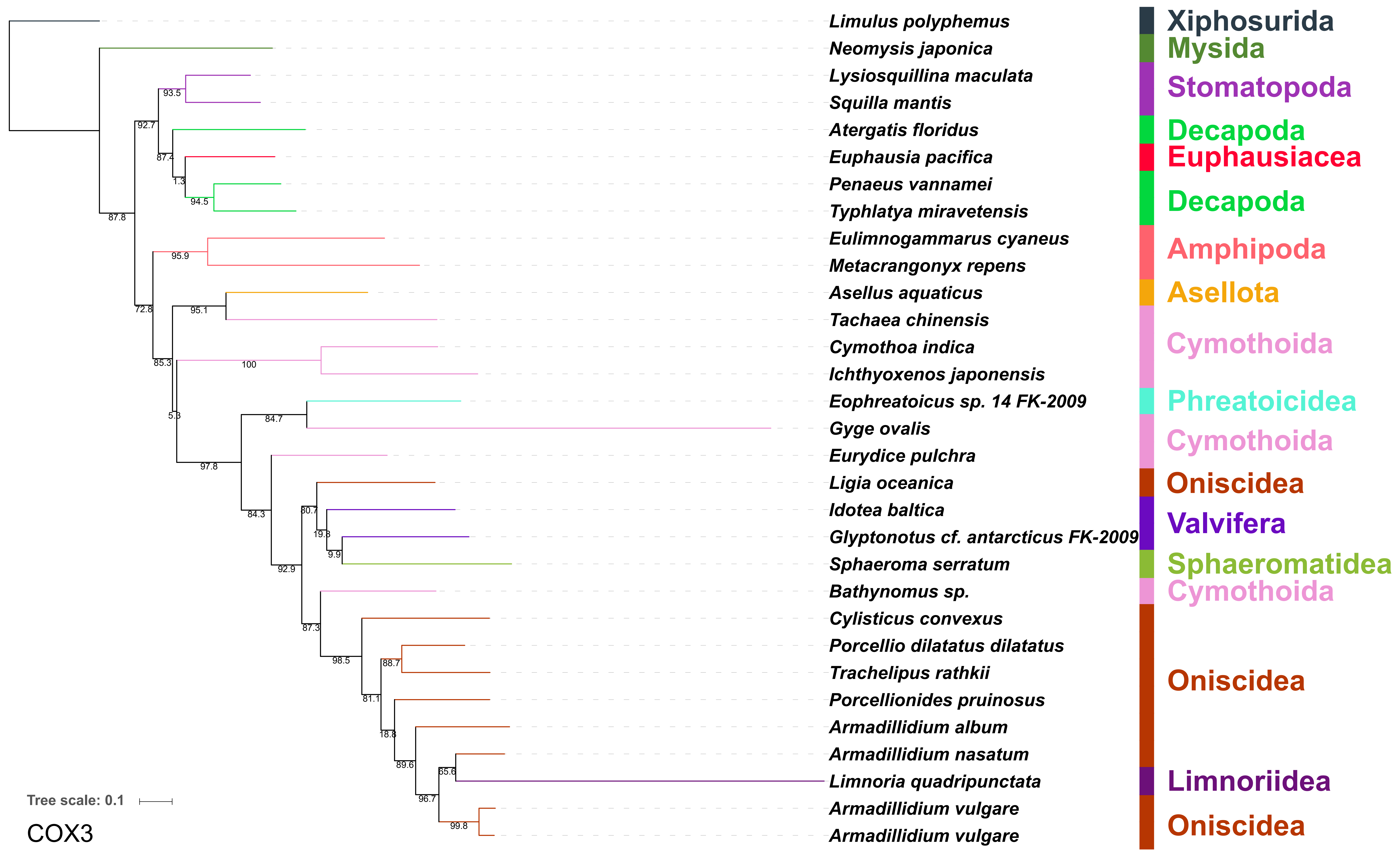
Limulus polyphemus
Neomysis japonica
Lysiosquillina maculata
Squilla mantis
Atergatis floridus
Euphausia pacifica
Penaeus vannamei
Typhlatya miravetensis
Eulimnogammarus cyaneus
Metacrangonyx repens
Asellus aquaticus
Tachaea chinensis
Cymothoa indica
Ichthyoxenos japonensis
Eophreatoicus sp. 14 FK-2009
Gyge ovalis
Eurydice pulchra
Ligia oceanica
Idotea baltica
Glyptonotus cf. antarcticus FK-2009
Sphaeroma serratum
Bathynomus sp.
Cylisticus convexus
Porcellio dilatatus dilatatus
Trachelipus rathkii
Porcellionides pruinosus
Armadillidium album
Armadillidium nasatum
Limnoria quadripunctata
Armadillidium vulgare
Armadillidium vulgare
Xiphosurida
Mysida
Stomatopoda
Decapoda
Euphausiacea
Decapoda
Amphipoda
Asellota
Cymothoida
Phreatoicidea
Cymothoida
Oniscidea
Valvifera
Sphaeromatidea
Cymothoida
Oniscidea
Limnoriidea
Oniscidea
Tree scale: 0.1
COX3

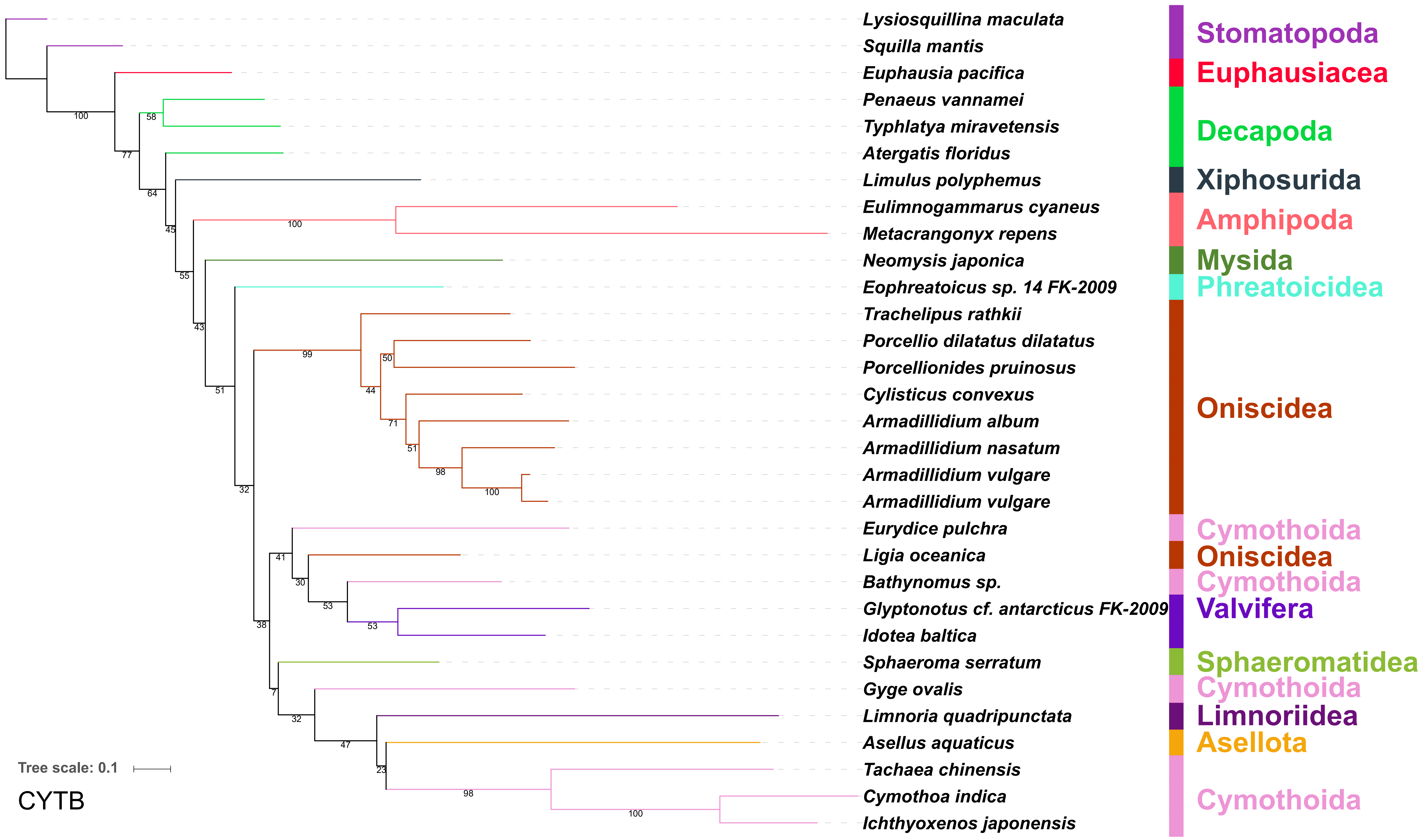
Lysiosquillina maculata
Squilla mantis
Euphausia pacifica
Penaeus vannamei
Typhlatya miravetensis
Atergatis floridus
Limulus polyphemus
Eulimnogammarus cyaneus
Metacrangonyx repens
Neomysis japonica
Eophreatoicus sp. 14 FK-2009
Trachelipus rathkii
Porcellio dilatatus dilatatus
Porcellionides pruinosus
Cylisticus convexus
Armadillidium album
Armadillidium nasatum
Armadillidium vulgare
Armadillidium vulgare
Eurydice pulchra
Ligia oceanica
Bathynomus sp.
Glyptonotus cf. antarcticus FK-2009
Idotea baltica
Sphaeroma serratum
Gyge ovalis
Limnoria quadripunctata
Asellus aquaticus
Tachaea chinensis
Cymothoa indica
Ichthyoxenos japonensis
Stomatopoda
Euphausiacea
Decapoda
Xiphosurida
Amphipoda
Mysida
Phreatoicidea
Oniscidea
Cymothoida
Oniscidea
Cymothoida
Valvifera
Sphaeromatidea
Cymothoida
Limnoriidea
Asellota
Cymothoida
Tree scale: 0.1
CYTB

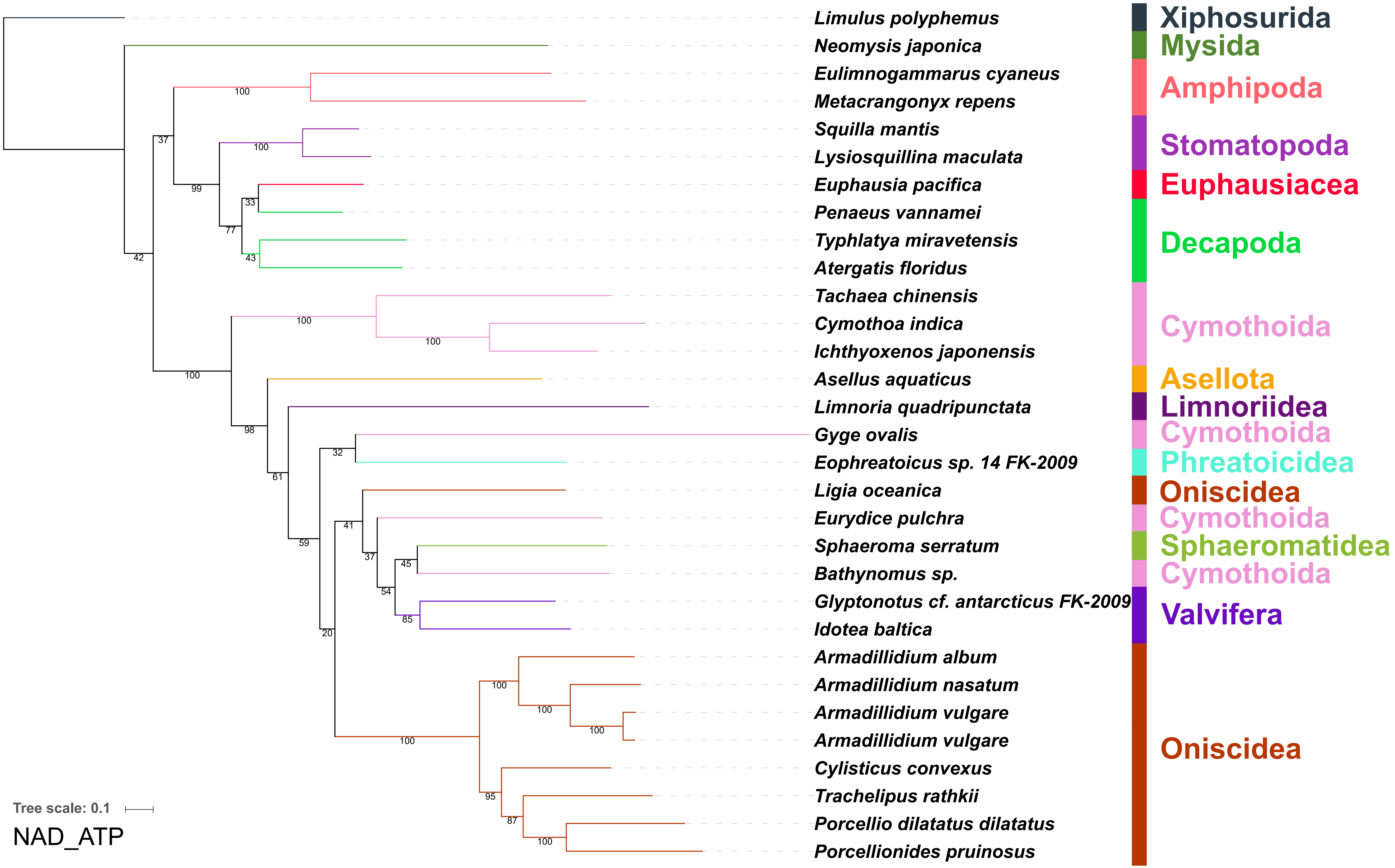
Limulus polyphemus
Neomysis japonica
Eulimnogammarus cyaneus
Metacrangonyx repens
Squilla mantis
Lysiosquillina maculata
Euphausia pacifica
Penaeus vannamei
Typhlatya miravetensis
Atergatis floridus
Tachaea chinensis
Cymothoa indica
Ichthyoxenos japonensis
Asellus aquaticus
Limnoria quadripunctata
Gyge ovalis
Eophreatoicus sp. 14 FK-2009
Ligia oceanica
Eurydice pulchra
Sphaeroma serratum
Bathynomus sp.
Glyptonotus cf. antarcticus FK-2009
Idotea baltica
Armadillidium album
Armadillidium nasatum
Armadillidium vulgare
Armadillidium vulgare
Cylisticus convexus
Trachelipus rathkii
Porcellio dilatatus dilatatus
Porcellionides pruinosus
Xiphosurida
Mysida
Amphipoda
Stomatopoda
Euphausiacea
Decapoda
Cymothoida
Asellota
Limnoriidea
Cymothoida
Phreatoicidea
Oniscidea
Cymothoida
Sphaeromatidea
Cymothoida
Valvifera
Oniscidea
Tree scale: 0.1
NAD_ATP

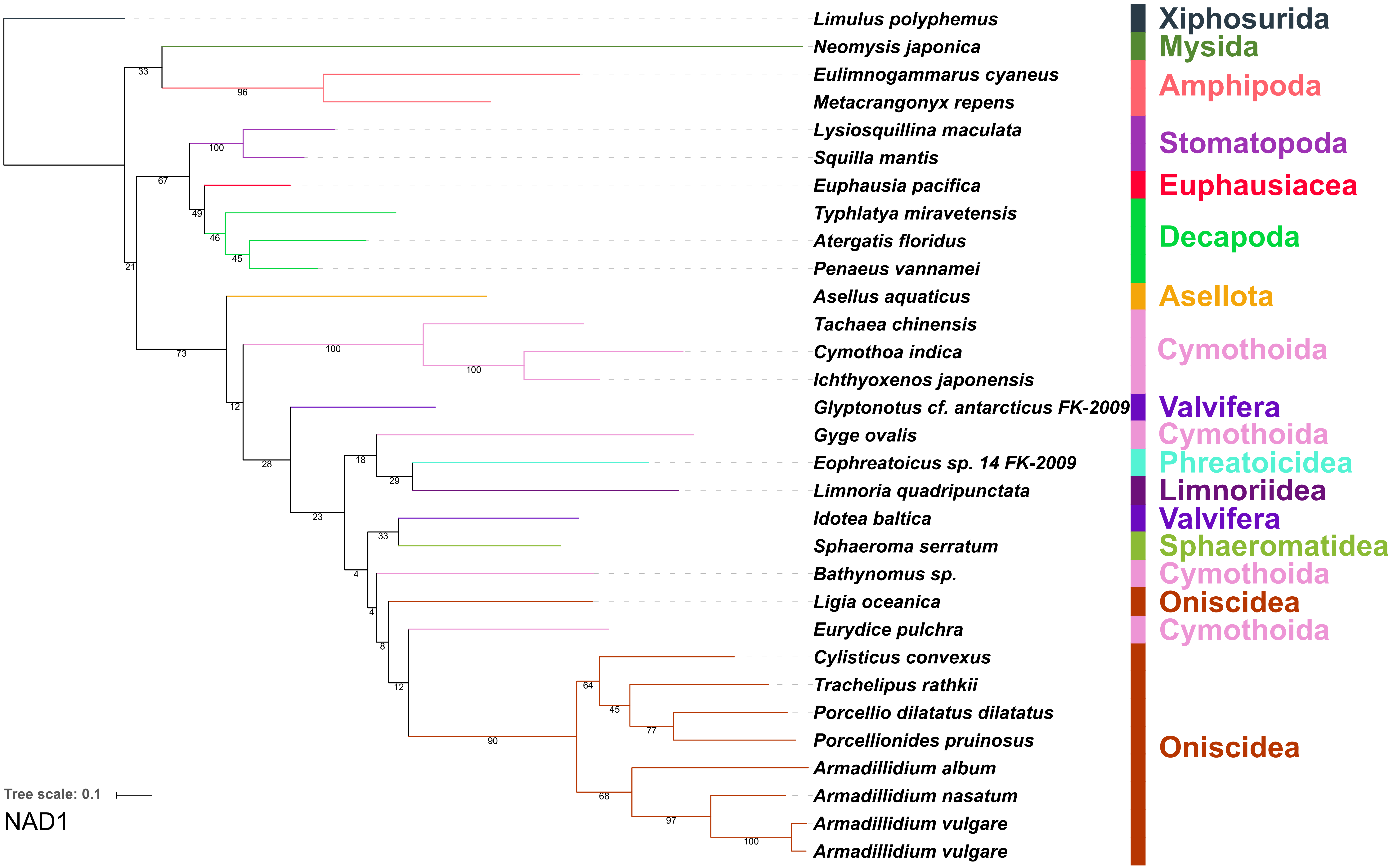
Limulus polyphemus
Neomysis japonica
Eulimnogammarus cyaneus
Metacrangonyx repens
Lysiosquillina maculata
Squilla mantis
Euphausia pacifica
Typhlatya miravetensis
Atergatis floridus
Penaeus vannamei
Asellus aquaticus
Tachaea chinensis
Cymothoa indica
Ichthyoxenos japonensis
Glyptonotus cf. antarcticus FK-2009
Gyge ovalis
Eophreatoicus sp. 14 FK-2009
Limnoria quadripunctata
Idotea baltica
Sphaeroma serratum
Bathynomus sp.
Ligia oceanica
Eurydice pulchra
Cylisticus convexus
Trachelipus rathkii
Porcellio dilatatus dilatatus
Porcellionides pruinosus
Armadillidium album
Armadillidium nasatum
Armadillidium vulgare
Armadillidium vulgare
Xiphosurida
Mysida
Amphipoda
Stomatopoda
Euphausiacea
Decapoda
Asellota
Cymothoida
Valvifera
Cymothoida
Phreatoicidea
Limnoriidea
Valvifera
Sphaeromatidea
Cymothoida
Oniscidea
Cymothoida
Oniscidea
Tree scale: 0.1
NAD1

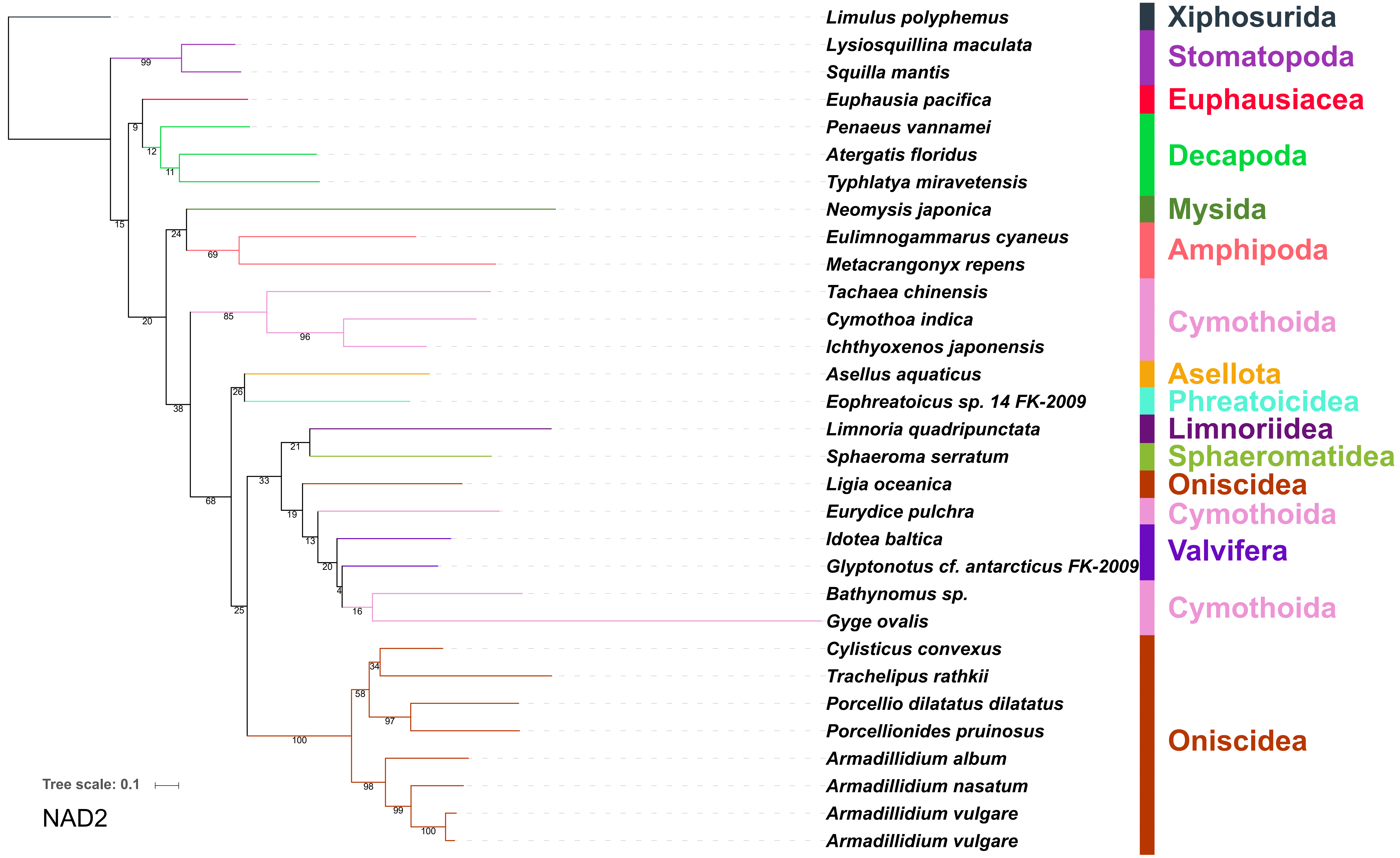

Limulus polyphemus
Lysiosquillina maculata
Squilla mantis
Euphausia pacifica
Penaeus vannamei
Atergatis floridus
Typhlatya miravetensis
Neomysis japonica
Eulimnogammarus cyaneus
Metacrangonyx repens
Tachaea chinensis
Cymothoa indica
Ichthyoxenos japonensis
Asellus aquaticus
Eophreatoicus sp. 14 FK-2009
Limnoria quadripunctata
Sphaeroma serratum
Ligia oceanica
Eurydice pulchra
Idotea baltica
Glyptonotus cf. antarcticus FK-2009
Bathynomus sp.
Gyge ovalis
Cylisticus convexus
Trachelipus rathkii
Porcellio dilatatus dilatatus
Porcellionides pruinosus
Armadillidium album
Armadillidium nasatum
Armadillidium vulgare
Armadillidium vulgare
Xiphosurida
Stomatopoda
Euphausiacea
Decapoda
Mysida
Amphipoda
Cymothoida
Asellota
Phreatoicidea
Limnoriidea
Sphaeromatidea
Oniscidea
Cymothoida
Valvifera
Cymothoida
Oniscidea
Tree scale: 0.1
NAD2

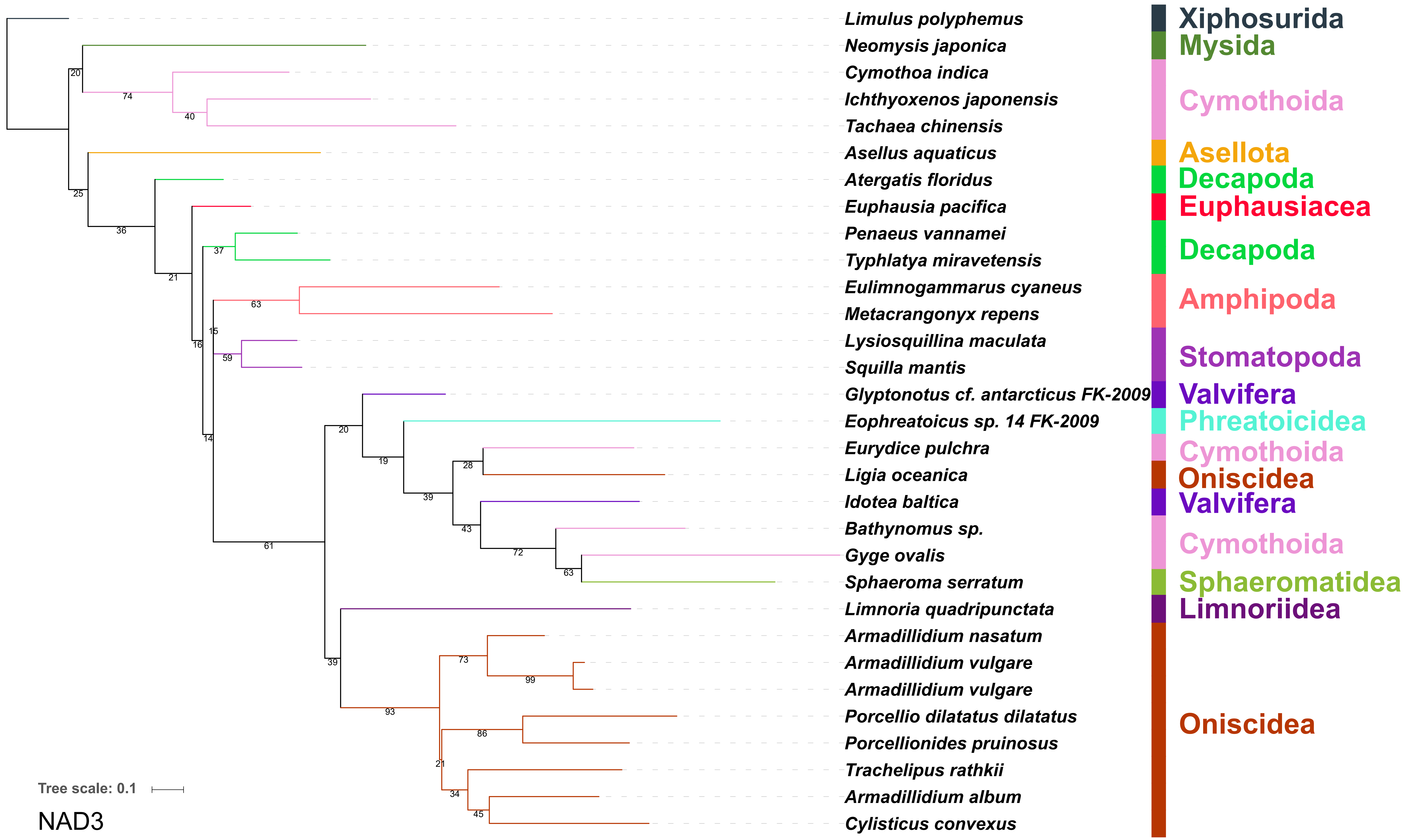
Limulus polyphemus
Neomysis japonica
Cymothoa indica
Ichthyoxenos japonensis
Tachaea chinensis
Asellus aquaticus
Atergatis floridus
Euphausia pacifica
Penaeus vannamei
Typhlatya miravetensis
Eulimnogammarus cyaneus
Metacrangonyx repens
Lysiosquillina maculata
Squilla mantis
Glyptonotus cf. antarcticus FK-2009
Eophreatoicus sp. 14 FK-2009
Eurydice pulchra
Ligia oceanica
Idotea baltica
Bathynomus sp.
Gyge ovalis
Sphaeroma serratum
Limnoria quadripunctata
Armadillidium nasatum
Armadillidium vulgare
Armadillidium vulgare
Porcellio dilatatus dilatatus
Porcellionides pruinosus
Trachelipus rathkii
Armadillidium album
Cylisticus convexus
Xiphosurida
Mysida
Cymothoida
Asellota
Decapoda
Euphausiacea
Decapoda
Amphipoda
Stomatopoda
Valvifera
Phreatoicidea
Cymothoida
Oniscidea
Valvifera
Cymothoida
Sphaeromatidea
Limnoriidea
Oniscidea
Tree scale: 0.1

NAD3

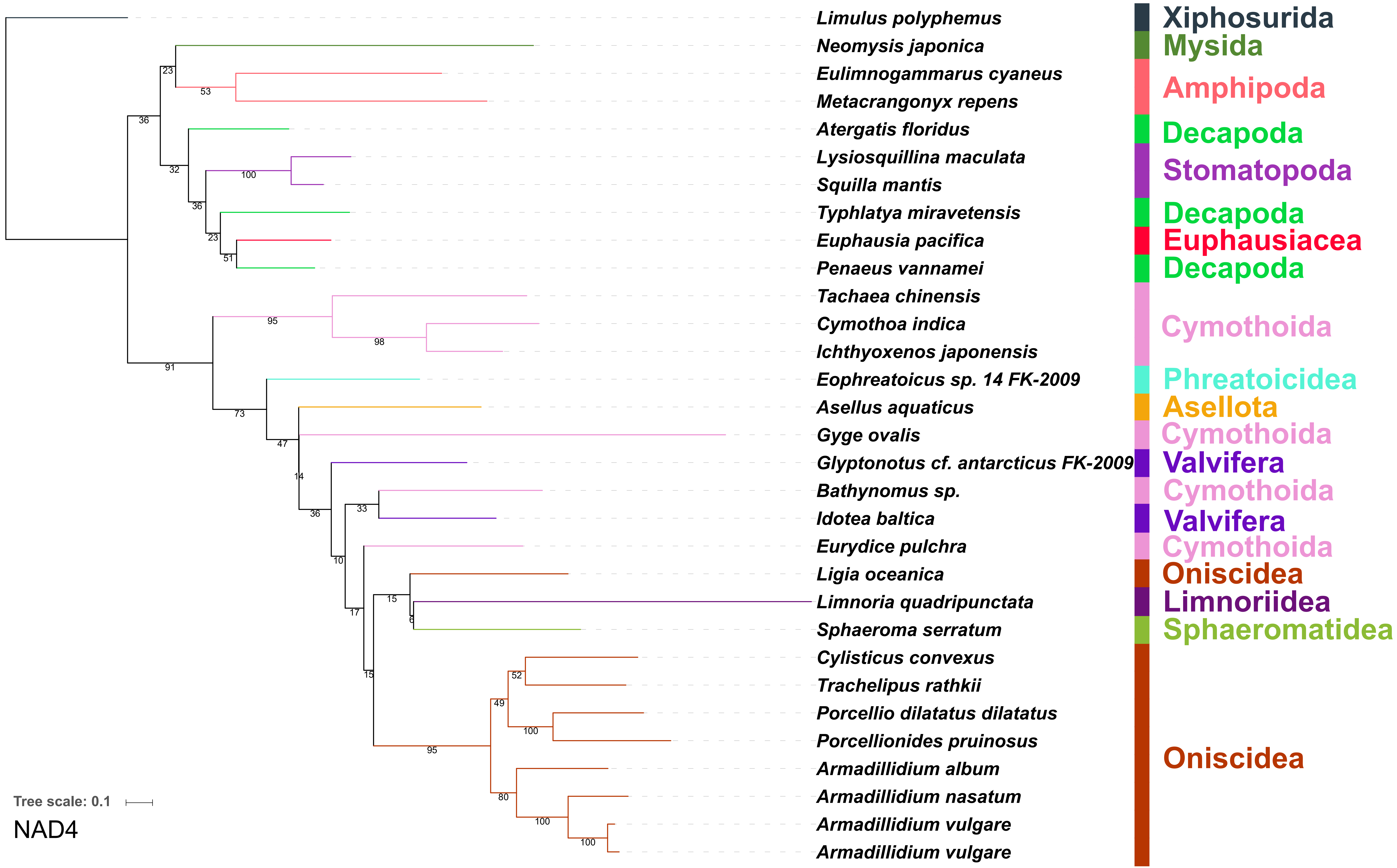
Limulus polyphemus
Neomysis japonica
Eulimnogammarus cyaneus
Metacrangonyx repens
Atergatis floridus
Lysiosquillina maculata
Squilla mantis
Typhlatya miravetensis
Euphausia pacifica
Penaeus vannamei
Tachaea chinensis
Cymothoa indica
Ichthyoxenos japonensis
Eophreatoicus sp. 14 FK-2009
Asellus aquaticus
Gyge ovalis
Glyptonotus cf. antarcticus FK-2009
Bathynomus sp.
Idotea baltica
Eurydice pulchra
Ligia oceanica
Limnoria quadripunctata
Sphaeroma serratum
Cylisticus convexus
Trachelipus rathkii
Porcellio dilatatus dilatatus
Porcellionides pruinosus
Armadillidium album
Armadillidium nasatum
Armadillidium vulgare
Armadillidium vulgare
Xiphosurida
Mysida
Amphipoda
Decapoda
Stomatopoda
Decapoda
Euphausiacea
Decapoda
Cymothoida
Phreatoicidea
Asellota
Cymothoida
Valvifera
Cymothoida
Valvifera
Cymothoida
Oniscidea
Limnoriidea
Sphaeromatidea
Oniscidea
Tree scale: 0.1
NAD4

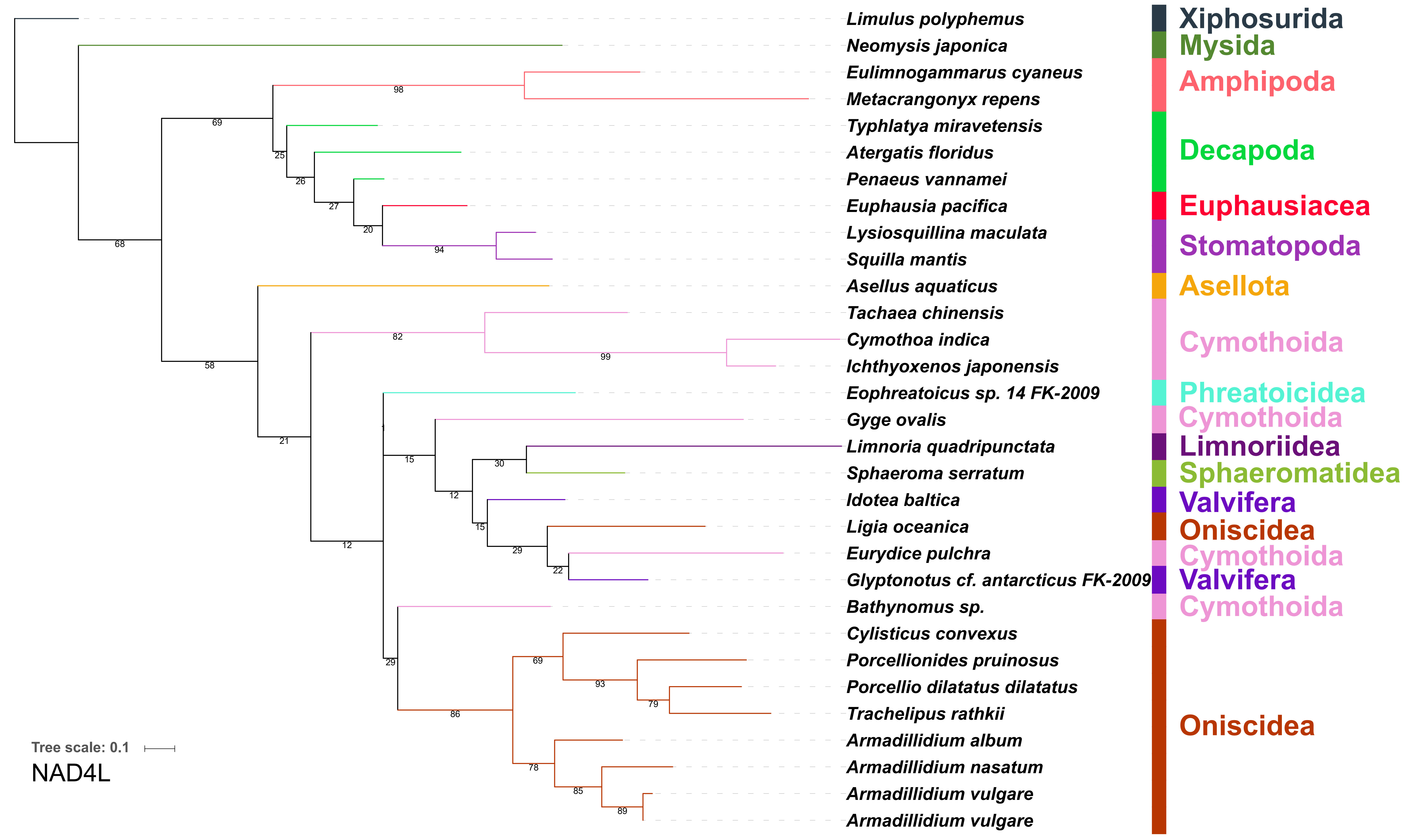
Limulus polyphemus
Neomysis japonica
Eulimnogammarus cyaneus
Metacrangonyx repens
Typhlatya miravetensis
Atergatis floridus
Penaeus vannamei
Euphausia pacifica
Lysiosquillina maculata
Squilla mantis
Asellus aquaticus
Tachaea chinensis
Cymothoa indica
Ichthyoxenos japonensis
Eophreatoicus sp. 14 FK-2009
Gyge ovalis
Limnoria quadripunctata
Sphaeroma serratum
Idotea baltica
Ligia oceanica
Eurydice pulchra
Glyptonotus cf. antarcticus FK-2009
Bathynomus sp.
Cylisticus convexus
Porcellionides pruinosus
Porcellio dilatatus dilatatus
Trachelipus rathkii
Armadillidium album
Armadillidium nasatum
Armadillidium vulgare
Armadillidium vulgare
Xiphosurida
Mysida
Amphipoda
Decapoda
Euphausiacea
Stomatopoda
Asellota
Cymothoida
Phreatoicidea
Cymothoida
Limnoriidea
Sphaeromatidea
Valvifera
Oniscidea
Cymothoida
Valvifera
Cymothoida
Oniscidea
Tree scale: 0.1
NAD4L

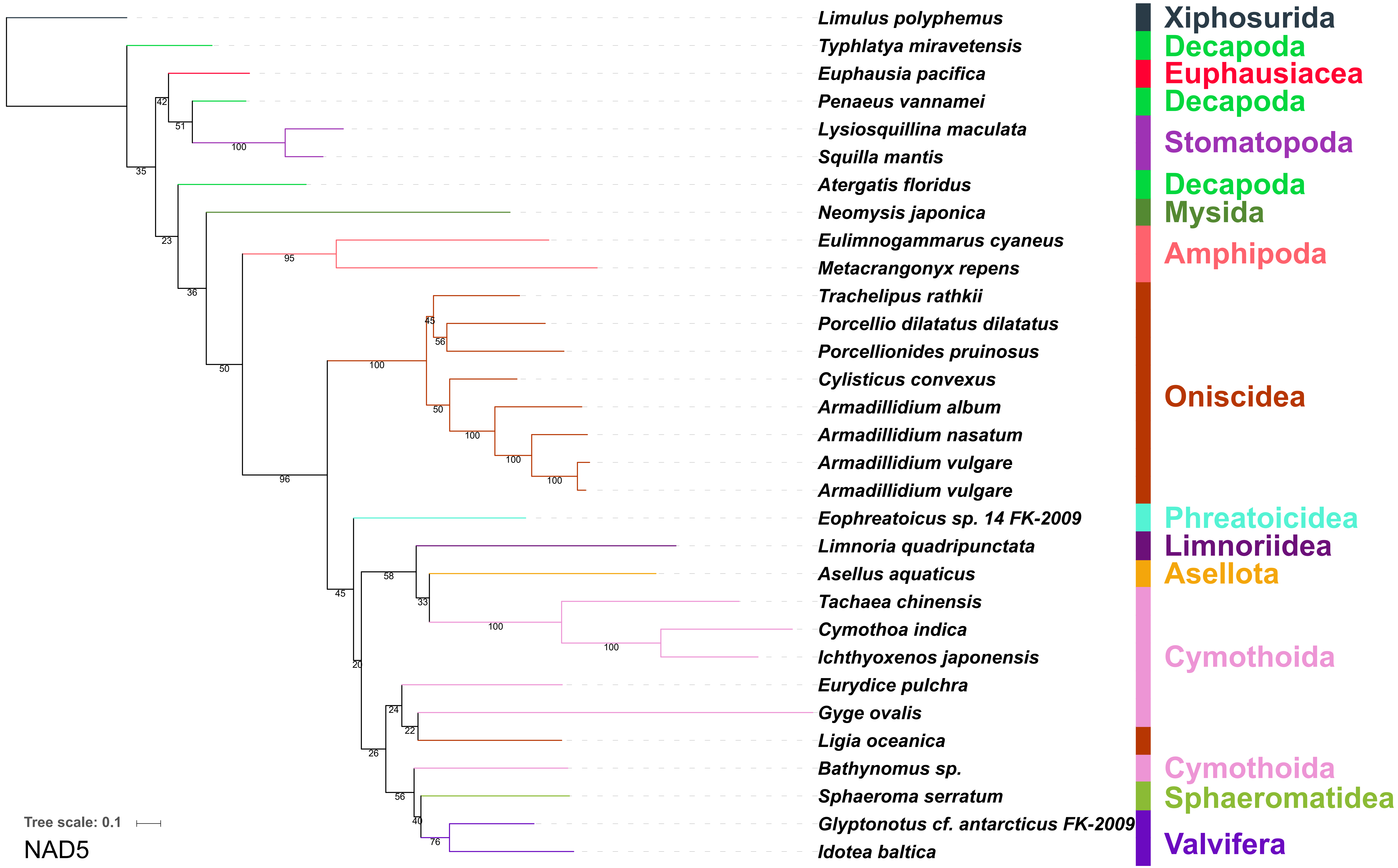

Limulus polyphemus
Typhlatya miravetensis
Euphausia pacifica
Penaeus vannamei
Lysiosquillina maculata
Squilla mantis
Atergatis floridus
Neomysis japonica
Eulimnogammarus cyaneus
Metacrangonyx repens
Trachelipus rathkii
Porcellio dilatatus dilatatus
Porcellionides pruinosus
Cylisticus convexus
Armadillidium album
Armadillidium nasatum
Armadillidium vulgare
Armadillidium vulgare
Eophreatoicus sp. 14 FK-2009
Limnoria quadripunctata
Asellus aquaticus
Tachaea chinensis
Cymothoa indica
Ichthyoxenos japonensis
Eurydice pulchra
Gyge ovalis
Ligia oceanica
Bathynomus sp.
Sphaeroma serratum
Glyptonotus cf. antarcticus FK-2009
Idotea baltica
Xiphosurida
Decapoda
Euphausiacea
Decapoda
Stomatopoda
Decapoda
Mysida
Amphipoda
Oniscidea
Phreatoicidea
Limnoriidea
Asellota
Cymothoida
Cymothoida
Sphaeromatidea
Valvifera
Tree scale: 0.1
NAD5

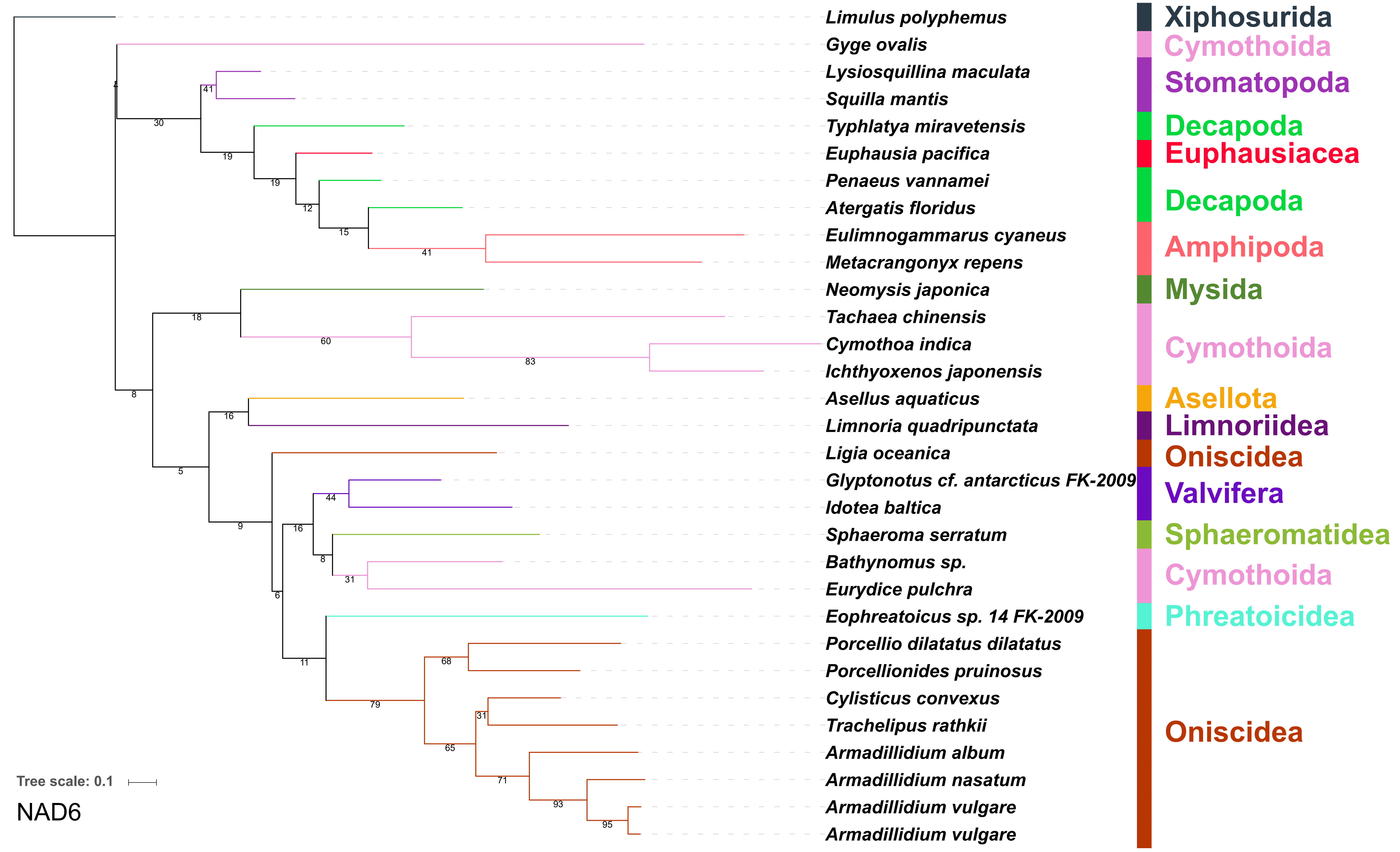

Limulus polyphemus
Gyge ovalis
Lysiosquillina maculata
Squilla mantis
Typhlatya miravetensis
Euphausia pacifica
Penaeus vannamei
Atergatis floridus
Eulimnogammarus cyaneus
Metacrangonyx repens
Neomysis japonica
Tachaea chinensis
Cymothoa indica
Ichthyoxenos japonensis
Asellus aquaticus
Limnoria quadripunctata
Ligia oceanica
Glyptonotus cf. antarcticus FK-2009
Idotea baltica
Sphaeroma serratum
Bathynomus sp.
Eurydice pulchra
Eophreatoicus sp. 14 FK-2009
Porcellio dilatatus dilatatus
Porcellionides pruinosus
Cylisticus convexus
Trachelipus rathkii
Armadillidium album
Armadillidium nasatum
Armadillidium vulgare
Armadillidium vulgare
Xiphosurida
Cymothoida
Stomatopoda
Decapoda
Euphausiacea
Decapoda
Amphipoda
Mysida
Cymothoida
Asellota
Limnoriidea
Oniscidea
Valvifera
Sphaeromatidea
Cymothoida
Phreatoicidea
Oniscidea
Tree scale: 0.1
NAD6

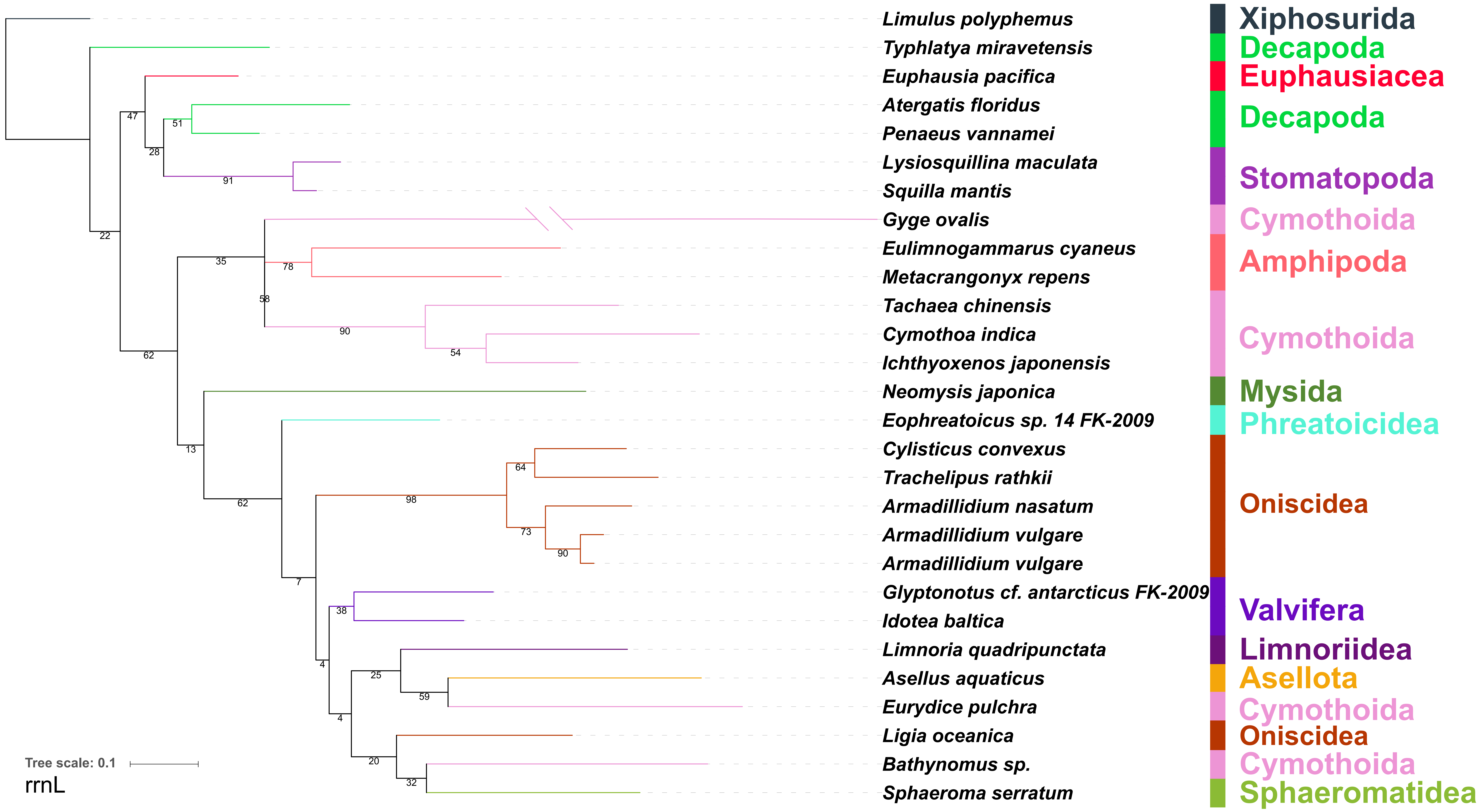
Limulus polyphemus
Typhlatya miravetensis
Euphausia pacifica
Atergatis floridus
Penaeus vannamei
Lysiosquillina maculata
Squilla mantis
Gyge ovalis
Eulimnogammarus cyaneus
Metacrangonyx repens
Tachaea chinensis
Cymothoa indica
Ichthyoxenos japonensis
Neomysis japonica
Eophreatoicus sp. 14 FK-2009
Cylisticus convexus
Trachelipus rathkii
Armadillidium nasatum
Armadillidium vulgare
Armadillidium vulgare
Glyptonotus cf. antarcticus FK-2009
Idotea baltica
Limnoria quadripunctata
Asellus aquaticus
Eurydice pulchra
Ligia oceanica
Bathynomus sp.
Sphaeroma serratum
Xiphosurida
Decapoda
Euphausiacea
Decapoda
Stomatopoda
Cymothoida
Amphipoda
Cymothoida
Mysida
Phreatoicidea
Oniscidea
Valvifera
Limnoriidea
Asellota
Cymothoida
Oniscidea
Cymothoida
Sphaeromatidea
47
51
28
91
22
35
78
58
90
54
62
13
62
64
98
73
90
7
38
4
25
59
4
20
32
Tree scale: 0.1
rrnL

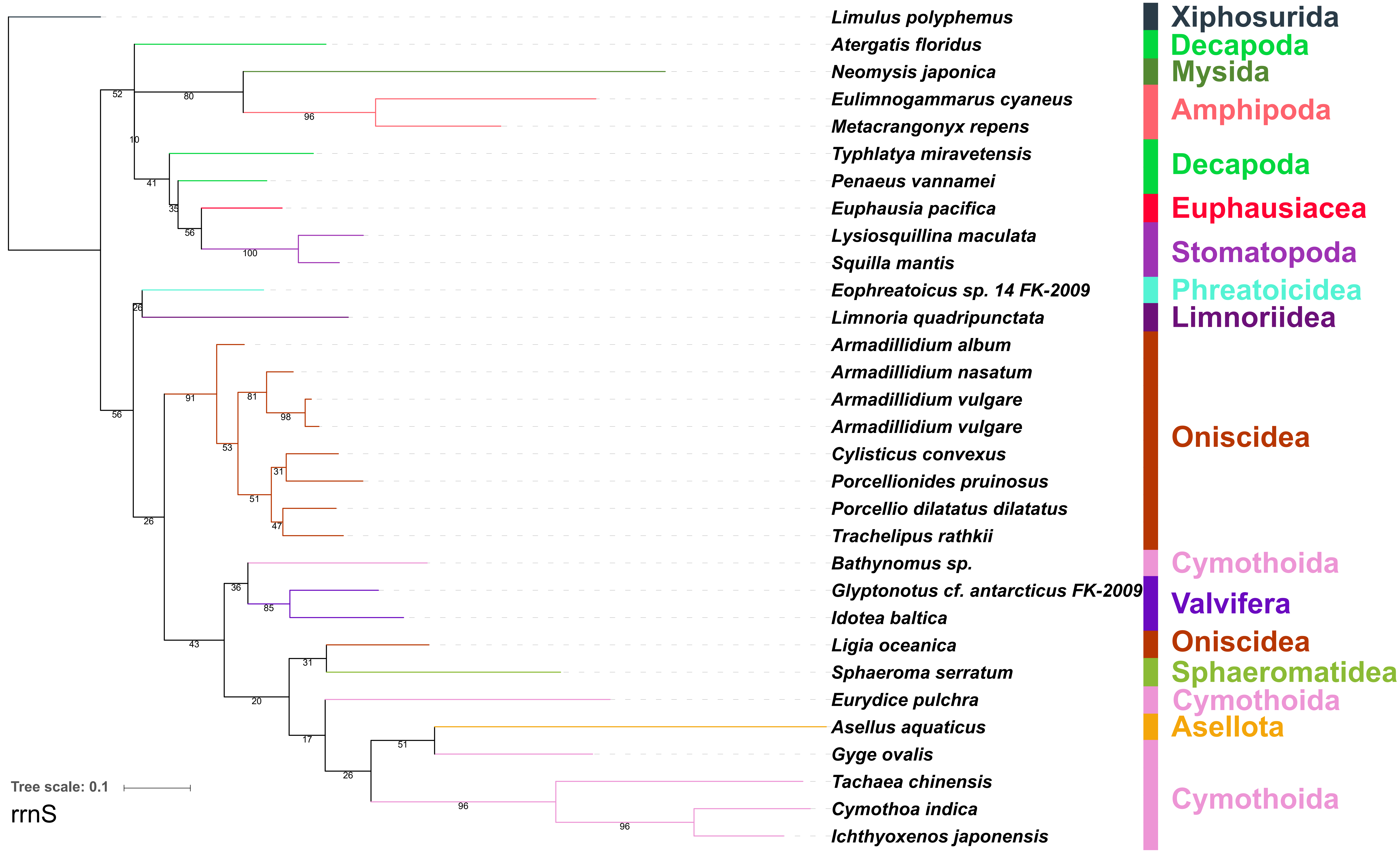
Limulus polyphemus
Atergatis floridus
Neomysis japonica
Eulimnogammarus cyaneus
Metacrangonyx repens
Typhlatya miravetensis
Penaeus vannamei
Euphausia pacifica
Lysiosquillina maculata
Squilla mantis
Eophreatoicus sp. 14 FK-2009
Limnoria quadripunctata
Armadillidium album
Armadillidium nasatum
Armadillidium vulgare
Armadillidium vulgare
Cylisticus convexus
Porcellionides pruinosus
Porcellio dilatatus dilatatus
Trachelipus rathkii
Bathynomus sp.
Glyptonotus cf. antarcticus FK-2009
Idotea baltica
Ligia oceanica
Sphaeroma serratum
Eurydice pulchra
Asellus aquaticus
Gyge ovalis
Tachaea chinensis
Cymothoa indica
Ichthyoxenos japonensis
Xiphosurida
Decapoda
Mysida
Amphipoda
Decapoda
Euphausiacea
Stomatopoda
Phreatoicidea
Limnoriidea
Oniscidea
Cymothoida
Valvifera
Oniscidea
Sphaeromatidea
Cymothoida
Asellota
Cymothoida
Tree scale: 0.1
rrnS
